# Harness engineering enables a multi-agent LLM system to generate cross-domain hypotheses beyond single-investigator reach

**DOI:** 10.64898/2026.10.25.761203

**Authors:** Keiro Watanabe, Takeshi Omasa

## Abstract

Large language models (LLMs) are increasingly used to generate scientific hypotheses, and a more capable model is widely assumed to drive output quality. We tested this with a biology-focused multi-agent system coupling an LLM to five Model Context Protocol servers exposing 66 external tools. Upgrading the Synthesizer and Critic from Claude Sonnet 4.6 to Opus 4.8 initially appeared to improve quality, count, and reliability. Under a fully matched harness (identical prompts, schema, hypothesis count, and queries), the differences vanished: scores were indistinguishable on an iPSC-reprogramming task and favoured Sonnet 4.6 on a CHO cell-line-engineering task (18.4 vs 17.4; Welch t = 2.89, p < 0.05). The dominant, controllable driver of quality is therefore the prompting harness, not the model tier. We then introduce the ABCD-Stack (contract-gated prompting, diversity-first generation, adversarial refinement, and cross-family evaluation), which builds these properties in as architecture and shifts outputs toward explicitly falsifiable, causally framed hypotheses beyond the reach of a single-domain investigator. The hypotheses are falsifiable by construction but not yet experimentally validated.

## 1. Introduction

The generation of novel scientific hypotheses has traditionally depended on the accumulated expertise of individual researchers who manually integrate knowledge across literature, structural biology, biochemical networks, and medicinal chemistry. This process is inherently slow, domain-siloed, and subject to cognitive biases that favor incremental rather than cross-disciplinary innovation. The rapid expansion of biomedical literature — exceeding 1.5 million PubMed publications annually — combined with the proliferation of large-scale biological databases (protein interaction networks, structural repositories, clinical trial registries) has made comprehensive evidence synthesis increasingly intractable for human researchers alone.

Recent advances in large language models (LLMs) have demonstrated remarkable capability for scientific reasoning, literature comprehension, and structured knowledge integration [1–3,5,6,9,10]. In parallel, retrieval-augmented generation (RAG) frameworks have enabled LLMs to draw their outputs from curated, domain-specific knowledge bases via RAG, rather than relying solely on the parameters learned during pre-training [7]. Yet applying these capabilities to the hardest stage of the research cycle — the generation of genuinely novel, mechanistically grounded, experimentally falsifiable hypotheses — remains largely unexplored [4,31,32].

A key architectural limitation of existing scientific AI assistants is that they typically operate within a single knowledge domain. A chemistry-oriented model may lack access to protein interaction network topology; a literature review agent may be unaware of structural biology data that would critically constrain or redirect a hypothesis. Meaningful scientific breakthroughs frequently arise from the intersection of multiple fields — for example, the recognition that the clinical activity of APR-246 (PRIMA-1Met), a compound developed to restore the tumour-suppressor function of p53 [34,35], may involve redox-related mechanisms beyond p53-specific refolding [36,39,43,44], a cross-domain insight that simultaneously reframes mechanism-of-action understanding and opens new therapeutic directions. Such connections are precisely the kind that a single-domain agent is unlikely to surface.

The Model Context Protocol (MCP), introduced as a standardized tool-use interface for LLM agents [10], provides a practical mechanism to address this limitation. Through MCP, an LLM can invoke specialized, externally hosted tools via a structured request/response interface, effectively extending the model’s capabilities to arbitrary computational backends without retraining. This architecture separates scientific reasoning (handled by the LLM) from data access and computation (delegated to MCP servers), and, by combining the cross-domain material that serves as the substrate of reasoning, enables the construction of a heterogeneous, domain-specific tool ecosystem composable at inference time.

LangGraph [12], a graph-based multi-agent orchestration framework built on LangChain [11] and grounded in the Pregel model [13], defines agent workflows as directed graphs with conditional routing, shared-state reducers, and bounded tool-use loops. This programmable, stateful architecture suits the structured, multi-stage nature of scientific hypothesis generation, in which specialized agents sequentially collect evidence and an integrating node consolidates their outputs.

Here we report the design, implementation, and evaluation of a multi-agent system for automated novel scientific hypothesis generation, built on five MCP servers collectively exposing 66 domain-specific tools. The system implements a six-node LangGraph pipeline in which four specialist evidence-collection agents (Literature Scout, Network Analyst, Structure Function, Chemistry Opportunity) sequentially query literature databases [22], protein interaction networks [18,19], AlphaFold structural predictions [15–17], and medicinal chemistry repositories [21,29,30]. Their outputs are integrated by a Hypothesis Synthesizer, which generates 2–5 cross-domain hypotheses in structured JSON(JavaScript Object Notation) format, and a Hypothesis Critic, which independently scores each hypothesis on novelty, testability, impact, and — in enhanced experiments — evidence consistency.

A recurring concern with automated hypothesis generation is that fluent, well-cited text does not by itself constitute a valid scientific hypothesis: an LLM’s output can sometimes present a mechanistically plausible narrative that is unfalsifiable, already known, or even contradicted by the very evidence cited in its support [31,32]. The value of a generated hypothesis is therefore inseparable from whether it can be tested by an experiment that could refute it, and whether an impartial evaluation separated from the generator’s inductive bias independently assesses its novelty and validity. We therefore treat experimental falsifiability and independent, impartial evaluation as primary design goals rather than downstream conveniences. Each hypothesis is required to carry an explicit Popperian falsification condition and a calibration-neutral quantitative prediction.

We evaluated the system across three research questions spanning oncology drug discovery (TP53-mutant solid tumors, KRAS G12C pancreatic cancer) and biopharmaceutical manufacturing (CHO cell line engineering), using two Claude model generations, Sonnet 4.6 and Opus 4.8 with Adaptive Thinking. An initial comparison appeared to show that the Opus configuration produced more structured, more numerous, and more reliable hypotheses. That apparent advantage, however, did not survive a controlled re-test: once the two configurations were matched on prompt, output schema, hypothesis count, and query coverage, the model differences vanished, and the gains traced instead to the prompting harness. The central result of this paper is therefore methodological — in a multi-agent discovery pipeline, output quality, count, and reliability are governed by how the task is framed and contracted, not by the LLM tier. We turn that negative result into a constructive one by building an enhanced pipeline, the ABCD-Stack, that makes the decisive harness properties architectural, and we show that it generates diverse, contract-checked, independently evaluated cross-domain hypotheses while surfacing its own coverage and agreement limits. Throughout, we treat experimental falsifiability and independent evaluability not as afterthoughts but as primary design objectives, and frame future validatability in terms of two empirical programs — wet-laboratory testing and blinded expert benchmarking — and discuss the AI-generated hypotheses accordingly.

A final scope note frames what the system does and does not establish. The pipeline automates hypothesis quality assurance in a specific, bounded sense: an LLM Critic scores each hypothesis for novelty, testability, impact, consistency, and — in the enhanced architecture — faithfulness to the brief, and the harness enforces, for every output, an explicit falsification condition and a criteria-based quantitative prediction. Rather than certifying that a hypothesis is correct, these are properties of a future-verifiable, well-formed scientific hypothesis. Whether any generated hypothesis is actually true, or scientifically valuable, cannot be settled by the system itself; that judgement rests with domain experts and, ultimately, with wet-laboratory experiment. A central benefit is biological rather than merely computational: by assigning integrative cross-domain reasoning and structure/chemistry-grounded analysis to parallel agents that share the same evidence, the system behaves like a team of same-rigour specialists and reaches cross-domain hypotheses that a single investigator, deep in one domain, is unlikely to formulate alone.

## 2. Materials and Methods

### 2.1 Computational Environment

All experiments were conducted inside an LXC container (Ubuntu 24.04 LTS, hostname: *mylangchain-container*) running Python 3.13 via a Conda environment on the local host. Service orchestration was managed with Docker Compose [67], providing network isolation and reproducible port mappings (:3001–:3005 for MCP endpoints, :8888 for JupyterLab, :7474/:7687 for Neo4j). Jupyter notebooks [68] were used as the primary experimental interface.

### 2.2 Large Language Models

Two Claude model generations were evaluated across sequential experimental phases (Table 1): Text embeddings for the Aurora pgvector database were generated with Amazon Titan Embed Text v2 [25] (dimension = 1,024). All LLM calls were made via the LangChain AWS Bedrock integration (`ChatBedrock`) [11] and, for Adaptive Thinking inference [78] (Anthropic’s inference-time extended-reasoning mode, in which the model is given a token budget to plan and self-check on an internal scratchpad before responding), the native `AnthropicBedrockMantle` Python client with `thinking={“type”: “adaptive”}` and `max_tokens=8000`.

**Table 1.**
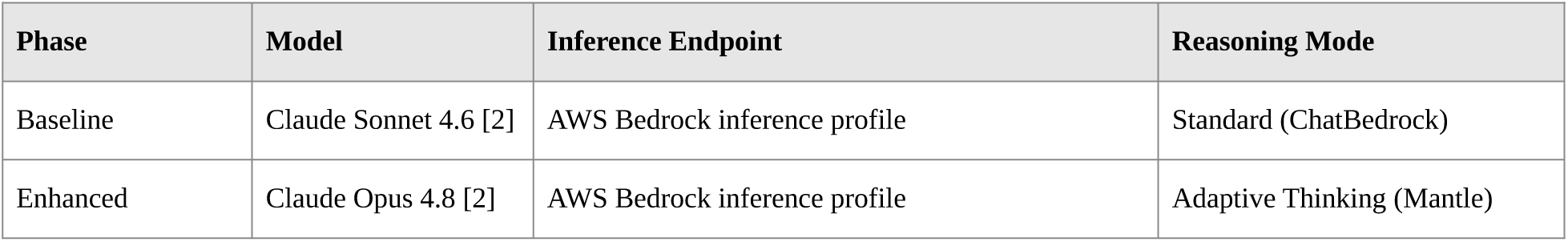
Model configurations for the baseline and enhanced pipelines, with inference endpoint and reasoning mode.

### 2.3 MCP Server Infrastructure

Five Model Context Protocol servers [10] were deployed as Docker containers within the LXC environment, collectively exposing **66 tools** accessible via SSE (Server-Sent Events) transport (Figure 3). Each server was assigned a dedicated role and tool subset (Table 2): Across all five servers the same access pattern holds: each server is implemented with FastMCP and served over SSE on its assigned port, and the Claude LLM — not a fixed query — is the agent that selects among a server’s tools from within the relevant node’s ReAct tool-use loop [8] (via llm.bind_tools() over MCP), formulating its own arguments and thresholds per call. The per-server descriptions below therefore summarize each backend’s role and tool groups; the complete per-tool inventory for all 66 tools is given in Figure 5, and the verbatim tool schemas are not repeated in the text.

**Table 2.** The five MCP tool servers: port, backend, tool count, and primary function (66 tools total). The full per-tool inventory for all 66 tools is shown in Figure 5.

| Server | Port | Backend | Tools | Primary Function |
| --- | --- | --- | --- | --- |
| mcp-chem-calc | :3001 | RDKit [29]/PubChem [21] (local), Rowan QM/ADMET [30] (cloud, optional) | 15 | Molecular property calculation, ADMET prediction, quantum chemistry |
| mcp-biochem-calc | :3002 | BioPython [28] (local), BioMCP [27] CLI (federated DBs) | 16 | Protein/DNA analysis, UniProt [20], clinical variant/drug annotation |
| mcp-paper-search | :3003 | Aurora pgvector [24], Semantic Scholar [22], NCBI E-utilities, paper-search-mcp [73] | 17 | Literature search, paper indexing, full-text retrieval |
| mcp-bio-graphrag | :3004 | Neo4j GraphRAG (STRING [18] + CTD [19], APOC) | 7 | Protein interaction network queries, disease–gene association |
| mcp-alphafold | :3005 | AlphaFold EBI public API [15] | 11 | Structure retrieval, per-residue pLDDT analysis, batch queries, organism coverage, viewer export |

#### 2.3.1 Biological Knowledge Graph

The mcp-bio-graphrag server was backed by a local Neo4j instance populated with two public datasets:

- **STRING DB v12.0** [18] (CC BY 4.0): 19,699 human protein nodes, 473,860 INTERACTS_WITH edges (combined_score ≥ 700, high-confidence interactions only).
- **CTD** [19] (Comparative Toxicogenomics Database; freely available to non-commercial users under the CTD terms of use): 5,735 disease nodes, 31,638 ASSOCIATED_WITH edges linking genes to diseases.

Following the graph-RAG approach of Edge et al. [26], the server executes parameterized Cypher queries and returns compact, structured subgraphs rather than raw text, exposing seven tools in two phases. Four Phase 1 tools operate over the STRING interaction layer — retrieving a gene’s interaction neighbourhood (score-filtered and edge-capped so a hub cannot flood the context), ranking hub proteins by degree, tracing shortest interaction paths, and full-text-searching functional annotations. Three Phase 2 tools add the CTD disease dimension; the central cross_query_disease_network chains both datasets in one query — selecting disease-associated seed genes above a GDA threshold, then expanding each seed’s STRING neighbourhood — answering cross-database questions (e.g. “interaction hubs of Alzheimer-associated proteins”) that neither dataset resolves alone.

#### 2.3.2 Semantic Paper Search Backend

The mcp-paper-search server combines a persistent semantic index with live external retrieval, exposing seventeen tools in four groups: two vector-store tools over an Amazon Aurora PostgreSQL/pgvector store [23,24] (semantic search and DOI ingestion), five Semantic Scholar Graph-API tools [22], a PubMed E-utilities tool, and nine tools wrapping the open-source paper-search-mcp package [73] (arXiv, PubMed, bioRxiv, medRxiv, Google Scholar), whose heterogeneous outputs are normalized to a common schema. Together these form an embed-then-retrieve pipeline: index_paper_by_doi resolves a DOI, deduplicates by a DOI-derived UUID, and embeds the abstract with Amazon Titan Embed Text v2 [25] into the pgvector store, while later queries embed with the same model and return nearest abstracts by cosine similarity. External connectors supply breadth and freshness on first encounter; the pgvector index supplies fast meaning-based recall of anything indexed before — the path verified end to end in Section 3.1.1.

#### 2.3.3 Structure Prediction Backend

The mcp-alphafold server exposes eleven tools over the AlphaFold EBI public API [15], grouped into structure retrieval, confidence analysis, and scale/context support. Beyond fetching structures and metadata, two confidence-analysis tools compute per-residue and per-region pLDDT summaries so the agent can judge which parts of a predicted structure are reliable enough to reason over (e.g. a well-folded pocket versus a disordered loop); the remaining tools add batch queries, organism coverage, and viewer/export helpers. The Structure-Function node selects among these per call, so structural confidence is treated as evidence rather than assumed.

#### 2.3.4 Biochemical Computation Backend

The mcp-biochem-calc server exposes sixteen tools in two groups. Four local BioPython [28] tools perform sequence-level protein/DNA analysis (composition, molecular weight, translation, and related properties); the remaining twelve wrap the BioMCP command-line interface [27] to reach federated databases for UniProt [20] annotation and clinical variant/drug information. This two-group design lets one server answer both offline sequence questions and live database lookups within the same tool-use loop.

#### 2.3.5 Chemical Computation Backend

The mcp-chem-calc server exposes fifteen tools in a tiered design. Three local RDKit [29]/PubChem [21] tools give immediate, deterministic molecular-property and descriptor calculation; the remaining twelve wrap the optional Rowan cloud platform [30] for quantum-chemistry and ADMET prediction. The tiering gives the Chemistry-Opportunity node both fast local computation and, when warranted, higher-fidelity cloud calculations, chosen per call.

### 2.4 Multi-Agent System Architecture

#### 2.4.1 Baseline Research Agent

A parallel multi-agent system was implemented using LangGraph’s `StateGraph` API [12]. Three specialist agents operated concurrently (Literature, Bioinformatics, Chemistry) and integrated by a Coordinator node. Each agent executed a bounded tool-use loop (max 12 iterations) using `llm.bind_tools()`, following the ReAct paradigm [8].

#### 2.4.2 Novel Hypothesis Generation Agent

A sequential six-node pipeline was designed for cross-domain hypothesis generation (Figure 1):

**Figure 1.**
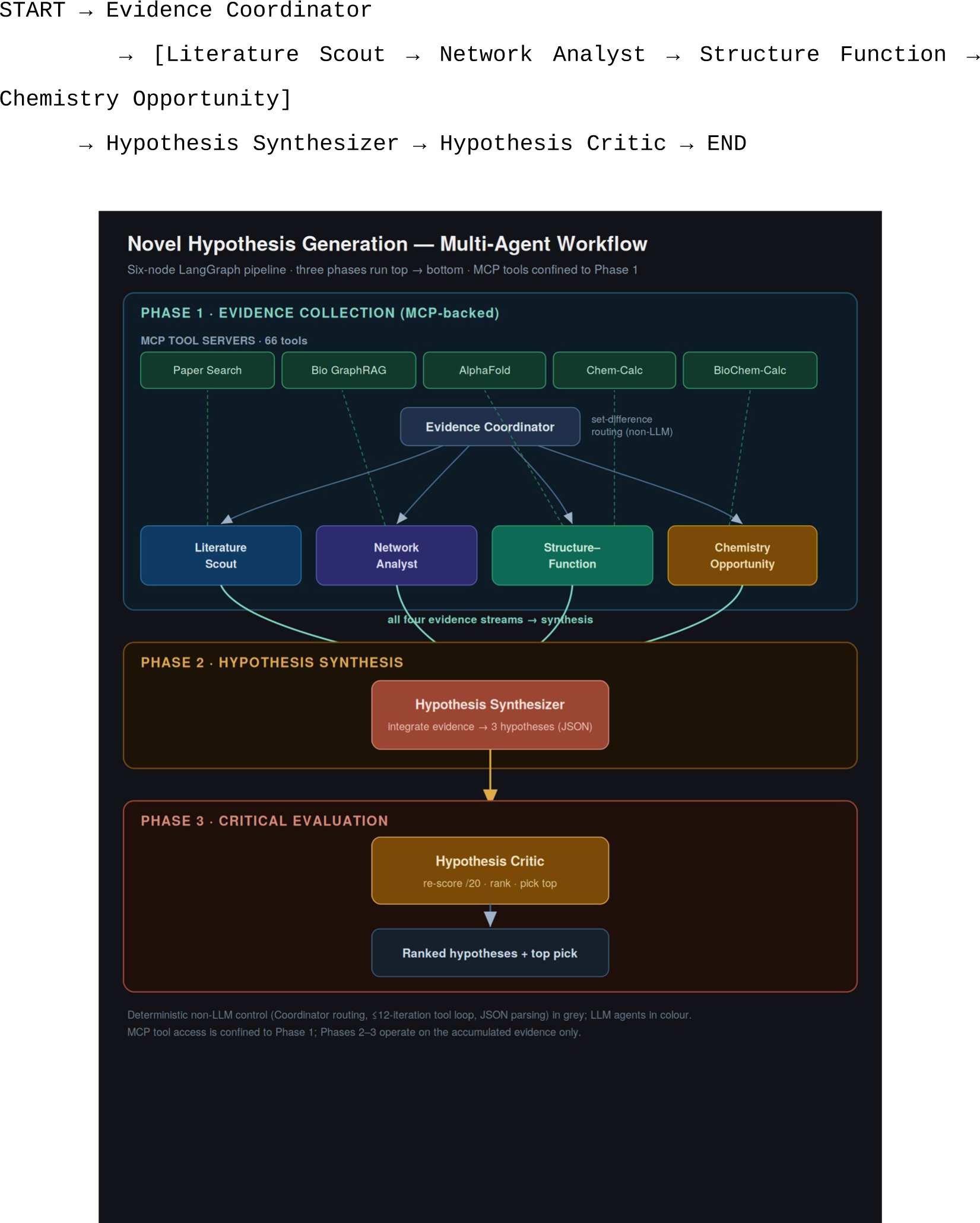
Novel hypothesis generation — multi-agent workflow. Six-node LangGraph pipeline. Phase 1 (Evidence Collection): the Evidence Coordinator sequentially dispatches Literature Scout, Network Analyst, Structure Function, and Chemistry Opportunity, each using an exclusive, non-overlapping tool subset accessed over SSE (:3001–:3005, 66 tools total). Phase 2 (Hypothesis Synthesis): the Hypothesis Synthesizer (Claude Sonnet 4.6, or Claude Opus 4.8 + Adaptive Thinking in the enhanced configuration) integrates all evidence into three structured-JSON hypotheses with explicit falsification conditions, quantitative predictions, and differentiation from prior art. Phase 3 (Critical Evaluation): the Hypothesis Critic independently scores novelty, testability, impact, and consistency (each /5, max 20) and designates the top-ranked hypothesis.

**Phase 1 — Evidence Collection.** The Evidence Coordinator dispatched four specialist sub-agents in fixed sequential order. Each sub-agent used an exclusive, non-overlapping tool subset and executed up to 12 tool-call iterations per activation.

**Phase 2 — Hypothesis Synthesis.** The Hypothesis Synthesizer received concatenated evidence from all four agents and produced 2–3 hypotheses per query in strict JSON format. Required fields: `title`, `statement` (if…then form), `mechanism`, `evidence_basis`, `falsification` (Popperian refutation condition), `differentiation` (departure from prior art), `quantitative_prediction`, `experiment`, and three self-scores (`novelty_score`, `testability_score`, `impact_score`, each /5). In the enhanced Opus 4.8 version, Adaptive Thinking inference was used for this node.

**Phase 3 — Critical Evaluation.** The Hypothesis Critic independently re-scored each hypothesis, adding `strength`, `weakness`, `devil_advocate`, `total_score`, and designating the top-ranked hypothesis via `top_hypothesis_id`. The Hypothesis Critic node ran in both model configurations: in the Sonnet 4.6 baseline it used the LangChain `ChatBedrock` client, and in the enhanced configuration it used the Bedrock Mantle client with Adaptive Thinking (`max_tokens=6000`).

#### 2.4.3 Evaluation Scoring Schema

Each generated hypothesis is scored on an explicit rubric rather than by holistic judgment. In the standard four-criterion schema the Critic assigns an integer from 1 to 5 on each of novelty, testability, impact, and consistency, summing to a maximum of 20; this schema is applied identically to every Sonnet 4.6 and Opus 4.8 run once the harness is unified. Only the ABCD-Stack (Section 2.4.4) adds a fifth criterion, faithfulness to the brief, raising its maximum to 25; every other experiment in this paper uses the four-criterion, max-20 schema. In other words, 25-point totals appear only for ABCD-Stack runs (Section 3.6), never for the standard Sonnet 4.6 or Opus 4.8 runs. The criteria are defined in Table 3, where the fifth criterion is flagged as ABCD-Stack-only.

**Table 3.** Evaluation scoring schema. Criteria 1–4 constitute the four-criterion schema (maximum 20) used throughout Sections 3.3–3.5 and applied identically to both model tiers once the harness is unified. Criterion 5 (faithfulness to the brief) is applied only in the ABCD-Stack (Section 3.6), which uses this five-criterion, maximum-25 schema; every other experiment in this paper — all standard Sonnet 4.6 and Opus 4.8 runs of Sections 3.3–3.5 — uses the four-criterion, maximum-20 schema. This was confirmed against the implemented critics: the standard Critic scores only the first four criteria, and the fifth is added solely by the ABCD-Stack cross-family Critic (verbatim prompts in Supplementary Information S4). Standard scores are self-assigned by a Critic of the same model family that generated the hypothesis; cross-family scoring is examined in Section 3.6.2.

| Criterion | Max | Definition (score 5 → score 1) |
| --- | --- | --- |
| Novelty | /5 | 5 = fully novel (no prior explicit report); 1 = known mechanism rephrased |
| Testability | /5 | 5 = immediately verifiable with current technology; 1 = currently impossible |
| Impact | /5 | 5 = paradigm-shifting; 1 = limited domain significance |
| Consistency | /5 | 5 = fully consistent with all four evidence agents; 1 = contradicts evidence |
| Faithfulness to brief | /5 | ABCD-Stack only (Section 2.4.4): 5 = directly answers the brief as posed; 1 = fluent but off-brief |

#### 2.4.4 ABCD-Stack Enhanced Pipeline

The ABCD-Stack is an enhanced pipeline that keeps the baseline workflow (Figure 1) and its four MCP-backed evidence agents unchanged, and strengthens the harness by adding deterministic, non-LLM control stages at the points where the baseline was weakest. It adds four architectural pillars — A-CGP (contract-gated prompting), B-DFG (diversity-first generation), C-ARL (adversarial refinement loop), and D-CFEC (cross-family evaluation) — realized as pipeline phases: a contract-gated Decomposer (Phase 0) that expands the brief into five to eight measurable sub-questions; three parallel persona Synthesizers (Phase 2) with a deterministic schema gate (Phase 2.5) and a keyword-coverage audit (Phase 2.6) against those sub-questions; a greedy Jaccard de-duplication selector (Phase 4); an adversarial Red-Team/Reviser loop (Phase 3, C-ARL); and cross-family scoring by two independent model families with a per-run agreement statistic (Phase 5, D-CFEC). Each stage is a prompt or a deterministic rule executed by the same model, so the added quality is a property of the harness, not the model tier.

The rationale for each phase — which diagnosed harness weakness it closes — and the full per-phase mechanics are given in Section 3.6.1, and are not repeated here. Implementation note. All ABCD-Stack LLM calls used AWS Bedrock, with adaptive thinking on the Opus 4.8 Decomposer, the persona synthesizers, and the critics; the scoring rubric adds the fifth criterion, faithfulness to the brief, for a maximum of 25 points (Table 3). Each of the two condition-matched case studies (CHO, iPSC) was run five times independently, producing the ten reports analyzed in Section 3.6.

#### 2.4.5 Prompt-Harness Implementation

Concretely, these prompts occupy the four harness layers of Figure 2. (i) The seven node prompts are the per-node system prompts of the pipeline in Figure 1: the Evidence Coordinator (a deterministic routing prompt that cycles the four evidence agents in a fixed order and then hands off to synthesis), the four evidence-collection agents (Literature Scout, Network Analyst, Structure-Function, and Chemistry Opportunity, each with a role definition, numbered investigation lenses, and a prescribed output format), the Hypothesis Synthesizer, and the Hypothesis Critic. They constitute the per-node system-prompt layer of Figure 2. (ii) The Synthesizer and Critic quality contracts are the output-contract layer of Figure 2: the Synthesizer contract enumerates the five mandatory conditions (cross-domain integration, novelty beyond the explicit literature, a stated falsification condition, differentiation from prior art, and a quantitative prediction), a list of forbidden patterns, and a strict JSON-only field set (title, statement, mechanism, evidence_basis, falsification, differentiation, quantitative_prediction, experiment, and self-scores); the Critic contract re-scores each hypothesis on the four criteria of Table 3, adds strength, weakness, and devil’s-advocate fields, and designates a top hypothesis, again JSON-only. (iii) The deterministic non-LLM control code is the control-code layer of Figure 2: set-difference routing in the Evidence Coordinator, the bounded (≤12-iteration) tool-use loop, the extract_json_str parser, and the additive-reducer state together make the pipeline reproducible, and only the Synthesizer and Critic differ by model tier. (iv) The ABCD-Stack (Section 2.4.4) is the enhancement layer of Figure 2: it adds four deterministic guards — a schema gate, a coverage audit, Jaccard diversity selection, and paired cross-family ICC scoring — together with three generation-time synthesizer personas (an integrative cross-domain bridger, a structure-driven hypothesizer, and a contrarian that challenges the field consensus) prepended to the same Synthesizer contract. Because all of these are executed by the same model, the robustness and diversity they add are properties of this prompt/architecture layer — not of the model tier — which is exactly the harness-control view that Figure 2 depicts.

**Figure 2.**
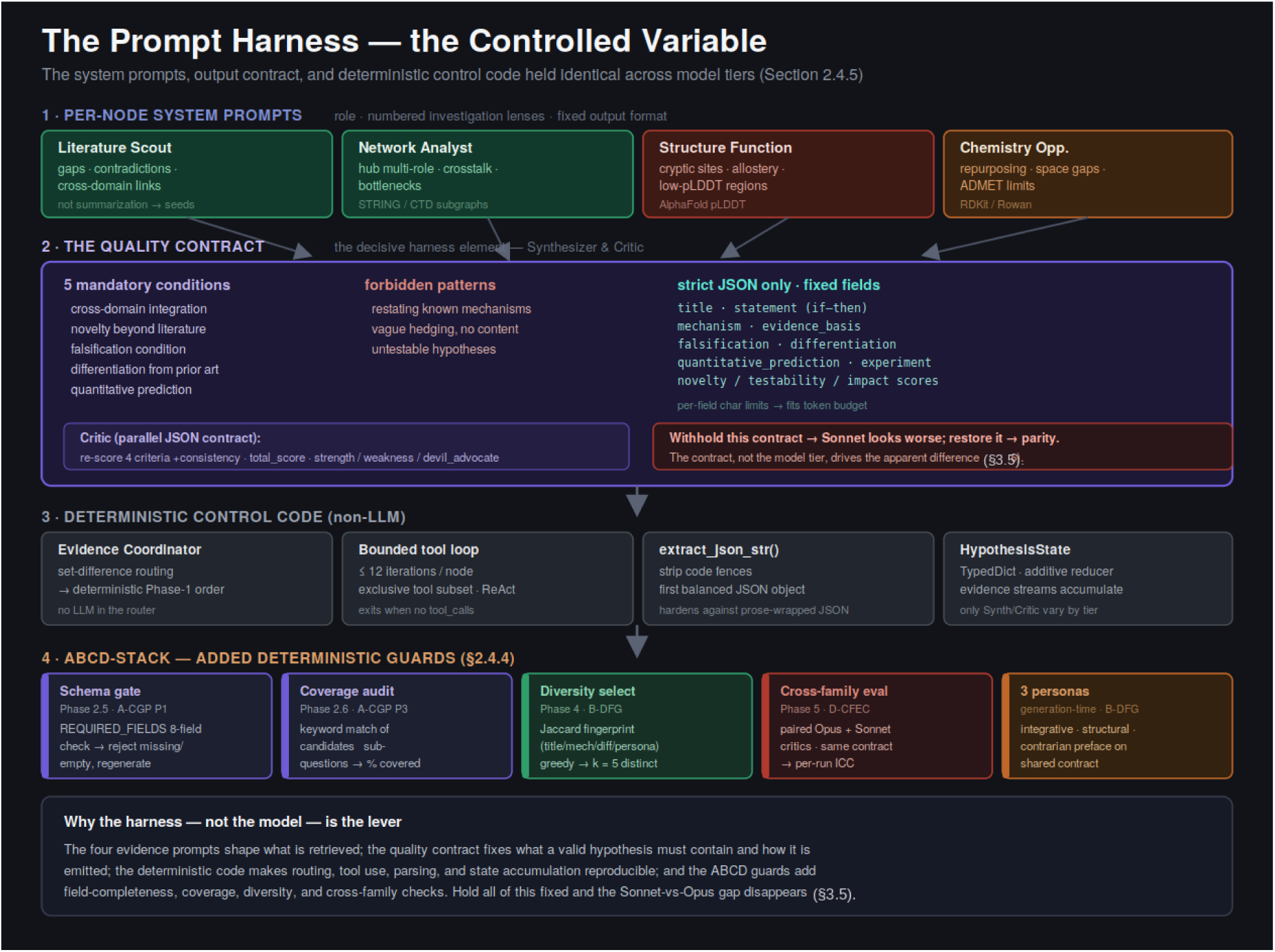
The prompt harness — the controlled variable. Four layers surround every LLM call and are held identical across model tiers in the matched-harness experiment (Figure 2). (1) Per-node system prompts give each of the four evidence agents a role, numbered investigation lenses, and a fixed output format, steering them toward hypothesis seeds rather than summaries. (2) The quality contract — the decisive element — states five mandatory conditions (cross-domain integration, novelty, falsification, differentiation, quantitative prediction), forbids restatement and vague hedging, and mandates strict JSON with a fixed field set and per-field character limits; the Critic carries a parallel contract. (3) Deterministic non-LLM control code (set-difference routing, a ≤12-iteration bounded tool loop, extract_json_str parsing, and an additive-reducer state) makes the pipeline reproducible; only the Synthesizer and Critic differ by tier. (4) The ABCD-Stack adds four deterministic guards — schema gate, coverage audit, Jaccard diversity selection, and paired cross-family ICC scoring — plus three generation-time personas. Holding all of this fixed is what makes the Sonnet-versus-Opus comparison a test of the model rather than the prompt (Section 3.5).

Per-node system prompts. Each of the seven nodes carries a fixed, role-specific system prompt with the same three-part structure: a role definition, a set of numbered investigation lenses, and a prescribed output format. The four evidence nodes are directed away from summarization and toward hypothesis seeds — the Literature Scout is told to report unresolved problems, contradictory findings, and unexpected cross-domain links rather than to summarize; the Network Analyst probes hub multi-functionality, pathway cross-talk, cross-disease modules, and intervention bottlenecks; the Structure Function agent targets cryptic binding sites, allosteric mechanisms, and the functional meaning of low-pLDDT disordered regions; and the Chemistry Opportunity agent looks for repurposing candidates, chemical-space gaps, and ADMET limits. Each evidence prompt fixes an output format with headed fields (e.g., current consensus, knowledge gap, contradictory findings, hypothesis seed), so that the four streams arrive in a comparable shape for synthesis.

The quality contract. The decisive element of the harness is the Synthesizer contract. Its system prompt states five mandatory conditions every hypothesis must meet — cross-domain integration of multiple evidence sources, novelty beyond the explicit literature, a stated falsification condition, explicit differentiation from prior art, and a quantitative prediction — and an explicit list of forbidden patterns, including restatements of known mechanisms and vague hedging language with no concrete content. It then mandates output as strict JSON only (no surrounding prose) with a fixed field set: title, statement (in if-then form), mechanism, evidence_basis, falsification, differentiation, quantitative_prediction, experiment, and three self-scores (novelty, testability, impact). Per-field character limits are imposed to keep the structured output within the token budget and prevent the nested-array overflow that unconstrained generation produced in early runs. The Critic node carries a parallel JSON-only contract, re-scoring each hypothesis on the four criteria (adding consistency), computing total_score, and adding strength, weakness, and a devil’s-advocate rebuttal before designating top_hypothesis_id. It is precisely this contract — the mandated fields and the JSON-only instruction — that, when withheld from the Sonnet baseline and then restored, accounts for the apparent model differences of Section 3.5.

Deterministic control code. The harness is not only prompts. Routing is handled by a non-LLM Evidence Coordinator that inspects which evidence agents have completed and returns the next one (or advances to synthesis) by simple set difference, making Phase 1 order deterministic. Each evidence node runs a hand-written tool-use loop bounded at twelve iterations: the tool-bound model is invoked, its tool_calls are executed against the node’s exclusive tool subset and fed back as ToolMessages, and the loop exits when the model returns no further tool calls. A shared extract_json_str routine strips Markdown code fences and extracts the first balanced JSON object, hardening the pipeline against models that wrap JSON in prose. State flows through a HypothesisState TypedDict whose evidence_results field uses an additive reducer so the four streams accumulate. Only the Synthesizer and Critic differ between configurations — in the Opus 4.8 setting they use the Bedrock Mantle client with adaptive thinking — while every prompt, tool subset, loop bound, and parser is held fixed.

Additional harness machinery in the ABCD-Stack. The ABCD-Stack (Section 2.4.4) adds four deterministic, non-LLM guards — a schema gate (Phase 2.5), a coverage audit (Phase 2.6), Jaccard diversity selection (Phase 4), and paired cross-family ICC scoring (Phase 5) — together with an adversarial Red-Team/Reviser refinement loop (Phase 3, C-ARL) and three generation-time synthesizer personas; these mechanisms are specified in full in Section 2.4.4 and are not repeated here. What matters for the present section is only their prompt-layer basis: every one of these guards enforces a guarantee that a system prompt has already made explicit — the schema gate mirrors the Synthesizer quality contract, the coverage audit scores against the Decomposer’s enumerated sub-questions, the Red-Team and Reviser are themselves system prompts, and the cross-family Critic reuses the single JSON-only Critic contract. All are reproduced verbatim in Supplementary Information S4.

### 2.5 Case Studies

Three independent research questions were evaluated:

1. **TP53-mutant solid tumors** — innovative drug discovery strategies for lung, colorectal, and pancreatic cancers harboring TP53 mutations, with focus on: literature gaps, unexpected network connections beyond MDM2/MDM4, cryptic AlphaFold binding sites, and chemical limitations of APR-246/Nutlin-3a [36,37,40,41].
2. **KRAS G12C pancreatic cancer** — novel therapeutic strategies to overcome sotorasib resistance [48–53], exploring: resistance mechanism gaps, MAPK/PI3K network vulnerabilities, structural targets beyond the GTP/GDP-binding site, and chemical challenges.
3. **Antibody manufacturing (CHO cells)** — genome-engineered cell line strategies to overcome genetic instability and lot-to-lot production variability [60,62,63].
4. **Novel iPSC reprogramming factors** — iPSC reprogramming, asking for Yamanaka-factor substitutes (small/medium-molecule or peptide candidates) with proposed mechanisms (the classical four-factor set is Oct4/Sox2/Klf4/c-Myc [74,75]; factor-substitution precedents include the nuclear receptor Nr5a2 replacing Oct4 [76] and a small-molecule TGF-β inhibitor replacing Sox2 [77])

Queries 1 and 2 (TP53, KRAS G12C) were executed on both Sonnet 4.6 (baseline) and Opus 4.8 (enhanced). Queries 3 and 4 were executed on both Sonnet 4.6 and Opus 4.8, with five independent repeats per model. After the experimental conditions were unified across case studies, all runs under both models applied the same 4-criterion schema including consistency (max 20).

### 2.6 Software and Data Availability

All notebooks, MCP server source code, ingestion scripts, and configuration files will be released in a public GitHub repository upon journal publication, after removal of all credentials; the generated hypotheses and per-run scores are provided in the Supplementary Information. MCP servers are packaged as Docker containers with `Dockerfile` and `requirements.txt` per server. Environment variables are documented in `config/.env.example`. STRING DB and CTD data were downloaded locally to avoid rate-limiting during agent runs.

## 3. Results

### 3.1 System Deployment and Tool Connectivity

The overall system architecture is shown in Figure 3, and the complete inventory of the 66 tools exposed by the five MCP servers is reproduced in Figure 5; the two figures are referenced together here because deployment correctness was confirmed both at the level of container/endpoint connectivity (Figure 4) and at the level of individual tool registration (Figure 5).

**Figure 3.**
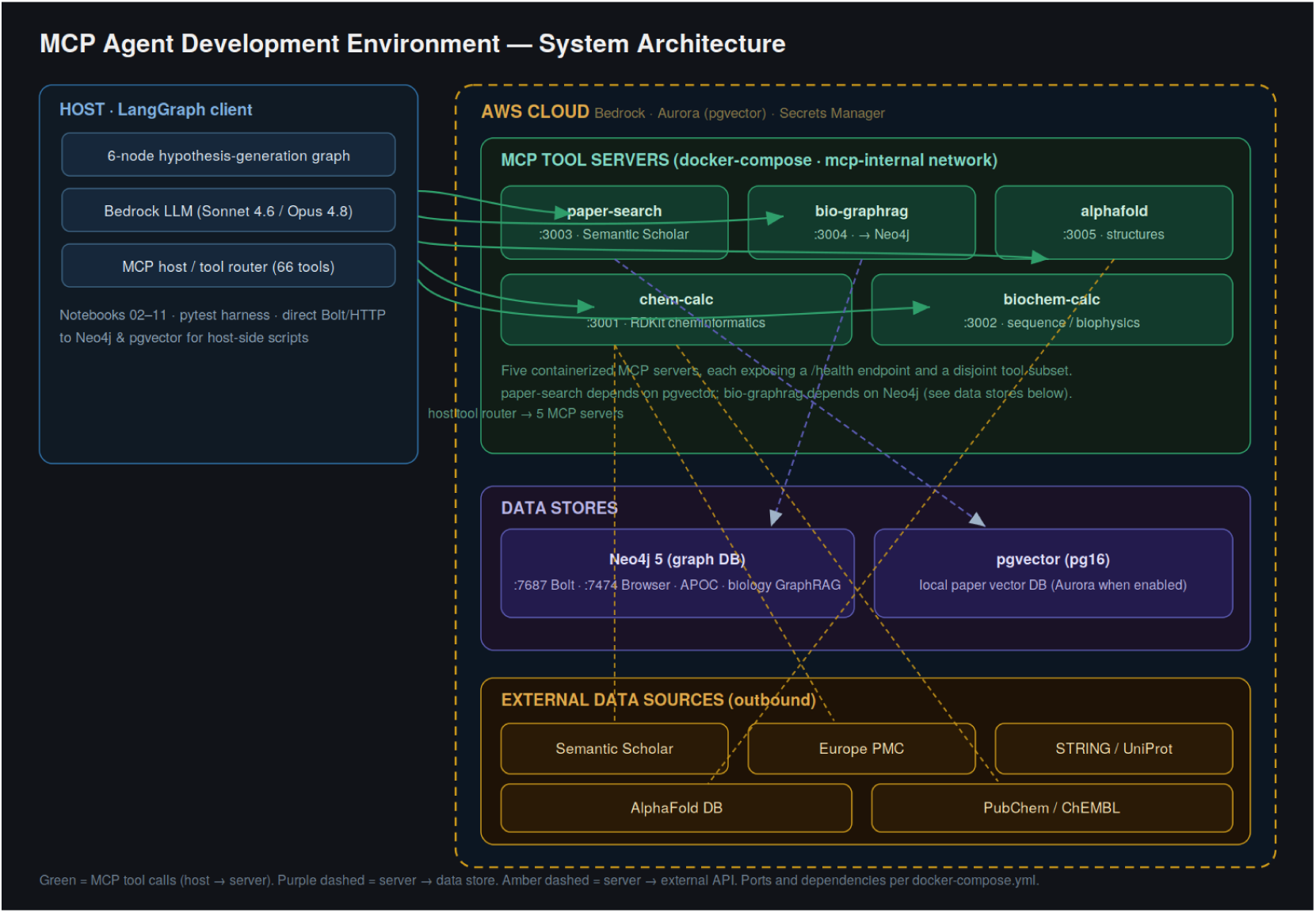
MCP agent development environment — system architecture. A client layer (JupyterLab on :8888, a LangGraph ReAct agent that is the sole MCP client over SSE, and built-in filesystem/memory MCP servers) drives the five Dockerized MCP servers — mcp-chem-calc (:3001), mcp-biochem-calc (:3002), mcp-paper-search (:3003), mcp-bio-graphrag (:3004), and mcp-alphafold (:3005) — inside a single LXC container (Ubuntu 24.04 LTS, Python 3.13, Docker Compose). Servers reach external public APIs (PubChem, UniProt, BioMCP, Semantic Scholar, AlphaFold, Rowan) and AWS Cloud backends (Amazon Bedrock for inference; Aurora PostgreSQL 14 + pgvector and Titan embeddings for semantic paper search), exposing 66 tools in total. The two backing data stores are shown below the server group: a Neo4j 5 graph database (Bolt :7687) for mcp-bio-graphrag and a pgvector store (Aurora when enabled) for mcp-paper-search. Arrow legend: green = MCP tool call (host → server); purple dashed = server → data store; amber dashed = server → external API. Ports and inter-service dependencies follow docker-compose.yml.

**Figure 4.**
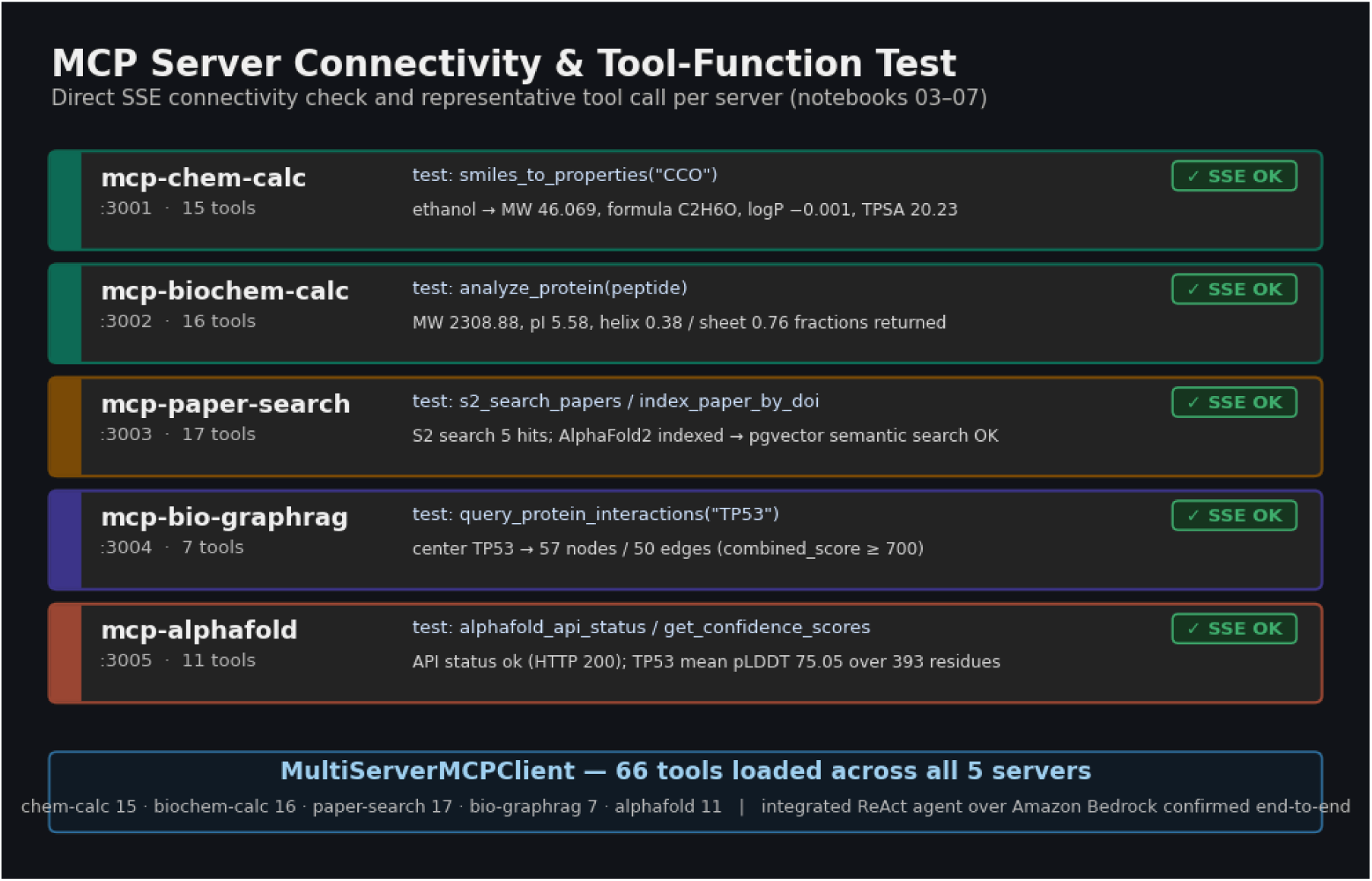
MCP server connectivity and tool-function test. For each of the five servers, a direct SSE session confirmed connectivity (✓ SSE OK), enumerated the registered tools, and invoked a representative tool against its live backend; observed results are shown per card. The footer records that MultiServerMCPClient loaded all 66 tools simultaneously and that the integrated ReAct agent over Amazon Bedrock operated end-to-end.

**Figure 5.**
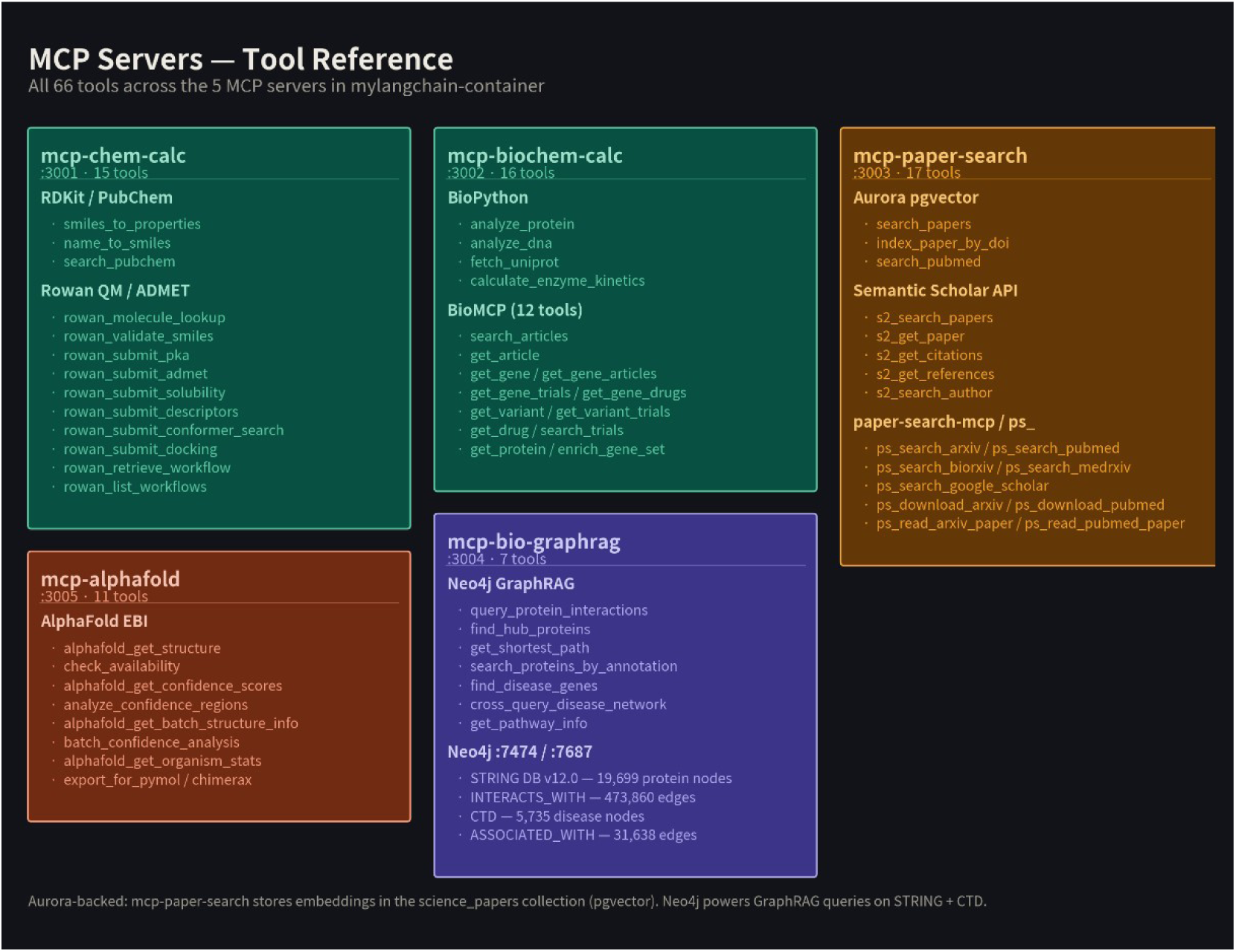
MCP server tool reference. All 66 tools distributed across the five MCP servers (mcp-chem-calc, 15; mcp-biochem-calc, 16; mcp-paper-search, 17; mcp-bio-graphrag, 7; mcp-alphafold, 11), grouped by backend (RDKit/PubChem, Rowan QM/ADMET, BioPython/BioMCP, Aurora pgvector, Semantic Scholar, paper-search-mcp, Neo4j GraphRAG, AlphaFold EBI). The Neo4j store underlying mcp-bio-graphrag holds 19,699 protein nodes (473,860 INTERACTS_WITH edges) and 5,735 disease nodes (31,638 ASSOCIATED_WITH edges).

The multi-agent development environment was successfully deployed as a single LXC container hosting five Dockerized MCP servers (Figure 3). Upon initialization, the `MultiServerMCPClient` dynamically loaded 66 tools across all five servers (mcp-chem-calc: 15, mcp-biochem-calc: 16, mcp-paper-search: 17, mcp-bio-graphrag: 7, mcp-alphafold: 11), confirming full connectivity to all downstream data sources including STRING DB [18], CTD [19], Aurora pgvector [24], and the AlphaFold EBI API [15] (Figure 4). The biological knowledge graph (Neo4j) contained 19,699 human protein nodes connected by 473,860 high-confidence INTERACTS_WITH edges (combined_score ≥ 700) and 31,638 disease–gene ASSOCIATED_WITH edges, enabling network-based evidence collection at genome scale.

#### 3.1.1 Per-Server Connectivity and Tool-Function Tests

Beyond the aggregate tool count, each MCP server was independently verified for connectivity and tool-level function before the agent pipeline was assembled (Figure 4). For each server, a direct SSE session enumerated the registered tools and a representative tool was invoked end-to-end against its live backend.

For mcp-alphafold (:3005), alphafold_api_status returned status ok (HTTP 200) against the AlphaFold EBI API [15]; the server registered all 11 tools, and alphafold_get_confidence_scores for TP53 (P04637) returned a mean pLDDT of 75.05 over 393 residues (207 very-high, 28 high, 41 low, 117 disordered), with a high-confidence DNA-binding-domain segment beginning at residues 96–114. For mcp-paper-search (:3003), all 17 tools registered; s2_search_papers returned five hits per query, index_paper_by_doi indexed a DOI into the Aurora pgvector store (status indexed, then already_indexed on repeat), and the subsequent search_papers semantic query retrieved the indexed record, confirming the embed-then-retrieve path; one Semantic Scholar call returned an HTTP 429 rate-limit, handled gracefully. For mcp-bio-graphrag (:3004), all 7 tools registered; query_protein_interactions for TP53 returned a 57-node, 50-edge neighborhood (BRCA1 returned 51 nodes / 50 edges), all at combined_score ≥ 700.

For mcp-biochem-calc (:3002), analyze_protein returned a complete physicochemical profile (e.g., molecular weight 2308.88, isoelectric point 5.58, with helix/sheet fractions) for a test peptide. For mcp-chem-calc (:3001), smiles_to_properties for ethanol (SMILES “CCO”) returned molecular weight 46.069, formula C2H6O, logP −0.001, and TPSA 20.23. Finally, the integrated MultiServerMCPClient loaded all 66 tools simultaneously (15 + 16 + 17 + 7 + 11), and a ReAct agent over Amazon Bedrock successfully routed queries to the correct servers (literature, network, structure, and chemistry tools), confirming end-to-end operation of the full tool ecosystem.

### 3.2 Baseline Multi-Agent Research

Before deploying the six-node hypothesis-generation pipeline, we first verified that the agents actually function end to end — that each specialist agent selects and calls the right tools, retrieves live evidence, and that the Coordinator integrates their outputs — by tracing the evidence each agent returned on real queries. To this end the baseline three-agent research system — the Literature, Bioinformatics, and Chemistry agents, integrated by a Coordinator node (Figure 1) — was evaluated on two well-characterized oncology targets, chosen because their resistance biology is documented well enough to check the retrieved evidence against the known literature. The two case studies below are reported as that functional evidence. In both, every agent autonomously established its MCP tool connections and completed its tool calls without manual intervention — the Literature Agent reaching the paper-search backends, the Bioinformatics Agent the Neo4j STRING/GraphRAG store, and the Chemistry Agent the RDKit/QM and AlphaFold services — and the Coordinator successfully consolidated the three tool-grounded evidence streams into a single integrated report. We therefore confirmed that the autonomous node-to-tool connectivity verified structurally in Section 3.1 (Figures 4 and 5) also operates correctly end to end on live scientific queries: the agents connect to their tools, retrieve real evidence, and hand it to synthesis as designed.

#### Imatinib/BCR-ABL in CML

The system integrated evidence from three independent domains within a single execution. The Literature Agent identified two key resistance papers (Quintás-Cardama et al., 2009 [57]; Bixby and Talpaz, 2009 [58]) and attributed ∼50% of imatinib [54] resistance to BCR-ABL kinase domain point mutations [55,56], particularly the T315I gatekeeper mutation. The Bioinformatics Agent queried the Neo4j STRING DB and ranked the top five BCR-ABL interactors: GRB2 (score 999), BCR (score 999), SRC (score 997), CRKL (score 994), PIK3R1 (score 991), mapping the full RAS/MAPK and PI3K/AKT co-activation landscape. The Chemistry Agent retrieved imatinib molecular properties (MW 493.6 Da, LogP 3.72, HBD 2, HBA 9, TPSA 86.3 Å²) and confirmed full compliance with Lipinski’s Rule of Five [65]. The AlphaFold confidence score for the ABL1 kinase domain was pLDDT 91.3 [15]. The integrated coordinator report identified T315I-directed third-generation TKI development (ponatinib) [59] as the primary unresolved clinical challenge.

#### TP53-mutant solid tumors

The system classified TP53 mutations into three therapeutic subtypes: gain-of-function (GOF) mutations (R175H, R248W) amenable to APR-246 reactivation [36,40]; loss-of-function (null) mutations requiring synthetic lethality via WEE1/ATR/PARP inhibition [41]; and wild-type TP53 tumors with MDM2 overexpression responsive to Nutlin-3a [37,38]. Network analysis identified MDM2 (score 999), MDM4 (score 998), ATM (score 997), BRCA1 (score 994), and RB1 (score 991) as top interactors [18]. AlphaFold structural analysis identified a high-confidence DNA-binding domain (residues 100–290, pLDDT 88.4) [15] harboring six hotspot mutations, and a tetramerization domain (residues 323–356, pLDDT 92.1).

### 3.3 Novel Hypothesis Generation — Sonnet 4.6

The six-node hypothesis generation pipeline (Figure 1) was applied to two oncology queries using Claude Sonnet 4.6, under the unified harness (three hypotheses, quality-criteria schema, four-criterion Critic scoring, max 20).

The two oncology queries’ top-ranked hypotheses and their Critic sub-scores are summarised in Table 5; the full structured records — statement, mechanism, evidence basis, falsification condition, quantitative prediction, and differentiation from prior art — are given in Supplementary Information S1 (detailed descriptions) and S3 (verbatim records). Briefly, Sonnet 4.6’s TP53 top hypothesis proposes a bifunctional USP7–DNMT1/BRD4 degrader that breaks two mutually reinforcing epigenetic loops at once (18/20), and its KRAS G12C top hypothesis an IRS1–PTPN11 (SHP2) insulin-microenvironment axis driving pancreatic-specific sotorasib resistance, testable with sotorasib ± TNO155 (19/20).

**Table 4.** ABCD-Stack enhanced pipeline. The four control principles and the pipeline phases that discharge each, layered on the shared four-agent evidence core and 66-tool MCP backend (Section 2.3). The scoring rubric adds a fifth criterion (faithfulness to the brief; Table 3), for a maximum of 25 points. Each condition-matched case study (CHO, iPSC) was run five times, producing the ten reports analyzed in Section 3.6.

| Stage | Principle | Phase(s) | Mechanism (deterministic guards in *italics in text*) |
| --- | --- | --- | --- |
| A-CGP | Contract-gated generation | 0, 2.5, 2.6 | Opus 4.8 Decomposer expands the brief into 5–8 sub-questions; a schema gate rejects and regenerates any field-incomplete candidate; a keyword coverage audit scores candidates against the sub-questions |
| B-DFG | Diversity-first generation | 2, 4 | Three synthesizer personas (integrative, structural, contrarian) generate in parallel; a greedy Jaccard-distance selector reduces the pool to $k = 5$ maximally distinct hypotheses |
| C-ARL | Adversarial refinement | 3 | A Red-Team prompt attacks each hypothesis and a Reviser hardens it against the attack |
| D-CFEC | Cross-family evaluation | 5 | Each hypothesis is scored independently by an Opus 4.8 critic and a Sonnet 4.6 critic on the five-criterion schema (max 25); the two are averaged and their intraclass correlation is reported |

**Table 5.** Oncology case studies: top-ranked hypothesis and Critic sub-scores. n = 1 independent run per model per query; scores on the unified four-criterion schema (novelty, testability, impact, consistency; each 1–5, max 20). N = novelty, T = testability, I = impact, C = consistency (each scored 1–5). Each row is the Critic-designated top (★) hypothesis of its run; full structured records in Supplementary S1 (Sonnet 4.6) and S3 (verbatim).

| Case | Model | Top-ranked hypothesis | N | T | I | C | Total |
| --- | --- | --- | --- | --- | --- | --- | --- |
| TP53 | Sonnet 4.6 | USP7–DNMT1/BRD4 dual epigenetic-loop disruption | 4 | 5 | 4 | 5 | 18/20 |
| TP53 | Opus 4.8 | Oxidative-stress-dependent GOF & TrxR1 vulnerability | 5 | 5 | 4 | 5 | 19/20 |
| KRAS G12C | Sonnet 4.6 | IRS1–PTPN11 pancreatic-specific resistance axis | 5 | 5 | 4 | 5 | 19/20 |
| KRAS G12C | Opus 4.8 | switch-II flexibility, pre-emptive ON-state inhibition | 4 | 5 | 4 | 5 | 18/20 |

### 3.4 Enhanced Hypothesis Generation — Opus 4.8 + Adaptive Thinking

Claude Opus 4.8 with Adaptive Thinking was run on the same two oncology queries as Sonnet 4.6 (Table 5). Sonnet 4.6 and Opus 4.8 were then run on two further queries — CHO cell engineering and iPSC reprogramming — with five independent repeats each (Table 6). The eight top-ranked hypotheses (one per model for each of the four case studies) are collected in Figure 6. Full structured records for all top hypotheses are given verbatim in Supplementary Information S3, and the complete per-run detail in Supplementary S1–S2. Because the two oncology queries were run once per model (Table 5), no significance test is defined for them; the five-repeat CHO and iPSC queries (Table 6) are the basis for the statistical comparison, and there the only significant difference favours Sonnet 4.6 on CHO (18.4 vs 17.4, Welch t = 2.89, p < 0.05), while iPSC is statistically indistinguishable (18.6 vs 18.0, t = 1.50, n.s.).

**Figure 6.**
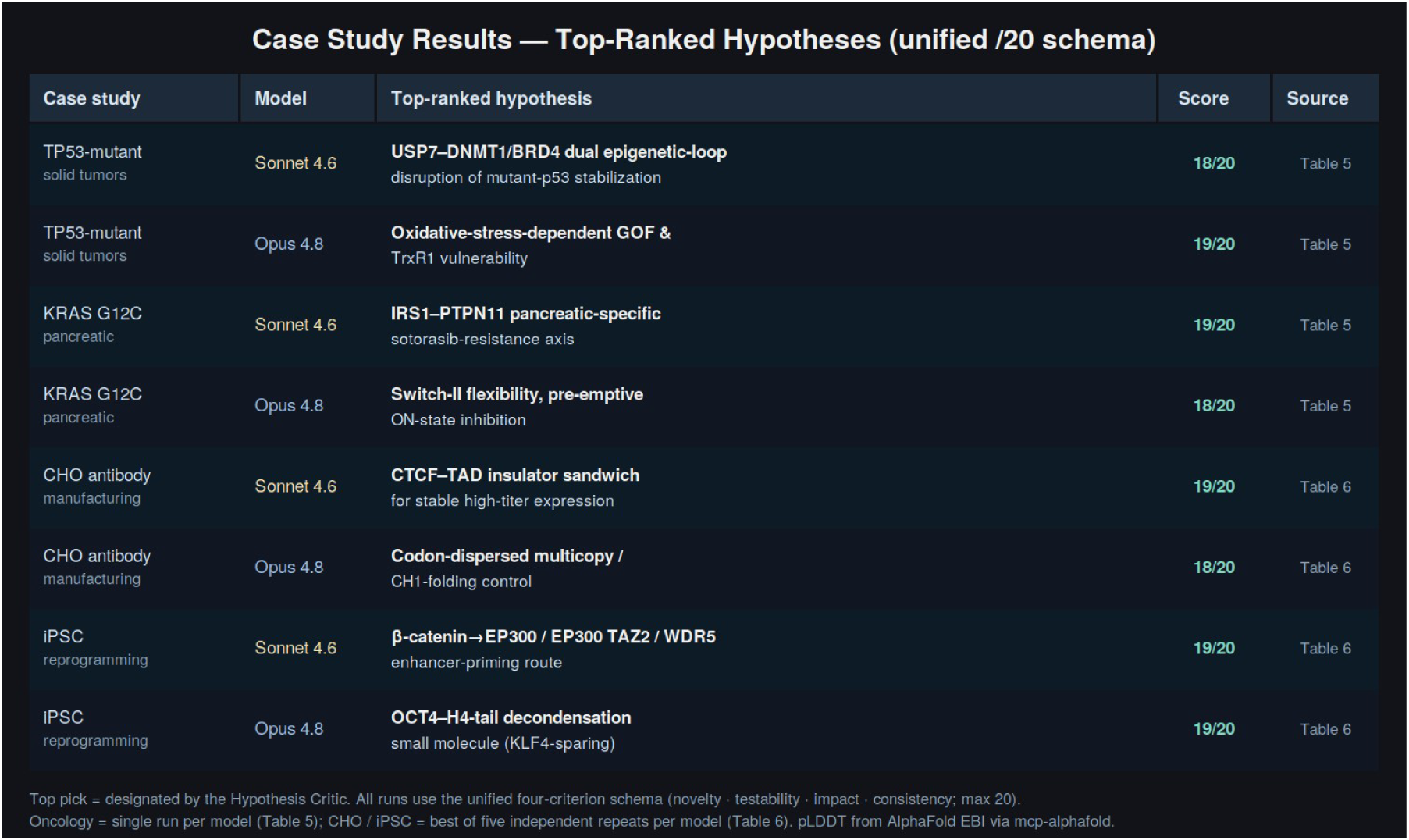
Case study results — AI-generated scientific hypotheses. Summary of the eight top-ranked hypotheses — one per case study for each of the two models — across the four case studies (case study, model, top-ranked hypothesis, total score, and source table). ★ = top-ranked by the Hypothesis Critic. All runs under both models use the unified 4-criterion schema including consistency (max 20). AlphaFold pLDDT values from the AlphaFold EBI API via mcp-alphafold.

**Table 6.** Manufacturing and reprogramming case studies: top-hypothesis total score (mean ± SD) over independent repeats per model, on the unified four-criterion schema (max 20). n = 5 independent repeats per model per case study (10 reports per case). Values are mean ± SD of the top-hypothesis total score across the n = 5 repeats. CHO: Sonnet 4.6 18.4 ± 0.5 vs Opus 4.8 17.4 ± 0.5, Welch’s t = 2.89, df ≈ 8, p < 0.05 (Sonnet higher); iPSC: Sonnet 4.6 18.6 ± 0.5 vs Opus 4.8 18.0 ± 0.7, Welch’s t = 1.50, df ≈ 8, n.s. Full records in Supplementary S2–S3.

| Case | Model | Mean ± SD | Best run (hypothesis, score) |
| --- | --- | --- | --- |
| CHO | Sonnet 4.6 | 18.4 ± 0.5 | CTCF–TAD insulator sandwich, 19/20 |
| CHO | Opus 4.8 | 17.4 ± 0.5 | codon-dispersed multicopy / CH1 folding, 18/20 |
| iPSC | Sonnet 4.6 | 18.6 ± 0.5 | β-catenin → EP300 / EP300 TAZ2 / WDR5, 19/20 |
| iPSC | Opus 4.8 | 18.0 ± 0.7 | OCT4–H4-tail decondensation molecule, 19/20 |

Expected effect of Adaptive Thinking. Adaptive (extended) Thinking is Anthropic’s inference-time reasoning mode: instead of answering immediately, the model is given a token budget to plan, explore alternatives, and self-check on an internal scratchpad before emitting its final response, with the depth of reasoning scaled to task difficulty and controlled by a thinking budget [78]. Anthropic positions it for precisely the class of work this pipeline performs — multi-step analysis and decision tasks that weigh several paths — while cautioning that it adds latency and can even degrade fluency-driven tasks, so its benefit is task-dependent rather than universal [78]. For hypothesis generation the expected contribution is threefold: at synthesis, the Synthesizer can enumerate and compare several candidate mechanisms, reconcile the four evidence streams, and pre-test each hypothesis against its own falsification condition before committing to the JSON contract; at critique, the Critic can reason through the rubric rather than pattern-match a score; and across both, the added deliberation is expected to lower run-to-run variance. Whether these expected gains actually materialise as higher or more reliable scores is tested in Sections 3.5–3.5.1; the first-pass results here already show no large or significant advantage for the Opus configuration, and the matched-harness re-test attributes what differences remain to the prompt contract rather than to Adaptive Thinking itself (Section 3.5).

Across the two oncology queries, Opus 4.8’s top hypotheses were a redox-conditional GOF/TrxR1 vulnerability for mutant p53 (19/20) and a switch-II-flexibility, pre-emptive ON-state inhibition strategy for KRAS G12C resistance (18/20). For the two five-repeat queries the top hypotheses spanned secretion-flux and genome-stability designs for CHO and druggable-handle substitutes for the Yamanaka factors in iPSC. Taken together with Table 6, the first-pass comparison shows the two models to be close: no large or significant Opus advantage is seen, the sole significant difference favouring Sonnet 4.6 on CHO; the model-level score comparison and the strategy analysis are developed in Sections 3.5–3.5.1.

### 3.5 Repeated-Run Analysis: Score Parity but Reproducible Strategic Divergence

To separate genuine model differences from run-to-run noise, we repeated the two condition-matched case studies (CHO and iPSC) five times each, independently, on both Sonnet 4.6 and Opus 4.8 (the latter with Adaptive Thinking), yielding twenty reports and a per-condition distribution of top-hypothesis scores. The repeated data revise one claim of the single-run comparison in Section 3.4 and strengthen another (Figures 6 and 7).

**Figure 7.**
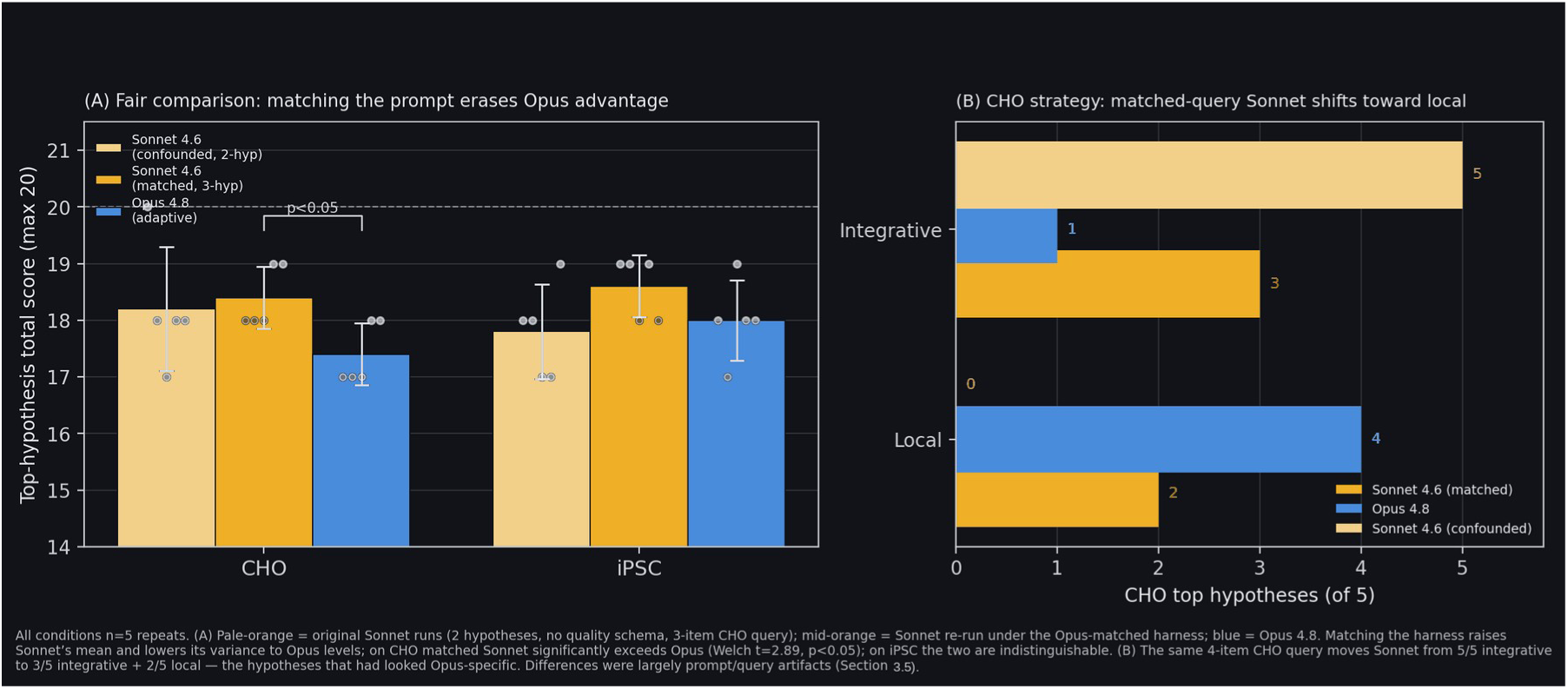
Score parity under a matched harness: equal hypothesis quality once the prompt is controlled (prompts, output schema, hypothesis count, and queries identical; Sonnet 4.6 held as the model). Five independent repeats per condition. (A) Top-hypothesis total score. Pale-orange = original Sonnet 4.6 runs (two hypotheses, no quality schema, three-item CHO query); mid-orange = Sonnet 4.6 re-run under the Opus-matched harness (three hypotheses, quality schema, four-item CHO query); blue = Opus 4.8 with Adaptive Thinking. Matching the harness raises Sonnet’s mean and lowers its variance to Opus levels; on CHO, matched Sonnet significantly exceeds Opus (18.4 vs 17.4, Welch t = 2.89, p < 0.05), and on iPSC the two are statistically indistinguishable (t = 1.50). Points are individual runs. (B) CHO strategy coding, showing that the hypothesis search space is controlled by the query’s sub-questions. Adding one sub-question to the CHO query — an explicit antibody-structural-design item, so that the matched query carries four sub-questions instead of the original three — moves Sonnet 4.6 from 5/5 integrative (pale) to 3/5 integrative + 2/5 local (mid-orange), reproducing the very local antibody-molecule hypotheses that had appeared Opus-specific (blue). Because adding the sub-question shifts the coded strategy in a controlled way, the region of hypothesis space each model explores is set by the sub-questions enumerated in the query, not by the model: the apparent integrative-versus-local model difference of earlier drafts was a query-coverage artifact of the harness (Section 3.5).

Figure 7 makes the matched-harness comparison explicit and is the central control of this section. The two Sonnet 4.6 bars in Figure 7A — the original configuration and the Opus-matched one (defined in the legend) — sit side by side for each case, so the gap between them isolates the effect of the harness alone, since the model is held fixed while only the prompt, schema, hypothesis count, and query change. Matching the harness raises Sonnet’s mean and compresses its run-to-run variance to Opus levels; once matched, Sonnet 4.6 significantly exceeds Opus 4.8 on CHO (18.4 vs 17.4, Welch t = 2.89, p < 0.05) and is statistically indistinguishable from it on iPSC (t = 1.50, n.s.). Figure 7B shows why earlier drafts had read a model-level strategy difference here. Each top hypothesis is coded on an intervention-scale axis — integrative (a single intervention proposed to resolve two or more distinct cellular phenomena at once, e.g. a CTCF–TAD insulator that couples chromatin insulation to expression stability) versus local (confined to one residue, domain, or pathway, e.g. a BiP linker point-mutant that retunes secretion). The original Sonnet 4.6 CHO runs were 5/5 integrative; giving Sonnet the same four-item CHO query — which adds an antibody-structural-design sub-question absent from the original prompt — moves it to 3/5 integrative and 2/5 local, reproducing precisely the local antibody-molecule hypotheses that had appeared Opus-specific. The apparent integrative-versus-local model difference was therefore a query-coverage difference in the harness, not a property of the model. Per-repeat scores and the integrative/local coding for all five repeats of each configuration are given in Supplementary Information S5.

#### 3.5.1 Experimental Setup and Overall Scores

Near-parity under the matched harness. Across five repeats on the unified max-20 schema the top-hypothesis totals are: CHO Sonnet 18.4 ± 0.5, CHO Opus 17.4 ± 0.5, iPSC Sonnet 18.6 ± 0.5, iPSC Opus 18.0 ± 0.7 (mean ± SD, max 20). On iPSC the two models are statistically indistinguishable (Welch t = 1.50, n.s. at α = 0.05); on CHO there is a small but significant edge to Sonnet 4.6 (Welch t = 2.89, p < 0.05). The single-run “CHO inversion” highlighted earlier (Sonnet’s CTCF–TAD insulator at 19/20 vs. an Opus local design at 17/20) sits at the high end of this distribution; across the five matched repeats the gap is about one point in Sonnet’s favour, and neither model dominates across both cases. We therefore retract any claim that Opus 4.8 produces higher-scoring hypotheses than Sonnet 4.6 on these tasks; on the unified rubric the two are at most one point apart on average, with the only significant difference favouring Sonnet on CHO.

As for reliability convergence under the matched harness, an earlier draft attributed lower run-to-run variance to Adaptive Thinking, but once Sonnet 4.6 is given the same quality-criteria schema its SD falls to Opus levels: CHO 0.5 (Sonnet) vs 0.5 (Opus) and iPSC 0.5 vs 0.7. The apparent reliability gap was therefore a property of requiring a structured quality schema — obtainable from either model — not of the model tier or of Adaptive Thinking. Reliability is thus not a differentiator between the two configurations once the harness is matched.

#### 3.5.2 Strategic Divergence: Integrative vs. Local

Before presenting the coding results, we define the intervention-scale axis on which each top hypothesis is classified, because the integrative-vs-local distinction is the substantive claim of this section. The axis is biological, not stylistic, and a hypothesis is coded on the basis of the causal structure of the mechanism it proposes. A hypothesis is integrative when a single molecular intervention is proposed to act through a network hub or shared regulatory node so as to resolve two or more distinct cellular phenomena at once — for example coupling chromatin insulation to transcriptional stability, or driving an entire pluripotency circuit through one super-hub — such that the predicted effect propagates across pathways rather than staying within one. Operationally, an integrative hypothesis (i) names a hub, bottleneck, or multi-pathway coordinator (e.g. a STRING two-hop connector such as EP300 or β-catenin, a TAD-boundary insulator, or a master regulator), (ii) makes a prediction that spans two or more otherwise-separate processes (e.g. genome stability and expression level, or self-renewal and metabolic state), and (iii) requires a systems-level readout (network, multi-omics, or phenotype) to falsify. A hypothesis is local when the proposed intervention is confined to a single residue, domain, binding pocket, or linear pathway and its predicted effect stays within that one process — for example a point mutation that retunes one chaperone’s cycling rate, a rigidified antibody domain, or a single ligand-binding pocket. Operationally, a local hypothesis (i) targets one structurally defined element (a residue, motif, domain interface, or pocket), (ii) predicts an effect on a single process (secretion rate, folding, one enzyme’s activity), and (iii) can be falsified by a focused biophysical or single-gene assay. The two categories are mutually exclusive at the level of the top-ranked hypothesis and are assigned by a single coder from the mechanism and quantitative-prediction fields of each record (inter-coder reliability is addressed in Experiment 1 of Section 4.5.3). With that definition fixed, the coding results follow.

Coding the top-ranked hypothesis of each of the five repeats per case shows the split is not a single-run artifact. On CHO, Sonnet 4.6 produced an integrative, genome-/epigenome-stability hypothesis in three of five runs and two more local translation/secretion designs, whereas Opus 4.8 produced a local, structurally defined design in all five. The iPSC repeats reproduce the same intervention-scale tilt: Sonnet 4.6 again favoured integrative, multi-factor hub interventions (e.g. EP300/β-catenin coordination) while Opus 4.8 converged on single, structurally tractable handles. Across both cases Sonnet 4.6 is coded integrative in 8/12 top hypotheses and Opus 4.8 local in 10/12 — a reproducible tilt in preferred search region, not a difference in score (full per-run codings in Supplementary S5.2).

**Figure 8.**
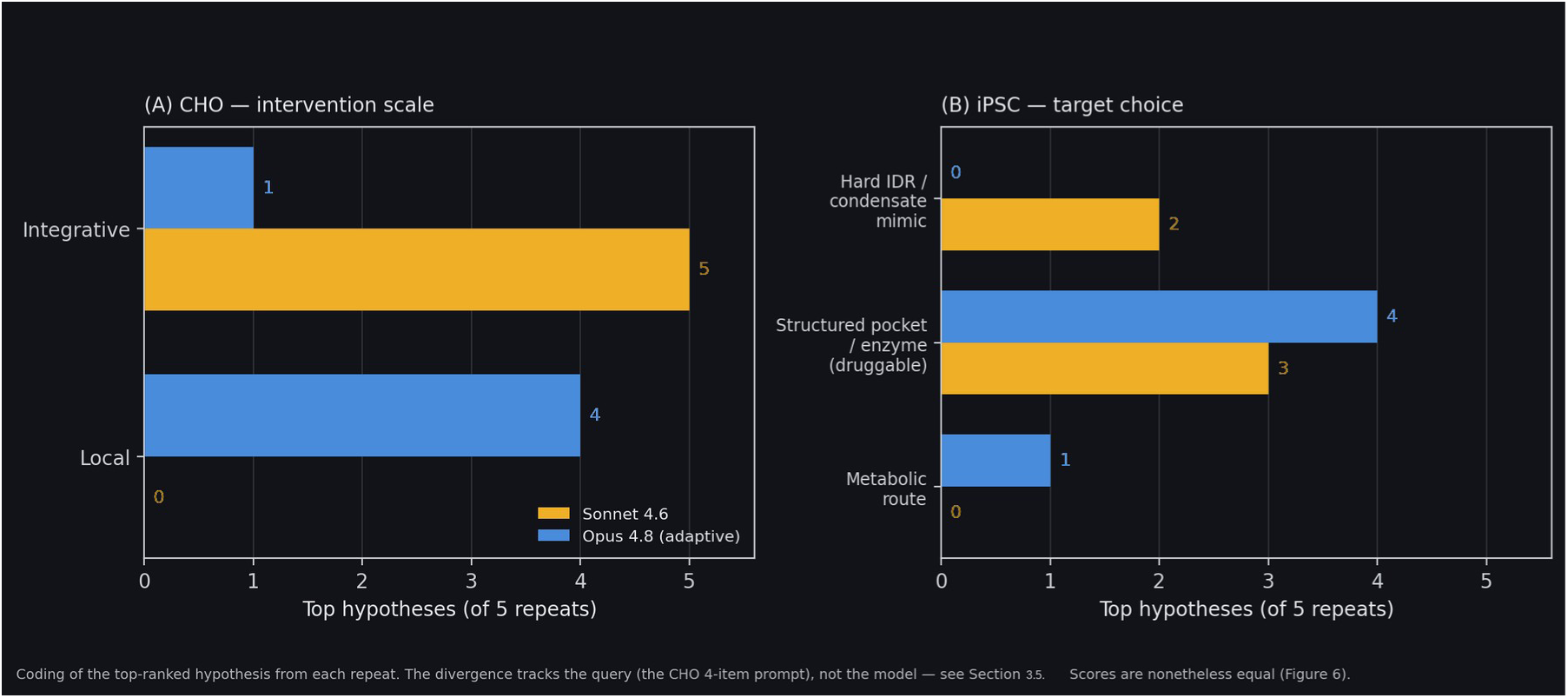
Reproducible strategic divergence at equal score: the same matched runs as Figure 7, now coded for intervention scale rather than scored (five matched repeats each; coding of the top-ranked hypothesis from each repeat). Whereas Figure 7B showed that adding or matching a sub-question moves a model’s coded strategy — so that the sub-questions control the search space — this panel isolates what remains once the sub-questions are held identical. With the queries and their sub-questions matched across the two models, the residual divergence cannot be attributed to a sub-question difference; instead it reflects how the breadth of the search space already implied by the query itself is taken up along each model’s own preferred search axis. Given the same matched brief, the two models score alike (Figure 7) yet distribute their top hypotheses along different intervention-scale axes, so the divergence here is a model-specific exploration tendency operating within a shared, query-defined search space rather than a consequence of differing sub-questions. (A) CHO intervention scale: Sonnet 4.6 produced an integrative, genome-/epigenome-stability hypothesis in 3/5 runs (answering the brief’s explicit chromosomal requirement), whereas all 5/5 Opus 4.8 CHO tops are local designs. (B) iPSC intervention scale: Sonnet 4.6 reached for integrative, multi-factor hub interventions in 3/5 runs (EP300/β-catenin, EP300 TAZ2, CLOCK/BMAL1–KDM1A), whereas Opus 4.8 chose a single local structural handle in 4/5. Across both case studies Sonnet 4.6 is coded integrative in 8/12 top hypotheses and Opus 4.8 local in 10/12: Sonnet tilts toward cross-phenomenon unification, Opus toward residue-level tractability. Total scores are nonetheless close and, on iPSC, statistically indistinguishable (Figure 7). Coding is single-coder; inter-coder reliability is addressed in Experiment 1 of Section 4.5.3.

Taken together, the repeated runs sharpen the interpretation: the two models are equally strong scorers with equal run-to-run variance under the matched harness, but they explore different regions of hypothesis space (integrative network hubs for Sonnet 4.6, local structural handles for Opus 4.8). The faithfulness blind spot noted earlier also persists across repeats — Opus’s recurrent off-brief (secretion/molecule) hypotheses continue to score on par with Sonnet’s on-brief (chromosomal) ones — reinforcing, rather than resting on a single case, the motivation for the fifth criterion proposed in Section 4.5.3. These two axes are complementary in a way that is directly useful for biology: the integrative mode assembles a novel hypothesis by combining several lines of reasoning across literature, network, and disease evidence, whereas the local mode grounds its hypothesis in structural and chemical computation. Assigning the two axes to parallel agents lets the system span domains that a single wet-lab investigator, trained deeply in one, rarely bridges, surfacing cross-domain questions that a single-viewpoint approach tends to miss. Neither axis is strictly better; because peak scores, hypothesis counts, and variance are equal once the harness is matched, the value lies in the coverage of complementary search regions rather than in a superior model.

#### 3.5.3 Rubric Ceiling and the Most Parsimonious Reading

A further consequence concerns the rubric itself. Across the twenty repeated runs of Section 3.5.1 (two case studies × two models × five repeats; Table 6), all twenty scores fall in a narrow 16–20 band, and every same-family Critic score clusters at 17–18 (SD ≤ 1.1). This compression is consistent with a ceiling and self-scoring effect: a Critic evaluating its own family’s output has limited dynamic range, which caps observable differences. This is the confound that the blinded cross-model scoring protocol of Section 4.5.3 (Experiment 2) is designed to break, and the variance reported here is therefore a lower bound on true variability, since it excludes evaluator disagreement.

Because the two agents share an identical six-stage architecture and differ only in the swapped Synthesizer/Critic model (Sonnet 4.6 versus Opus 4.8 with Adaptive Thinking), any residual divergence must originate in that one component. An earlier draft read three “signatures” off the outputs and attributed them to the model; under the matched harness each instead tracks a specific harness element. (i) Structured falsification reasoning was an output-contract effect: once both configurations emit the same mandatory fields, both produce them in their underlying JSON, and the earlier impression reflected how two report templates rendered the same schema. (ii) Exploration breadth (“three versus two hypotheses”) was a prompt effect: the original Sonnet prompt requested only two hypotheses (Supplementary S5), and under the matched harness both return three. (iii) Reliability was a schema effect: run-to-run variance converges once Sonnet is given the same quality-criteria schema (SD 0.5 on CHO for both models; 0.5 versus 0.7 on iPSC). None of the three is a reasoning-depth signature.

The most economical reading is therefore not that Adaptive Thinking makes Opus 4.8 more reliable or higher-ceiling — peak scores, hypothesis counts, and run-to-run variance are all equal once the harness is matched — but that the two models search different regions of hypothesis space (Section 3.5.2): Opus 4.8 converges on local, structurally tractable handles (a defined pocket, interface, or residue range), while Sonnet 4.6 reaches for integrative, network-hub interventions. Neither is strictly better — Opus 4.8’s local targets suit acting on a single structurally checkable lead, Sonnet 4.6’s integrative search suits early-stage ideation — and this complementarity in search region, not any difference in score, reliability, or reasoning schema, is the surviving finding of the model comparison. Even it is provisional pending the inter-coder and cross-scoring experiments of Section 4.5.3.

For every top-ranked hypothesis, the Synthesizer emits a complete structured record in the native schema (mechanism, evidence_basis, falsification, quantitative_prediction, experiment, and Critic scores). These records are the system’s primary output and the raw material for the validation program of Section 4.5; the verbatim records appear in Supplementary Information S3 and the full per-run narratives in S1–S2.

### 3.6 An Enhanced Architecture Under Harness Control: The ABCD-Stack

The controlled re-test of Section 3.5 established that output quality, count, reliability, and apparent research strategy are governed by the prompting harness rather than the LLM tier, and Section 4.6 abstracted that finding into four design principles — contract-gated output, diversity-first generation, adversarial refinement, and cross-family evaluation. Rather than leave those principles as recommendations, we implemented them as an executable pipeline, the ABCD-Stac, and re-ran the two condition-matched case studies (CHO and iPSC) under it. This section reports that experiment; it is the direct test of whether making the harness principles architectural, rather than incidental, changes what the system produces.

#### 3.6.1 Rationale and Architecture: From Diagnosed Harness Weaknesses to a Testable System

Aim. The ABCD-Stack was built to turn the harness-engineering findings of Sections 3.4–3.5 into a falsifiable engineering test. Those experiments did more than show that the prompting harness dominates the LLM’s reasoning capability; they localized four specific weaknesses in the baseline harness that a stronger harness ought to fix: (i) scoring was self-assigned by a critic of the same model family, an uncontrolled bias flagged throughout Section 3.5; (ii) the rubric lacked a faithfulness-to-brief axis, the blind spot diagnosed in Section 3.5; (iii) a single Synthesizer could collapse the candidate pool onto one line of reasoning, limiting diversity; and (iv) candidates were sometimes field-incomplete or left parts of the brief uncovered, with no mechanical check. We therefore treated each weakness as a design requirement and asked a concrete validation question: if the four harness-control principles (Section 4.6) are made architectural — deterministic contract-gating, diversity-first generation, adversarial refinement, and cross-family evaluation — rather than left as prompt-level recommendations, does the system measurably close these gaps on the same condition-matched case studies? The architecture below is the apparatus for that test; Sections 3.6.2–3.6.5 report the result.

Two distinct labelling schemes appear below, and we keep them separate to avoid confusion. The first is the set of four architectural pillars, labelled A–D, which name the pipeline stages: A-CGP (Contract-Gated Prompting), B-DFG (Diversity-First Generation), C-ARL (Adversarial Refinement Loop), and D-CFEC (Cross-Family Evaluation with Consistency check). Each phase in the pipeline below is tagged with the pillar it belongs to.

The second scheme is a set of three design principles, labelled P1–P3, that state what the harness must guarantee — independently of which pillar enforces them. P1, schema enforcement: every hypothesis must populate a fixed set of mandatory fields (mechanism, falsification, quantitative prediction, evidence basis) or be rejected and regenerated. P2, diversity by construction: breadth must be guaranteed by running several fixed persona prompts in parallel rather than by hoping a single prompt explores widely. P3, coverage accountability: the brief must be decomposed into an enumerated set of sub-questions so that the fraction actually addressed can be audited mechanically. These principles cut across the pillars rather than nesting inside one of them — P1 and P3 are enforced by the contract-gating machinery (A-CGP, at the schema gate and the coverage audit), while P2 is realized by the diversity-first stage (B-DFG). In the phase descriptions and in Figure 9 each phase is therefore tagged both with its pillar (A–D) and, where applicable, with the principle (P1–P3) it discharges.

**Figure 9.**
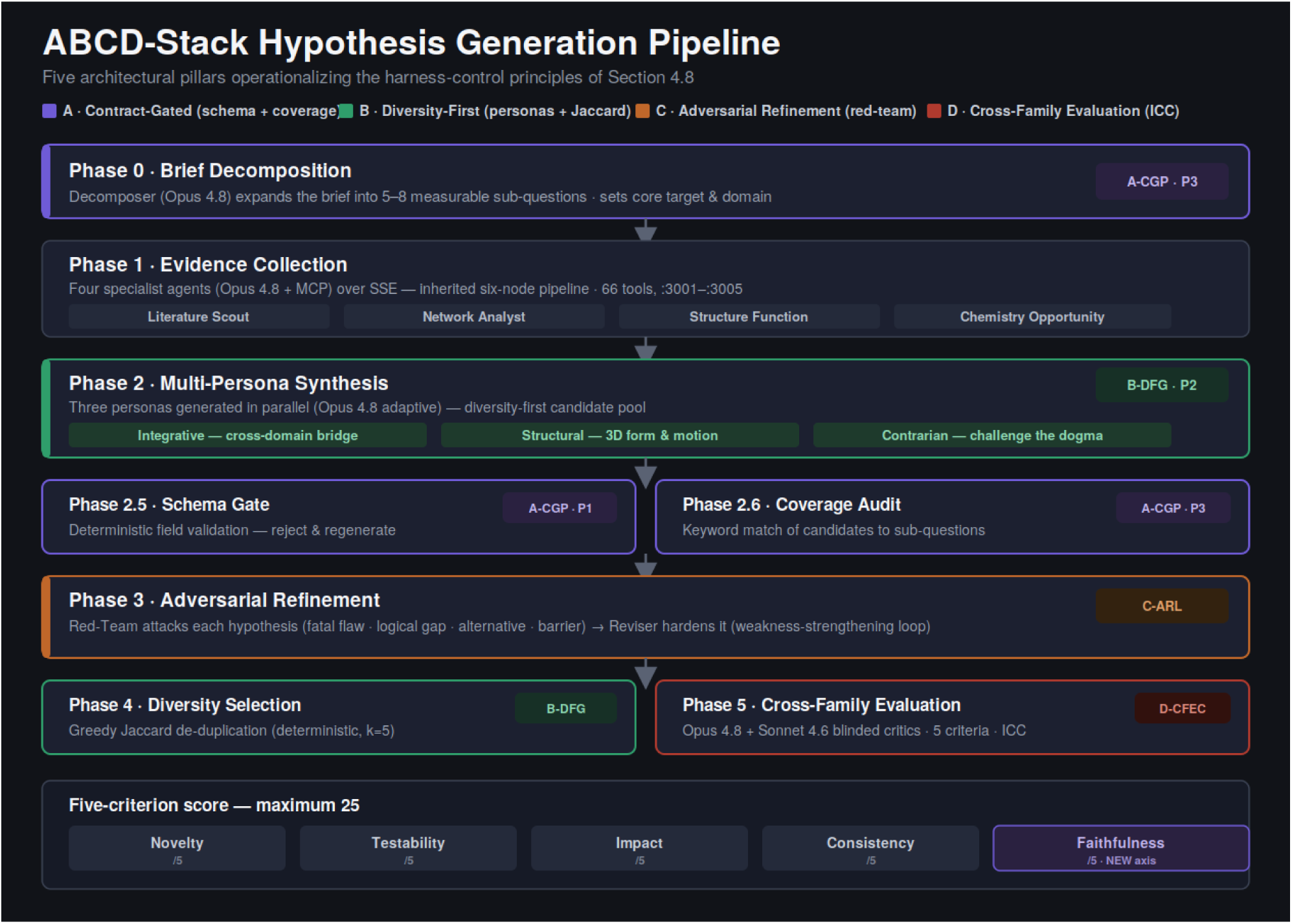
ABCD-Stack hypothesis-generation pipeline. The four architectural pillars (A–D) operationalize the harness-control principles of Section 4.7 across the pipeline phases, and each phase is tagged with its pillar and, where applicable, the design principle (P1–P3) it discharges. Phase 0 decomposes the brief (A-CGP P3); Phase 1 collects evidence over the inherited six-node core; Phase 2 synthesizes candidates from three personas in parallel (B-DFG P2); Phases 2.5/2.6 enforce a deterministic schema gate and coverage audit (A-CGP P1/P3); Phase 3 hardens each hypothesis through a Red-Team/Reviser loop (C-ARL); Phase 4 selects k = 5 by greedy Jaccard de-duplication (B-DFG); Phase 5 scores with paired Opus 4.8 and Sonnet 4.6 critics and reports their ICC (D-CFEC). The scoring rubric adds a fifth criterion — faithfulness to the brief — closing the blind spot identified in Section 3.5, for a maximum of 25 points.

Phase 0 (A-CGP, contract-gated decomposition, principle P3) runs first: an Opus 4.8 Decomposer reads the research brief and expands it into an enumerated set of five to eight measurable sub-questions naming a single core target and domain, which become the reference against which Phase 2.6 later audits coverage. Phase 1 inherits the four specialist agents (Literature Scout, Network Analyst, Structure Function, Chemistry Opportunity) over SSE against the same 66 tools. Phase 2 (B-DFG, diversity-first generation, principle P2) replaces the single Synthesizer with three personas run in parallel — an integrative cross-domain bridger, a structure-driven hypothesizer, and a contrarian that challenges the field consensus — producing a diverse candidate pool rather than one “best” answer. Phase 2.5 (A-CGP P1) is a deterministic schema gate that validates every required field and rejects and regenerates any candidate that omits a falsification condition, quantitative prediction, or evidence basis; Phase 2.6 (A-CGP P3) audits keyword coverage of the candidate pool against the Phase 0 sub-questions. Phase 3 (C-ARL, adversarial refinement loop) sends each surviving hypothesis to a Red-Team prompt that names its fatal flaw, logical gap, most plausible alternative explanation, and principal experimental barrier, and then to a Reviser that hardens the hypothesis against those attacks. Phase 4 (B-DFG) performs greedy Jaccard de-duplication to select k = 5 maximally distinct hypotheses. Phase 5 (D-CFEC, cross-family evaluation with consistency check) scores each hypothesis with two independent critics from different model families — Opus 4.8 and Sonnet 4.6 — and reports their agreement as an intraclass correlation (ICC).

Two rubric changes accompany the architecture. First, the score adds a fifth criterion, faithfulness to the brief (does the hypothesis answer the question as posed?), the exact axis whose absence Section 3.5 identified as a blind spot; the maximum score is therefore 25, not 20. Second, scoring is no longer self-assigned by a single same-family critic: every hypothesis is scored by both an Opus 4.8 and a Sonnet 4.6 critic, and the two are averaged, directly addressing the self-scoring confound flagged throughout Section 3.5.

#### 3.6.2 Repeated-Run Results

We executed the ABCD-Stack five independent times on the CHO brief and five on the iPSC brief, yielding ten reports and a per-condition distribution of top-hypothesis scores on the 25-point, five-criterion rubric (Figure 10A). Mean top-hypothesis scores were statistically indistinguishable between the two case studies — CHO 22.3 ± 1.1 and iPSC 22.2 ± 1.1 (mean ± SD, max 25; Welch t = 0.13, n.s.). Both sit near 89% of the maximum, comparable to — though not exceeding — the normalized scores reached under the matched four-criterion harness of Section 3.5 (Sonnet 4.6 ≈ 92–93%, Opus 4.8 ≈ 87–90%; Figure 11A). This parity is notable because the ABCD-Stack is scored on a stricter five-criterion rubric that adds faithfulness to the brief, a criterion on which points can be lost that the four-criterion rubric never charged; maintaining a comparable normalized score under the harder rubric is therefore consistent with the enhanced pipeline producing hypotheses of at least equal quality. Every one of the ten runs cleared 20/25 (spread 20.5–23.5).

**Figure 10.**
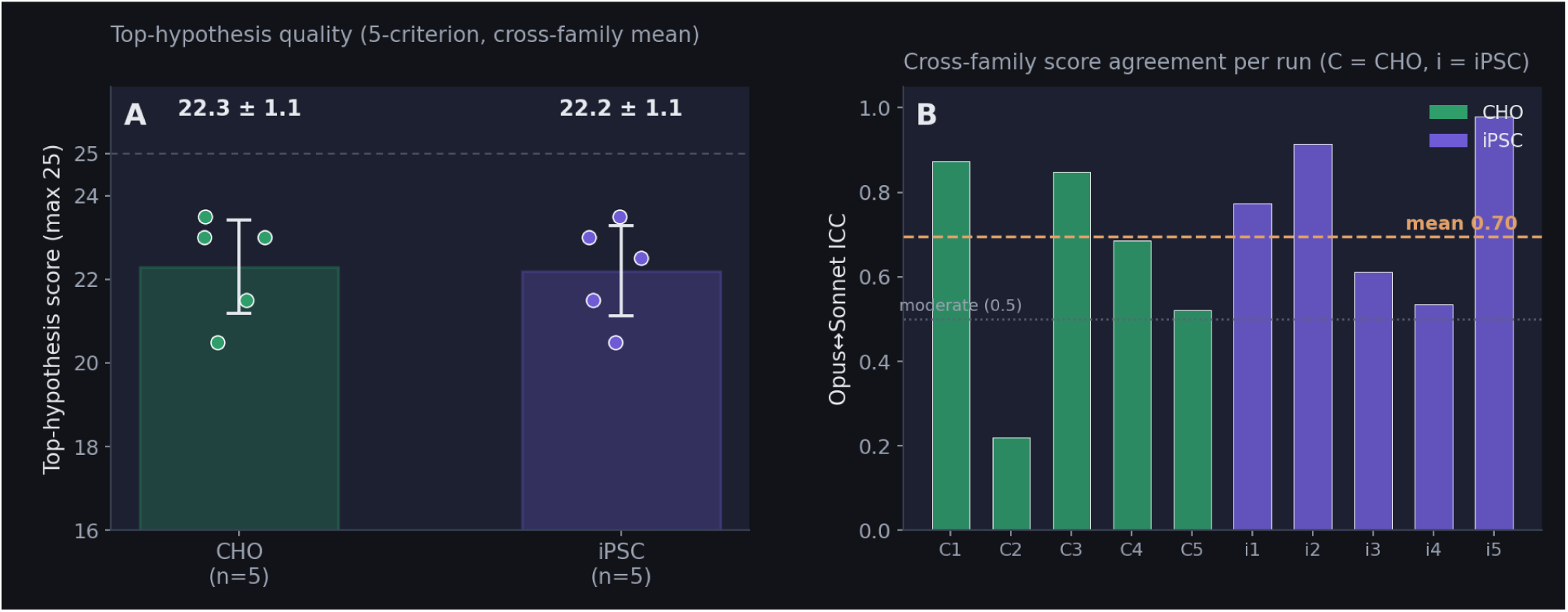
ABCD-Stack repeated-run results (five repeats per case). (A) Top-hypothesis score on the five-criterion rubric (max 25), shown as mean ± SD with individual runs overlaid; CHO and iPSC are statistically indistinguishable (22.3 ± 1.1 vs 22.2 ± 1.1; Welch t = 0.13, n.s.). (B) Cross-family score agreement per run, as the intraclass correlation between the Opus 4.8 and Sonnet 4.6 critics (C = CHO runs, i = iPSC runs); the mean ICC is 0.70, above the conventional moderate-agreement threshold of 0.5, but with wide run-to-run spread (0.22–0.98).

**Figure 11.**
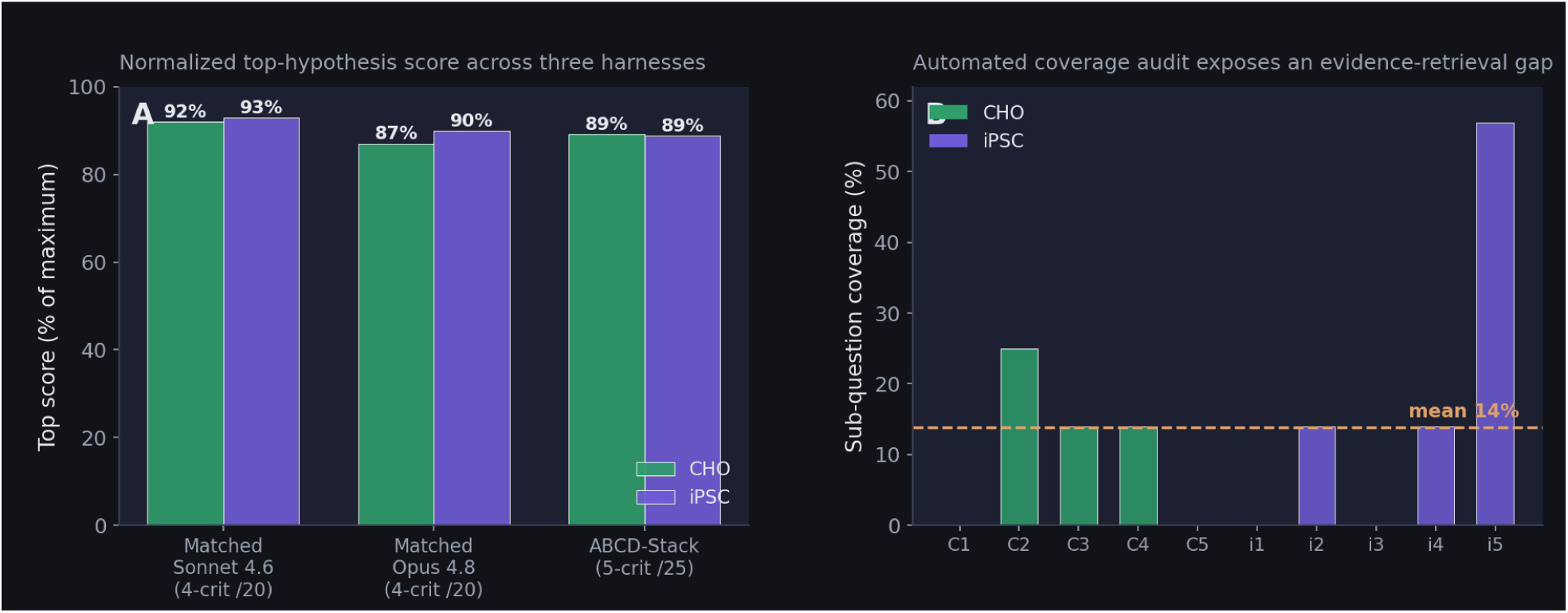
ABCD-Stack in context. (A) Normalized top-hypothesis score (percent of maximum) across three harnesses: matched Sonnet 4.6 and matched Opus 4.8 on the four-criterion /20 rubric (Section 3.5) and the ABCD-Stack on the five-criterion /25 rubric (this work). The ABCD-Stack reaches ∼89% of maximum on both case studies despite the stricter fifth criterion. (B) Per-run sub-question coverage from the automated audit (C = CHO, i = iPSC); coverage is low and variable (mean 14%), exposing a gap between the breadth of the decomposed brief and the facets a single five-hypothesis set addresses.

#### 3.6.3 Qualitative Shift in the Top Hypotheses

Beyond the score distribution, the content of the top-ranked hypotheses differs systematically from the four-criterion baseline of Section 3.5, and three changes recur across all ten runs (full records in Supplementary S6). First, every ABCD-Stack top hypothesis carries an explicit, quantitative falsification condition together with a named red-team revision, whereas the four-criterion records report a falsification field but no adversarial-revision step; the revision notes consistently show the Reviser narrowing scope before scoring — moving a causal test upfront, separating a confounder, or restricting the claim to a single organism (for example, the mouse-restriction and CRISPRi additions to the top iPSC hypothesis, S6.3). Second, the titles themselves more often encode a hedged, testable claim: six of the ten name a “contribution”, “conditional”, “causal-test”, or “limited” version, reflecting the faithfulness criterion and the adversarial loop penalising over-broad claims. Third, the mechanistic framing shifts from a single structural handle toward a causally testable pathway with built-in controls: where the matched Opus 4.8 four-criterion run converged on a local structural design (the BiP NBD–SBD linker variant, 17/20; Section 3.5, Supplementary S2.3), the ABCD-Stack CHO runs — driven by the identical CHO brief — instead foreground causal-chain interventions such as breaking a heavy-chain UPR–oxidative-stress feedback loop [61,64] (23.5/25, S6.1) or severing an AKT1–DNMT1–H3K9me3 silencing loop at the transgene locus (23.0/25, S6.2). A dimension-by-dimension contrast on the same CHO query is given in Table 7: it shows the shift from a single asserted structural mechanism to conditional, falsification-first claims with embedded controls, confirming that the change is driven by the harness rather than the query or the model tier. The net effect is a change in kind rather than only in number: the ABCD-Stack trades some of the single-model structural specificity for hypotheses that are explicitly falsifiable, scope-bounded, and causally framed — the properties the added harness stages were designed to enforce. Because these are the same ten runs whose scores appear in Figure 10, the shift is a property of the harness, not of a change in case study or model tier. Read across the case studies, the two configurations offer distinct advantages over an unaided investigator: the baseline already articulates hypotheses more explicitly than a typical single-pass human proposal, while the ABCD-Stack, by making the generate–critique–revise flow inspectable, converges on higher-confidence experimental hypotheses. In effect the pipeline convenes a team of reasoners that share the same evidence and logical rigour but adopt different search stances — a configuration a single investigator cannot occupy alone — and it is this constructed division of labour, rather than any one model, that yields hypotheses beyond single-investigator reach.

**Table 7.** Content contrast on the identical CHO brief between the baseline Opus 4.8 top hypothesis (four-criterion; Supplementary S2.3) and the five ABCD-Stack CHO runs (five-criterion; Supplementary S6). The ABCD-Stack column aggregates the five CHO runs; the per-top-hypothesis counts are descriptive summaries of the ten ABCD-Stack top hypotheses, not inferential statistics.

| Dimension | Baseline Opus 4.8 (four-criterion) | ABCD-Stack (five-criterion) |
| --- | --- | --- |
| Intervention target | A single structural handle — the BiP NBD–SBD chaperone linker variant (17/20) | A causal chain and its upstream node — e.g. heavy-chain UPR → oxidative-stress → copy-number loss (23.5/25); AKT1 → DNMT1 → H3K9me3 silencing loop (23.0/25) |
| Claim form | Asserted mechanism (“X accelerates chaperone release”) | Conditional, falsification-first (“if X contributes, then intervening on X improves Y”); 3.3 conditional markers per top hypothesis on average |
| Scope | Stated at full strength | Self-limited by the Reviser before scoring — “contribution”, “conditional”, “causal-test”, or “limited” versions in 6 of 10 titles |
| Experimental design | Mechanism-centred, controls implicit | Controls and confounder-separation embedded in the hypothesis (catalytic-dead controls, UCOE controls, copy-number correction); 1.4 explicit control mentions per top hypothesis on average |
| Adversarial revision | None (single-pass Critic) | Present in all ten runs (named red-team attack + Reviser repair before scoring) |

#### 3.6.4 Cross-Family Evaluation Reveals Uneven Critic Agreement

The paired-critic design turns the self-scoring confound of Section 3.5 into a measured quantity. Concretely, cross-family evaluation scores every hypothesis twice with two critics drawn from different model families — an Opus 4.8 critic and a Sonnet 4.6 critic — each applying the identical five-criterion rubric to the identical hypothesis text, and then quantifies how well their scores agree. A same-family critic, by contrast, evaluates output produced by its own family and tends to rate it within a narrow, generous band, so its scores confound the quality of the hypothesis with the shared inductive bias of the generating family; using a second family as an independent rater removes that shared bias from the agreement estimate.

Agreement is summarized per run as an intraclass correlation coefficient (ICC) over the two critics’ criterion scores, so that a run yields not only a hypothesis and its score but a number describing how much that score depends on which family did the scoring. Across the ten runs the Opus-versus-Sonnet ICC averaged 0.70 (SD 0.22), above the conventional 0.5 threshold for moderate agreement, but with a strikingly wide range — from 0.22 (CHO run 2) to 0.98 (iPSC run 5) (Figure 10B). Two observations follow. First, cross-family agreement is real but far from automatic: on most runs the two families rank the same hypotheses similarly, which is the precondition for treating either critic as a stand-in for the other, but the low-ICC runs show that agreement cannot be assumed on any single run. Second, the runs with the weakest agreement are also the ones where the two families would have designated different top hypotheses, which is exactly the situation a same-family critic would have hidden. The ICC is therefore not a nuisance statistic but a per-run reliability flag: a low value marks a report whose ranking is model-dependent and should not be acted upon without a human or third-family adjudication — the cross-family panel proposed as future work in Section 4.5.3, now instantiated and measured.

#### 3.6.5 The Coverage Audit Exposes an Evidence-Retrieval Gap

The automated coverage audit (Phase 2.6), added to catch the very confound that produced the CHO harness artifact of Section 3.5, reports an unexpected and important negative result. Keyword coverage of the generated hypotheses against the decomposed sub-questions was low and highly variable — CHO 11 ± 10% and iPSC 17 ± 21% (overall mean 14%), with three of ten runs at 0% and a single outlier at 57% (Figure 11B). This does not mean the hypotheses are off-brief; the faithfulness scores that the critics assigned were high, and the audit measures literal sub-question keyword overlap, a deliberately conservative proxy. What the low coverage flags is a gap between the breadth of the decomposed brief and the narrower slice of it that any single set of five hypotheses can address: the Decomposer routinely enumerates five to eight facets, but the synthesis stage, even with three personas, converges on the two or three most evidence-rich of them. We can only speculate as to why three personas do not widen coverage further, but three mechanisms are plausible and mutually reinforcing. First, the personas share a single evidence pool: all three read the same Phase-1 agent outputs, so a facet for which little evidence was retrieved is impoverished for every persona at once, and the personas differ in reasoning stance (integrative, structural, contrarian) rather than in which facets they attend to — they re-rank and re-frame the same well-evidenced facets instead of reaching for neglected ones. Second, the quality contract rewards depth over breadth: each hypothesis must carry a mechanism, a falsification condition, and a quantitative prediction, and those fields are far easier to satisfy for a facet backed by dense retrieved evidence, so all three personas are pulled toward the same evidence-rich core and away from thinly-supported facets that would score poorly on testability and consistency. Third, the downstream diversity selection (Phase 4) maximizes pairwise distance among the retained hypotheses but operates only on the candidates the personas actually produced; it can spread the final set across distinct mechanisms, but it cannot recover a sub-question that no persona chose to address. In short, persona diversity broadens how the same evidence is interpreted, not which parts of the brief get evidence — so coverage is bounded by retrieval breadth in Phase 1 rather than by generation diversity in Phase 2, which points to evidence retrieval, not synthesis, as the place to widen brief coverage in future work. The coverage audit thus earns its place not by confirming completeness but by quantifying incompleteness — surfacing, run by run, which facets of a multi-part brief went unaddressed, and providing the objective brief-coverage metric that Section 4.5.3 (Experiment 3) called for but had not yet operationalized.

## 4. Discussion

### 4.1 Cross-Domain Integration as the Core Driver of Novelty

The clearest signal in our results is where the strongest hypotheses come from. Across all four case studies, the highest-scoring and most actionable hypotheses are the ones that sit at the intersection of several evidence domains at once — a structural observation (an AlphaFold domain boundary or low-confidence linker), a network relation (a STRING two-hop path or a bottleneck node), a chemical handle (an ADMET-favourable scaffold), and a literature gap — rather than the ones grounded in any single source. The TrxR1/oxidative-GOF hypothesis for mutant p53 (Supplementary S3.1) is the archetype: it only becomes compelling when a chemistry observation (the APR-246 metabolite targets the thioredoxin system non-specifically [44]) is read together with a structural/regulatory one (the NRF2/SLC7A11 redox axis [42,43,45–47]) and a network one (a TSPYL5 stabilization route), so that four independent streams converge on a single redox switch.

This is the property the architecture was built to exploit. By giving four specialist agents non-overlapping tool subsets and forcing their outputs through a synthesizer that must cite an explicit evidence basis, the pipeline systematically manufactures the cross-domain collisions that a single-domain agent — or a single human working within one field — is unlikely to stage. The consistency criterion operationalizes this directly: it rewards hypotheses whose support converges across all four evidence agents and penalizes those resting on one or two, functioning as an automatic filter for the cross-domain integration that drives novelty. The remainder of the Discussion is organized around a single tension this raises — that the value of such a system depends far more on how it is prompted and contracted than on which model sits inside it (Section 4.2), a lesson we then turn into a concrete, better-engineered architecture (Sections 4.6).

It also bears repeating what these scores do and do not certify. A high Critic score means a hypothesis is well-formed — novel relative to the retrieved literature, mechanistically explicit, quantitatively predictive, internally consistent across evidence streams, and accompanied by a falsification condition — not that it is correct. The system’s contribution is to industrialise the production of such well-formed, falsifiable, cross-domain hypotheses and to make their quality machine-auditable; adjudicating their scientific truth remains a task for expert review and experimental test, which this paper scopes as future work (Section 4.5) rather than a claim of the present study.

### 4.2 The Prompting Harness, Not the Model Tier, Dominates Hypothesis Quality

The single most important lesson from the Sonnet-versus-Opus comparison is methodological rather than model-specific. In our initial, uncontrolled runs Opus 4.8 with Adaptive Thinking was expected to produce more structured, more numerous, and more reliable hypotheses, and at first glance seemed to do so(data not shown). A controlled re-test under a matched harness (Section 3.5) showed each of these apparent advantages to be an artifact of unequal prompts, output schemas, hypothesis counts, and query coverage rather than a property of the model. Three variables did all the work. The Opus prompt requested three hypotheses to Sonnet’s two; it alone mandated a structured quality schema (explicit falsification condition, quantitative prediction, prior-art differentiation); and the CHO query given to Opus carried an extra antibody-design sub-question. Because those were precisely the dimensions on which we had read “model differences” — output count, the presence of falsification/quantitative fields, and the CHO target framing — the comparison had conflated model with prompt. Once Sonnet 4.6 is given the same three-hypothesis count and the same mandatory schema, it emits the same structured fields, the same number of hypotheses, the same low run-to-run variance, and statistically equal or higher scores; the reliability edge we had attributed to Adaptive Thinking disappears with it.

The practical implication is that the largest, cheapest, and most reproducible lever on hypothesis quality in a system of this kind is the prompting harness — mandating falsification conditions and quantitative predictions, requesting an adequate and diverse set of candidates, ensuring the query enumerates every facet of the problem, and decoupling scoring from the generating model family — and this lever operates largely independently of the LLM tier. This is not a claim that model capability is irrelevant. Adaptive Thinking may well help on genuinely harder reasoning that these tasks did not expose, and a more capable model raises the ceiling a good harness can reach. It is a claim about attribution: on cross-domain hypothesis generation as evaluated here, no model-tier advantage is supportable unless the harness variables are held identical, runs are repeated, and — because each hypothesis is scored by a Critic of its own family — scoring is cross-checked across families. Every one of those requirements becomes an architectural commitment in the ABCD-Stack (Section 3.6), which is how we convert this diagnosis into a working system rather than a caveat.

The consistency criterion, once part of the unified schema, proved particularly valuable as an automatic filter for the cross-domain integration of Section 4.1. Hypotheses scoring 5/5 on consistency (e.g., the switch-II ON-state and TrxR1/GOF models) drew coherent support from all four evidence agents, whereas those scoring lower (e.g., the DBD cryptic-pocket hypothesis at 4/5) leaned on one or two sources. In the ABCD-Stack this filter is strengthened rather than replaced: consistency is retained and joined by a fifth criterion, faithfulness to the brief (Section 3.6), so that a fluently testable but off-brief hypothesis can no longer outscore an on-brief one — the specific rubric blind spot exposed in Section 3.5.

### 4.3 Comparison with Related Work

Our work sits at the intersection of three lines of prior research — tool-using scientific agents, LLM-based hypothesis generation, and the methodology of evaluating LLM systems — and its two contributions engage each differently. Along the first line, autonomous chemistry agents such as Coscientist [4] showed that an LLM can plan and execute laboratory procedures end to end, and tool-augmentation frameworks — ReAct [8], Toolformer [9], and multi-agent orchestration in AutoGen [14] — established the mechanics of interleaving reasoning with external calls. These systems are powerful but overwhelmingly single-domain: Coscientist reasons within chemistry, and drug-repurposing pipelines [33,66] typically query a static knowledge graph without dynamic literature integration. Along the second line, the most closely related prior work [32] demonstrated that an LLM can generate plausible hypotheses from literature alone, but did not integrate structural biology or interaction-network evidence and did not use a structured, tool-grounded multi-agent pipeline. Our architecture addresses these gaps by making dynamic literature retrieval [22], protein-interaction queries [18], AlphaFold structure analysis [15], and computational chemistry [29,30] accessible within a single inference run, and — as Section 4.1 argues — by rewarding hypotheses whose support converges across all four domains, so that cross-domain integration is an explicit selection objective rather than an incidental by-product.

The graph-based RAG approach [26] extends standard document-level RAG [7] to structured biological knowledge, enabling multi-hop reasoning over interaction networks that flat text retrieval cannot recover.

The paper’s headline result speaks to the third line — how LLM systems are compared — and here the relevant prior art is cautionary rather than architectural. A growing body of work shows that reported differences between models or prompting strategies are often dominated by evaluation confounds: sensitivity to prompt wording and output format, the number of samples drawn, and the choice of judge. Our controlled re-test (Section 3.5) is a concrete instance of this hazard in the scientific-discovery setting, where an apparent Opus-over-Sonnet advantage dissolved once prompt, schema, hypothesis count, and query coverage were matched. We are not aware of prior work that isolates the prompting harness as the dominant variable in multi-agent scientific hypothesis generation specifically, and we offer the matched-harness protocol — identical prompts and schema, repeated runs, and cross-family scoring — as a small, transferable methodological contribution for benchmarking agents of this kind.

Finally, the ABCD-Stack (Section 3.6) is best read not as a new mechanism but as a disciplined assembly of three established ideas, each bound to a phase that closes one diagnosed confound: diversity-first generation from parallel personas is a structured analogue of self-consistency sampling [69]; the adversarial red-team/reviser loop follows the multi-agent debate and self-refinement literature [70,71]; and cross-family evaluation operationalizes the LLM-as-a-judge paradigm [72] while directly targeting its known self-preference bias by requiring agreement between two independent model families and reporting that agreement as a per-run reliability statistic. What the ABCD-Stack adds is the integration: the pipeline reports its own coverage and agreement limits rather than emitting a single unqualified score.

### 4.4 Limitations

Several limitations of the current system should be noted; each motivates a specific validation experiment proposed in Section 4.5. First, the hypothesis evaluation is performed entirely by the same LLM family that generated the hypotheses, creating a potential systematic bias. Although the Hypothesis Critic node is prompted to provide adversarial evaluation (”devil’s advocate”), it cannot fully replicate independent expert peer review (addressed by the blinded expert evaluation in Section 4.5.1).

Second, the sequential execution architecture (four evidence agents run one after another within each query) requires 15–25 minutes per run. Parallelizing the evidence collection phase — which is architecturally feasible within the LangGraph framework using fanout patterns — would reduce latency to approximately the maximum single-agent execution time.

Third, a small number of citations retrieved by the Literature Scout during experimental runs were initially agent-reported and required verification: five such entries were flagged, and for this version each was manually checked against primary sources and, where the agent-reported details did not correspond to a real publication, replaced with the closest verified reference on the same topic. More generally, LLMs used in retrieval tasks may occasionally hallucinate bibliographic details even when using real-time search APIs, so agent-reported citations should always be confirmed against primary sources before publication.

Fourth, the system has not yet been evaluated at scale (a large number of independent queries with blinded expert scoring) or compared with hypothesis generation by human researchers given equivalent information access. Such controlled comparisons are necessary to establish whether the quality gains we observe translate to practical research acceleration, and are specified as concrete protocols in Sections 4.5.1 and 4.5.3. Fifth, the repeated-run analysis uses only five repeats per condition, so the significance test reported in Section 3.5.1 (the CHO Welch comparison, t = 2.89, df ≈ 8, p < 0.05) rests on a small sample and should be read as provisional; a larger number of repeats would be needed to estimate the effect size reliably, and the matched-harness conclusion we emphasize does not depend on that single comparison.

### 4.5 Proposed Validation Experiments and Future Directions

The hypotheses reported here are, by construction, experimentally falsifiable, but the present study establishes only their generation, not their correctness or their advantage over human-generated alternatives. The correctness of any individual hypothesis can ultimately be settled only at the bench, and two practical constraints make that settlement expensive. First, in a sufficiently specialized research area very few people can validly judge a hypothesis in advance: even the blinded expert evaluation we propose below (Section 4.5.1) is a proxy for correctness, not a substitute for it, and for the most novel cross-domain claims the pool of qualified judges is small and may itself disagree. Second, confirming each hypothesis by wet-lab experiment costs substantial time and money, so exhaustively testing every candidate is not realistic. Taken together, these constraints shift the value of the system away from producing a single best hypothesis and toward producing several falsifiable candidates and then identifying the most cost-effective one to test. This is precisely why every top-ranked hypothesis is emitted with an explicit falsification condition, a quantitative prediction, and its required reagents (Section 3.6, Supplementary S3): that structure lets the candidates be ranked by how cheap, fast, and decisive their key experiment would be, so that scarce wet-lab resources can be concentrated on the single most informative test rather than spent uniformly. Generating a diverse candidate set and then triaging it by experimental cost-effectiveness is therefore a central value proposition of the system, not an incidental feature. The nearest-term validation program then has two parts. The first is wet-laboratory testing of the generated mechanistic claims: because each top-ranked hypothesis already defines a minimal, decisive experiment, a practical near-term study would advance two or three of the highest-ranked, most cost-effective hypotheses to bench testing through an academic collaboration and report the single most informative statistic for the field — the fraction of AI-generated hypotheses that survive their own falsification test. We deliberately do not enumerate individual wet-lab protocols here, since each design follows mechanically from the hypothesis’s stated falsification condition and quantitative prediction. The second part evaluates the generation system itself — blinded expert scoring (Section 4.5.1), architectural ablations (Section 4.5.2), and the model-strategy divergence test (Section 4.5.3) — converting the qualitative quality signal observed here into controlled, blinded evidence of research acceleration.

#### 4.5.1 Blinded Expert Evaluation at Scale

The most important methodological limitation (Section 4.4) is that hypotheses were scored by the same LLM family that generated them. We propose a blinded evaluation in which a panel of domain experts (oncology, structural biology, bioprocess engineering) scores a corpus of ≥100 hypotheses, drawn from ≥30 independent research questions, on the same novelty/testability/impact/consistency rubric. Critically, the corpus should interleave (i) system-generated hypotheses, (ii) human-expert hypotheses, and (iii) recent genuine hypotheses extracted from the literature, all stripped of provenance and stylistically normalized so that AI origin cannot be inferred from surface form. Inter-rater reliability (Krippendorff’s α) and the agreement between expert and Critic scores would quantify how far the automated Critic can substitute for peer review, and would calibrate the scores reported here against an external standard.

#### 4.5.2 Architectural Ablation Studies

The matched-harness re-test (Section 3.5) already separated the model upgrade from the added quality/consistency schema, but the contribution of the individual evidence agents remains untested. We propose systematic ablations holding the query set fixed: (i) *evidence-agent leave-one-out* — removing each specialist agent (Literature, Network, Structure, Chemistry) in turn to quantify its marginal contribution to top-hypothesis quality and to confirm that cross-domain integration, rather than any single source, drives novelty (Section 4.1); (ii) *reasoning-mode ablation* — Opus 4.8 with and without Adaptive Thinking, and Sonnet 4.6 with and without the consistency criterion, to disentangle model capability from rubric design; and (iii) *evidence-order and parallelization* ablation — permuting agent execution order and comparing sequential versus fan-out evidence collection to verify that the sequential dependency is not silently biasing synthesis. Each ablation reuses the existing pipeline, requiring only configuration changes, and would produce the controlled attribution the current before/after comparison cannot.

#### 4.5.3 Validating the Model-Strategy Divergence and the Faithfulness Criterion

The qualitative coding of Section 3.5 yields two empirical claims that are themselves testable: (a) that Sonnet 4.6 and Opus 4.8 occupy complementary regions of hypothesis space (network-driven/integrative vs. structure-driven/local), and (b) that the current four-criterion rubric fails to reward faithfulness to the research brief. We propose four experiments, ordered by cost, that would confirm, quantify, or refute these claims. The first three reuse the existing pipeline with configuration changes only; the fourth requires wet-lab work and connects back to Section 4.5.

Experiment 1 — inter-coder reliability on the balanced corpus. The corpus coded here is already balanced (twelve top hypotheses per model, from five matched CHO and five matched iPSC repeats plus the single TP53 and KRAS runs). The natural extension is to code not only the top-ranked hypothesis but all three per run, and to have two or more blinded human coders assign five qualitative axis labels (intervention scale, design philosophy, primary evidence source, causal robustness, and faithfulness to the brief), so that inter-coder agreement (Cohen’s κ per axis) establishes whether the integrative-vs-local and substitution-vs-augmentation distinctions are reliably observable rather than artifacts of single-coder judgment. Pre-registered prediction: Opus 4.8 is coded local in significantly more cells than Sonnet 4.6 (one-sided proportion test, α = 0.05), and the CHO integrative/local divergence replicates across repetitions.

Experiment 2 — cross-model scoring to separate strategy from self-scoring bias. Because each hypothesis is currently scored by a Critic of its own family, the CHO score gap (Sonnet’s 19/20 CTCF–TAD top vs. Opus’s 17/20 local tops) confounds hypothesis quality with self-scoring leniency. We would re-score every hypothesis in the balanced corpus under a full cross design: each hypothesis evaluated by a Sonnet-4.6 Critic and an Opus-4.8 Critic, with generator identity hidden — the cross-family evaluation later operationalized in the ABCD-Stack (Section 3.6). The generator-vs-evaluator interaction term isolates self-scoring bias; the intraclass correlation between the two Critics quantifies how interchangeable they are. Pre-registered prediction: a positive same-family bias (a Critic scores its own family’s hypotheses higher), and a narrowing of the CHO gap when Sonnet’s hypotheses are scored by the Opus Critic.

Experiment 3 — adding and validating the faithfulness criterion. We would extend the rubric with a fifth criterion, faithfulness to the brief (does the hypothesis address the question’s explicit requirements?), and re-score the balanced corpus on all five criteria. The discriminative test is whether faithfulness separates hypotheses that the existing four criteria rank equally: specifically, whether the off-brief Opus CHO tops (BiP linker, heavy-chain rigidification, CH1 folding; records 5, 6, 9) drop relative to the on-brief Sonnet CHO tops (CTCF–TAD, BiP–TP53, RPL11; records 10, 13, 14) once faithfulness is scored, while their novelty/testability/impact/consistency scores are unchanged. As an objective anchor for the otherwise-subjective criterion, we would also compute an automated brief-coverage metric — embedding-similarity between each hypothesis and the explicit sub-requirements enumerated in the prompt (e.g., the CHO prompt’s three numbered items) — and test its correlation with the human faithfulness score.

Experiment 4 — the decisive wet-lab head-to-head. The strongest test of whether strategic divergence carries scientific consequence is to advance the two models’ top CHO hypotheses to the bench in parallel, since they make non-overlapping, individually falsifiable predictions on the same engineering goal. Sonnet 4.6’s CTCF–TAD insulator hypothesis (record 10, 19/20) predicts that a single-copy antibody cassette flanked by CTCF sites in a high-insulation TAD holds productivity drift below CV 5% over 60 passages, with copy-number/expression stability (Hi-C, ChIP-seq, qRT-PCR) as the decisive readout. Opus 4.8’s BiP-linker hypothesis (record 5, 17/20) predicts that a fast-cycling BiP NBD–SBD linker variant at least doubles specific productivity by pulse-chase, with a misfolding/aggregation control (SEC-HPLC) as the decisive guard. Running both in the same producer-line background, and reporting which prediction survives its own falsification condition, would convert the strategy-divergence claim from a coding observation into an experimentally grounded statement about which search strategy yields hypotheses that hold up — the single most informative outcome for choosing a generator model in practice.

### 4.6 From Model Selection to Harness Engineering: Design Principles and What the ABCD-Stack Confirms

If the prompting harness rather than the LLM’s reasoning capability is the dominant lever on hypothesis quality (Section 3.5), the most productive next step is not a larger model but a better-engineered agent. The matched-harness results point to four design principles that a next-generation system should make architectural rather than incidental: (i) mandatory falsification and quantitative-prediction fields as schema-enforced output contracts rather than optional prompt text, since requiring each hypothesis to state what would refute it gave the single largest reliability gain; (ii) the requested hypothesis count and an explicit diversity objective as controlled parameters, since output count was a pure harness variable that nonetheless changed what the system appeared to discover; (iii) automatic auditing of query coverage, since the CHO confound arose when one facet of the brief was enumerated for one configuration but not the other; and (iv) scoring decoupled from the generating model family, since every score here was self-assigned by a Critic of the Synthesizer’s own family. These principles motivate four concrete components — a contract-enforcing schema gate, an explicit diversity-promoting generator, a brief-decomposition and coverage auditor, and a cross-family evaluation panel — which we implemented together as the ABCD-Stack (Section 3.6) rather than leaving as recommendations.

Run on the two condition-matched case studies, the ABCD-Stack confirms the argument and sharpens it. Each principle produced its predicted effect: the schema-enforced contract (A-CGP P1) eliminated the truncated, field-incomplete hypotheses of the Sonnet 4.6 baseline without a model change, so that every one of the ten runs cleared 20/25 (Section 3.6.2); diversity-first generation from three personas (B-DFG) supplied the pool that greedy Jaccard selection pruned to five distinct hypotheses; adversarial refinement (C-ARL) required each hypothesis to survive a named fatal-flaw attack before scoring, and its effect is visible not only in the scores but in the content of the top hypotheses (Section 3.6.3): across all ten runs the Reviser narrowed each claim before it was scored — moving a causal test upfront, separating a confounder, or restricting scope to a single organism — so that the surviving hypotheses are explicitly falsifiable, scope-bounded, and causally framed rather than broad and structural. Cross-family evaluation (D-CFEC) converted the self-scoring confound from a caveat into a measured per-run statistic. We do not attribute the narrow score spread to any single one of these stages, since the repeated-run design does not isolate their individual contributions; the ablation of Section 4.5.2 is intended to do so. The harness-control thesis is thus not merely diagnostic but generative — the same analysis that dismantled the model comparison also specified an agent that works.

This qualitative shift is, we argue, the more important gain — more so than the modest change in score — because it makes the generated hypotheses better suited to their actual purpose. A hypothesis-generation system is useful only insofar as its outputs can be acted on, and the properties the ABCD-Stack adds map directly onto what makes a hypothesis actionable at the bench. First, because every candidate states an explicit falsification condition and quantitative prediction, it is testable by construction and can be triaged by the cost-effectiveness of its decisive experiment — the value proposition set out in Section 4.5. Second, because the Reviser narrows scope before scoring, the surviving claims are honest about their own boundary conditions (a specific subpopulation, a single organism, a contribution rather than a sole cause), which is exactly the information an experimentalist needs to avoid over-interpreting a lead. Third, because controls and confounder-separation are embedded in the hypothesis itself, the system hands the researcher not just a claim but the beginnings of an experimental design that can distinguish causation from correlation. On the identical CHO brief, the baseline Opus 4.8 converged on a single elegant structural handle, whereas the ABCD-Stack returned causally framed, self-limited, control-bearing hypotheses (Table 7); the former is arguably the better single idea, but the latter is the better raw material for a falsification-driven research program. In this sense the harness does not merely raise a score — it changes what the system is for, from proposing the most striking mechanism to proposing the most testable one.

Two instrumented phases return information no single-critic, single-synthesizer pipeline could. The cross-family ICC (mean 0.70, range 0.22–0.98) shows that agreement between model families is real on average but unreliable on any given run, so the blinded expert evaluation of Section 4.5.1 remains necessary precisely for the low-ICC cases. The coverage audit (mean 14%) quantifies that a five-hypothesis set addresses only a fraction of a brief the Decomposer expanded into five to eight facets; this is the point of the audit, not a failure of it, and it reframes a natural next objective as coverage-directed generation, in which unaddressed sub-questions are fed back as targeted prompts. Three limitations bound these results: the faithfulness criterion is scored but not yet validated against the independent coverage metric of Section 4.5.3; both critics remain Claude-family models, so cross-family here means cross-generation, not cross-vendor; and the runs still measure self-consistency, not scientific correctness. The decisive question — how many ABCD-Stack hypotheses survive their own falsification condition at the bench — is unchanged and remains the single most informative outstanding result. The ABCD-Stack closes the gap between the paper’s design recommendations and a working artifact; it does not close the gap between a well-engineered generator and a validated one, which the wet-lab and blinded-expert programs of Section 4.5 are designed to bridge.

## 5. Acknowledgments

This work was supported by the Ministry of Education, Culture, Sports, Science and Technology (MEXT), Japan, under the Grant No J261013007.

We thank the developers of LangChain/LangGraph [11,12], the STRING database team [18], the CTD team [19], the AlphaFold EBI team [15], and the BioMCP [27] and paper-search-mcp contributors for providing open-access tools and data that made this system possible.

## 7. AI Use Disclosure

This study concerns an AI system: the reported hypotheses were generated by large language models (Claude Sonnet 4.6 and Claude Opus 4.8) operating within the multi-agent pipeline described in Section 2. AI-based translation was used to render Japanese-language text into English, including the system prompts and hypothesis outputs reproduced in the Supplementary Information. During manuscript preparation, the authors also used Claude (Anthropic), an AI assistant, for document formatting, for cross-checking consistency between the main text and the Supplementary Information, and for verifying bibliographic references against their sources. The authors reviewed and edited all AI-assisted output, including all translations, and take full responsibility for the content of this manuscript. AI tools are not listed as authors.

