## Supplementary Information for "Harness engineering enables a multi-agent LLM system to generate cross-domain hypotheses beyond single-investigator reach"

This Supplementary Information provides the raw hypothesis data, the exact prompts, and the repeated-run measurements that support the consolidated results in the main text (Tables 5–7, Section 3). It is organized into six sections. Section S1 gives the full descriptions of the Sonnet 4.6 oncology hypotheses (main-text Table 5). Section S2 gives the full per-run descriptions of the Opus 4.8 hypotheses across all four case studies (main-text Tables 5–6). Section S3 reproduces, in the system's native field schema, the verbatim structured records emitted by the Hypothesis Synthesizer for the top-ranked hypotheses. Section S4 reproduces verbatim every system prompt and quality contract that constitutes the harness — the seven baseline node prompts (S4.1), the Synthesizer and Critic quality contracts (S4.2–S4.3), and the ABCD-Stack prompts including the Decomposer, the three synthesizer personas, the Red-Team, the Reviser, and the five-criterion cross-family Critic (S4.4) — so that every claim made about the prompt layer in Section 2.4.5 can be checked against the prompt itself. Section S5 provides the matched-harness repeated-run data underlying Section 3.5: the per-repeat top-hypothesis scores (S5.1), the integrative-versus-local strategy coding of each repeat (S5.2), and the data provenance (S5.3). Section S6 reproduces the top-ranked hypothesis of each of the ten ABCD-Stack runs underlying Section 3.6.2 on the five-criterion schema. Unless noted otherwise, oncology scores use the unified four-criterion Critic schema (novelty / testability / impact / consistency, max 20), while the ABCD-Stack cross-family Critic in S4.4 uses the five-criterion schema (adding faithfulness, max 25). Underlying reports: notebooks 08-2 (Sonnet 4.6), 09 (Opus 4.8), and 11 (ABCD-Stack).

### S1. Detailed Hypothesis Descriptions — Sonnet 4.6 (Oncology, Notebook 08-2)

#### S1.1 TP53-Mutant Solid Tumors (Sonnet 4.6)

**Top hypothesis — USP7–DNMT1/BRD4 Dual Epigenetic-Loop Disruption (score 18/20; novelty 4/5, testability 5/5, impact 4/5, consistency 5/5).** Under the unified max-20 harness, Sonnet 4.6’s top-ranked TP53 hypothesis proposes that TP53-mutant tumours sustain two mutually reinforcing epigenetic loops — USP7 hyperactivation stabilising DNMT1 to silence tumour-suppressor promoters, and GOF-mutant p53 co-occupying super-enhancers with BRD4 to over-transcribe oncogenic drivers — so that a single bifunctional molecule combining a USP7 catalytic inhibitor and a BRD4 bromodomain inhibitor would break both loops at once. Supporting evidence: the Network Analyst identified a USP7–DNMT1 (score 987)–UHRF1 (score 928) epigenetic triangle, and the Chemistry Opportunity agent identified GOF p53 (R248W/R273H)–BRD4 super-enhancer co-binding amenable to JQ1-class blockade. Consistency 5/5 (all four evidence streams converge); testability 5/5 (a PEG-linked P5091–JQ1 prototype is buildable from existing tool compounds and testable in R248W/R273H lines).

Falsification: if the bifunctional molecule fails to lower DNMT1 protein and BRD4-target transcription below single-agent controls in mutant lines, the hypothesis is rejected.

##### S1.2 KRAS G12C Pancreatic Cancer (Sonnet 4.6)

**Top hypothesis — IRS1–PTPN11 Pancreatic-Specific Resistance Axis (score 19/20; novelty 5/5, testability 5/5, impact 4/5, consistency 5/5).** Under the unified max-20 harness, Sonnet 4.6's top-ranked KRAS hypothesis proposes that the endocrine microenvironment of the pancreas drives sotorasib resistance in a tissue-specific way: insulin signalling through the IRS1–PTPN11 (SHP2) axis reactivates KRAS G12C, so that combining sotorasib with the SHP2 inhibitor TNO155 restores GDP-state trapping. Network analysis placed PTPN11 as a hub linking IRS1 (score 982), SOS1 (score 995) and PIK3CB (score 999), and pancreatic-cancer enrichment flagged endocrine resistance ($p = 1.33\times10^{-24}$), tying the insulin pathway to KRAS reactivation in a single connector. Consistency 5/5; testability 5/5 (immediately testable in MIA PaCa-2 under high/low insulin with sotorasib ± TNO155 and IRS1 knockdown rescue). Falsification: if SHP2 inhibition does not lower KRAS-GTP and restore sotorasib sensitivity under high-insulin conditions, the hypothesis is rejected.

#### S2. Detailed Hypothesis Descriptions — Opus 4.8 (All Case Studies, Notebook 09)

##### S2.1 TP53-Mutant Solid Tumors (Opus 4.8)

Three hypotheses were generated and ranked (Figure 6):

| Rank | Hypothesis | N | T | I | C | Total |
|---|---|---|---|---|---|---|
| 1 ★ | Oxidative Stress-Dependent GOF & TrxR1 Vulnerability | 5 | 5 | 4 | 5 | **19/20** |
| 2 | TSPYL5 Inhibition — Pancreatic Cancer-Specific p53 Degradation | 5 | 4 | 5 | 4 | **18/20** |
| 3 | DBD Cryptic Pocket Pan-Mutant Allosteric Stabilization | 4 | 4 | 5 | 4 | **17/20** |

N = Novelty, T = Testability, I = Impact, C = Consistency; ★ = recommended by Hypothesis Critic.

**Top hypothesis — Oxidative Stress-Dependent GOF & TrxR1 Vulnerability (19/20).** The Synthesizer integrated four independent evidence streams: (1) Literature Scout — GOF p53 phenotypes are stress-conditional rather than constitutive; (2) Chemistry Opportunity — APR-246 active metabolite (methylene quinuclidinone, MQ) targets TrxR1/glutathione non-specifically rather than p53 directly [36,40]; (3) Structure Function — NRF2/SLC7A11 axis activation by mutant p53 under oxidative conditions [43]; (4) Network Analyst — TSPYL5 as an MDM2-independent p53 stabilizer. The hypothesis proposes that TrxR1 inhibition selectively collapses GOF-dependent transcription under oxidative conditions, providing mutant-p53-selective killing. Consistency 5/5 (all four evidence agents converge on oxidative redox as the common switch); Testability 5/5 (immediately testable in R175H/R273H isogenic cell lines). Falsification condition: if GOF transcriptional targets (MYC, NF-κB) are not suppressed in TrxR1-knockdown cells under oxidative stress, the hypothesis is rejected.

**Second hypothesis — TSPYL5 Inhibition (18/20).** TSPYL5 was identified as a USP7-independent p53 stabilizer whose expression is amplified by KRAS signaling in pancreatic cancer, positioning it as a KRAS-p53 bridging factor. TSPYL5 knockout is predicted to selectively degrade mutant p53 (half-life reduction >50%) in KRAS-mutant but not KRAS-wild-type cells.

**Third hypothesis — DBD Cryptic Pocket Stabilization (17/20).** Transient cavities form within the β-sandwich core of the DNA-binding domain (pLDDT 88.4) [15] upon GOF misfolding; compounds occupying these pockets could pan-stabilize multiple structural mutation subtypes (R175H, R248Q) [36,40]. Quantitative prediction: lead compound raises Tm ≥ 2°C for both R175H and R248Q, restores ≥50% wild-type transcriptional activity ($IC_{50}$ < 5 μM).

#### S2.2 KRAS G12C Pancreatic Cancer (Opus 4.8)

Rank | Hypothesis | N | T | I | C | Total |

|——|————|—|—|—|—|——-|

1 ★ | switch-II Flexibility & Pre-emptive ON-state Inhibition | 4
5 | 4 | 5 | **18/20** |

2 | Wnt-YAP1 Non-MAPK Axis Latent Resistance | 5 | 4 | 4 | 4 |

**17/20** |

3 | Stromal GF-Driven GTP Reloading + Macropinocytosis Dual Block |

3 | 4 | 5 | 4 | **16/20** |

**Top hypothesis — switch-II Flexibility & Pre-emptive ON-state Inhibition (18/20).** Switch-II (residues 59–75, AlphaFold pLDDT 73–79) is intrinsically disordered [15], and GF stimulation from the pancreatic stroma can shift the GDP/GTP conformational equilibrium toward the ON-state without requiring secondary mutations (Y96D, G13D) [50,52]. This mutation-free resistance mechanism is invisible to sequencing-based monitoring. Pre-emptive use of RMC-6236 (pan-RAS(ON) macrocycle) [53] is proposed to capture residual KRAS activity before Y96D clonal expansion. Quantitative prediction: ON-state KRAS-GTP occupancy ≥2-fold in mutation-negative resistant cells; RMC-6236 pERK $IC_{50}$ < 100 nM with ≥80% growth inhibition.

**Second hypothesis — Wnt-YAP1 Non-MAPK Axis (17/20).** Network Analyst detected a KRAS→CTNNB1(β-catenin)→YAP1 two-hop path [18] not captured by standard MAPK/PI3K feedback monitoring. Sotorasib-driven MAPK blockade activates this ERK/AKT-phosphorylation-negative transcriptional survival axis. TEAD inhibitor co-treatment is predicted to eliminate >70% of residual colonies in sotorasib-resistant PDAC organoids.

#### S2.3 Antibody Drug Manufacturing — CHO Cell Engineering (Opus 4.8)

For the CHO case, the latest condition-matched run (notebook 09, Opus 4.8 with Adaptive Thinking) generated and ranked three hypotheses on the unified 4-criterion schema (novelty / testability / impact / consistency, max 20).

| Rank | Hypothesis | N | T | I | C | Total |
|---|---|---|---|---|---|---|
| 1 ★ | BiP NBD–SBD Linker Variant for Accelerated Chaperone Release | 4 | 5 | 4 | 4 | **17/20** |
| 2 | FUT8 Stem-Region Engineering for Homogeneous Low-Fucose Glycosylation | 4 | 5 | 4 | 4 | **17/20** |
| 3 | IDR Rigidification (XBP1s/ATF6) for Reduced Transcriptional-Noise Lines | 5 | 3 | 5 | 3 | **16/20** |

N = Novelty, T = Testability, I = Impact, C = Consistency; ★ = recommended by Hypothesis Critic. (Hypotheses 1 and 2 tied at 17/20; the Critic recommended hypothesis 1 for its mechanistic clarity and broad applicability to secretion flux.)

**Top hypothesis — BiP NBD–SBD Linker Variant for Accelerated Chaperone Release (17/20).** Structure analysis placed BiP/HSPA5 (pLDDT 90.0) as a high-confidence chaperone whose nucleotide-binding domain (NBD, 28–405) and substrate-binding domain (SBD, 420–629) are joined by a low-confidence linker (522–531) that acts as the hinge of ATP/ADP-dependent allostery. The hypothesis proposes engineering this linker with disease-derived fast-cycling point mutations to accelerate ADP/ATP exchange, speeding substrate release so that excess retention of immature antibody is relieved and secretion flux increases. The mechanism reframes the problem away from chaperone over-expression (which can itself suppress secretion) toward tuning the cycling rate of an existing chaperone. Quantitative prediction: specific productivity (qP) at least doubles and ER retention of immature antibody falls by ≥50%. Proposed experiment: introduce the fast-cycling BiP-linker variant into CHO and measure antibody secretion rate by pulse-chase. Falsification: if the fast-cycling variant shows secretion at or below wild-type, the hypothesis is rejected.

**Second hypothesis — FUT8 Stem-Region Engineering for Homogeneous Low-Fucose Glycosylation (17/20).** Rather than fully knocking out FUT8 (catalytic-domain pLDDT 92), the hypothesis proposes modifying the disordered stem region (67–106) that governs Golgi localization and oligomerization, weakening localization so that the enzyme retains catalytic activity but spends less time in the Golgi, lowering core fucosylation partially and uniformly. Quantitative prediction: core-fucose held to a narrow 5–15% window (CV < 3%), ADCC roughly doubled, and growth rate retained at ~1.5× that of a full-knockout line. The structural rationale is that the highly confident catalytic domain has dominated producer-line design while the disordered stem — proposed here as a localization/oligomerization control element — has been overlooked. Proposed experiment: compare the glycan profile and ADCC of a stem-localization-motif-modified CHO line against a full FUT8 knockout.

A third hypothesis — IDR rigidification of XBP1s/ATF6 for reduced transcriptional-noise lines (16/20) — proposed rigidifying stochastic phosphorylation sites within the intrinsically disordered regions of XBP1s (mean pLDDT 68.3) and ATF6 (55.2) by alanine substitution, predicting that lot-to-lot productivity CV would halve (from 15–20% to 7–10%) while mean titer is retained at ≥1.3×. With hypotheses 1 and 2 tied at 17/20, the Critic recommended the BiP-linker variant for the established BiP allostery it builds on and the broad, secretion-flux-level applicability of the intervention; it flagged the principal risk that faster release could increase misfolded-antibody secretion (devil's advocate), defining the misfolding/aggregation readout as the decisive control.

#### S2.4 iPSC Reprogramming — Yamanaka-Factor Substitutes (Opus 4.8)

A fourth case study asked the system to propose mechanisms for small/medium-molecule or peptide substitutes for the Yamanaka factors used to generate iPSCs. In the condition-matched run, the Synthesizer/Critic generated and ranked three hypotheses on the unified 4-criterion schema (max 20).

| Rank | Hypothesis | N | T | I | C | Total |
|---|---|---|---|---|---|---|
| 1 ★ | LSD1/KDM1A Biased Dual-Switch (KLF4-Free Reprogramming) | 5 | 5 | 4 | 4 | **18/20** |
| 2 | p300 Metabolic-Acylation Glue (OCT4 Substitution) | 5 | 4 | 5 | 4 | **18/20** |
| 3 | 2-OG Dioxygenase Agonist (c-MYC Substitution) | 5 | 4 | 4 | 4 | **17/20** |

N = Novelty, T = Testability, I = Impact, C = Consistency; ★ = recommended by Hypothesis Critic. (Hypotheses 1 and 2 tied at 18/20; the Critic recommended hypothesis 1 for its approved-drug starting point and one-hop evidence.)

**Top hypothesis — LSD1/KDM1A Biased Dual-Switch for KLF4-Free Reprogramming (18/20).** A CoREST-dependent, biased LSD1 modulator is proposed to simultaneously activate SOX2 (via demethylation) and lift the p53 barrier (via p53 demethylation), making KLF4 dispensable. Mechanistically, KDM1A/LSD1 demethylates SOX2 Lys42/117 to activate it and p53 Lys370 to suppress its barrier function; an allosteric inhibitor that spares H3K4me erasure but biases substrate selectivity toward p53/SOX2 induces a reprogramming-permissive state while keeping p53 as a promoting factor. Supporting evidence: KDM1A–SOX2 are a one-hop direct link and KDM1A–NuRD–p53 converge; the approved drug tranylcypromine (predicted BBB 0.987, DILI 0.037) was the sole KDM1A hit, giving an approved-drug starting point. Quantitative prediction: with KLF4 omitted, iPSC efficiency is maintained at ≥0.7× the KLF4-

containing condition and p53 Lys370 demethylation increases ≥2-fold. Proposed experiment: add a CoREST-binding tranylcypromine derivative to a KLF4-free cocktail and quantify colony formation and p53/SOX2 methylation by ChIP. Falsification: if a biased inhibitor fails to reduce efficiency in p53-deficient cells and no SOX2 demethylation is detected, the hypothesis is rejected.

**Second hypothesis — p300 Metabolic-Acylation Glue for OCT4 Substitution (18/20).** Glycolysis-derived lactyl/crotonyl-CoA donors are proposed to drive p300-mediated non-acetyl acylation of pluripotency enhancers and IDR phase separation, substituting for OCT4 function. Mechanistically, p300 uses lactyl/crotonyl-CoA as substrates to modify pluripotency loci and activate enhancers; altered lysine modification within OCT4/SOX2 IDRs promotes phase-separation hub formation that boots the endogenous pluripotency loop. Supporting evidence: p300 is a lactylation/crotonylation enzyme and HIF1A–MYC–p300 converge at one hop; 60–88% of the Yamanaka factors are IDR-rich (LLPS-dependent), and metabolic acyl donors are an unexplored chemical space. Quantitative prediction: donor addition raises H3K18la/H3K9cr ≥3-fold and restores iPSC efficiency ≥2-fold under low-OCT4 conditions. Proposed experiment: add a cell-permeable crotonate ester to a low-OCT4 cocktail and measure lactylation/crotonylation-specific ChIP-seq and colony counts.

A third hypothesis — a 2-OG dioxygenase agonist for c-MYC substitution (17/20) — proposed metal-chelating mid-size molecules that actively accelerate TET/JmjC demethylation to substitute for c-MYC's metabolic–epigenomic function (predicted ≥1.5-fold 5hmC increase; efficiency ≥0.8× the c-MYC-containing condition). With hypotheses 1 and 2 tied at 18/20, the Critic recommended the LSD1 biased dual-switch for

#### S3. Verbatim Structured Records

This section reproduces, in full, the original structured records emitted by the Hypothesis Synthesizer for the seven top-ranked hypotheses across the four Opus 4.8 case studies (Section 3.4), as designated by the Hypothesis Critic. Each record is given in the system's native field schema (\`title\`, \`statement\` in if…then form, \`mechanism\`, \`evidence_basis\`, \`falsification\`, \`differentiation\`, \`quantitative_prediction\`, \`experiment\`, and the Critic-assigned scores). Field text is reproduced as generated, with only minor typographical normalization (Unicode subscripts, en-dashes); no mechanistic content was altered. These records are the primary experimental output of the system and the direct input to the validation program of Section 4.5.

##### S3.1 TP53 — Oxidative Stress-Dependent GOF & TrxR1

Vulnerability (Opus 4.8; 19/20) ★

**title:** Oxidative Stress-Dependent Gain-of-Function and TrxR1 Vulnerability in Mutant p53

**statement:** IF gain-of-function (GOF) mutant p53 (e.g., R175H, R273H) drives its oncogenic transcriptional program only under elevated oxidative stress, THEN selective inhibition of thioredoxin reductase 1 (TrxR1) will collapse GOF-dependent transcription and produce mutant-p53-selective cytotoxicity, while sparing TP53-null cells.

**mechanism:** GOF p53 phenotypes are stress-conditional rather than constitutive. The APR-246 active metabolite methylene quinuclidinone (MQ) targets the TrxR1/glutathione redox system non-specifically rather than refolding p53 directly [36,40]; mutant p53 activates the NRF2/SLC7A11 antioxidant axis under oxidative conditions [43]. TrxR1 inhibition removes the redox buffering on which GOF transcription depends, selectively disabling the oncogenic program. TSPYL5 acts as an MDM2-independent stabilizer that keeps mutant p53 protein abundant, ensuring the GOF program is poised to fire once redox conditions permit.

**evidence_basis:** Literature Scout: GOF p53 is stress-conditional. Chemistry Opportunity: MQ targets TrxR1/glutathione non-specifically [36,40]. Structure Function: NRF2/SLC7A11 axis activation by mutant p53 under oxidative stress [43]. Network Analyst: TSPYL5 as an MDM2-independent p53 stabilizer. All four streams converge on oxidative redox as the shared switch (Consistency 5/5).

**falsification:** If GOF transcriptional targets (e.g., MYC, NF-κB) are NOT suppressed in TrxR1-knockdown cells under oxidative stress, the hypothesis is rejected.

**differentiation:** Departs from the prevailing APR-246 "p53-refolding" mechanism-of-action narrative by reframing efficacy as redox-dependent collapse of a stress-conditional GOF program, predicting selectivity tied to oxidative state rather than to mutation subtype.

**quantitative_prediction:** ≥50% suppression of GOF target transcription and selective loss of viability in GOF lines (R175H/R273H) but not in TP53-null lines, under oxidative challenge.

**experiment:** Isogenic R175H/R273H vs. TP53-null panels (e.g., H1299 reconstitution). Cross TrxR1 perturbation (siRNA/shRNA; auranofin) with oxidative challenge ($H_2O_2$ or erastin); read out GOF target transcription (RT-qPCR/RNA-seq) and mutant-p53-selective viability.

**scores:** Novelty 5/5; Testability 5/5; Impact 4/5; Consistency 5/5; Total 19/20 (★ top-ranked).

#### S3.2 TP53 — TSPYL5 Inhibition for Pancreatic Cancer-Specific

p53 Degradation (Opus 4.8; 18/20)

**title:** TSPYL5 as a KRAS-p53 Bridging Factor Enabling Pancreatic Cancer-Specific Mutant p53 Degradation

**statement:** IF TSPYL5 is a USP7-independent stabilizer of mutant p53 whose expression is amplified by KRAS signaling, THEN TSPYL5 knockout will selectively degrade mutant p53 in KRAS-mutant but not KRAS-wild-type cells.

**mechanism:** TSPYL5 was identified as a USP7-independent p53 stabilizer positioned as a KRAS-p53 bridging factor; its expression is amplified by oncogenic KRAS signaling in pancreatic cancer, coupling KRAS activity to mutant p53 protein stability.

**evidence_basis:** Network Analyst: TSPYL5 as MDM2/USP7-independent p53 stabilizer; KRAS-driven expression linkage in PDAC. Consistency 4/5.

**falsification:** If TSPYL5 knockout does not reduce mutant p53 half-life selectively in KRAS-mutant cells (no differential vs. KRAS-wild-type), the hypothesis is rejected.

**differentiation:** Targets a KRAS-conditional, USP7-independent stabilization route rather than the canonical MDM2/MDM4 axis, predicting context-selective (KRAS-mutant) mutant-p53 degradation.

**quantitative_prediction:** Mutant p53 half-life reduction >50% upon TSPYL5 knockout in KRAS-mutant cells, with no comparable reduction in KRAS-wild-type cells.

**experiment:** TSPYL5 knockout/knockdown in matched KRAS-mutant vs. KRAS-wild-type lines; cycloheximide-chase half-life measurement of mutant p53; viability readout.

**scores:** Novelty 5/5; Testability 4/5; Impact 5/5; Consistency 4/5; Total 18/20.

##### S3.3 TP53 — DBD Cryptic Pocket Pan-Mutant Allosteric

Stabilization (Opus 4.8; 17/20)

**title:** Cryptic β-Sandwich Pocket in the p53 DNA-Binding Domain Enabling Pan-Mutant Allosteric Stabilization

**statement:** IF transient cavities form within the β-sandwich core of the p53 DNA-binding domain (DBD) upon GOF misfolding, THEN compounds occupying these cryptic pockets will pan-stabilize multiple structural mutation subtypes (e.g., R175H, R248Q) and restore wild-type-like transcriptional activity.

**mechanism:** AlphaFold analysis of the DBD (residues 100–290, pLDDT 88.4) [15] indicates a folded β-sandwich core within which transient cavities open upon destabilizing GOF mutations; a small molecule occupying such a cryptic pocket would act as a thermodynamic chaperone, raising melting temperature and restoring native fold across structurally distinct mutants.

**evidence_basis:** Structure Function: high-confidence DBD fold (pLDDT 88.4) [15]; Chemistry Opportunity: precedent for pharmacological chaperone stabilization of mutant p53 [36,40]. Consistency 4/5.

**falsification:** If a lead compound fails to raise Tm for both R175H and R248Q (no thermal stabilization), the pan-mutant stabilization hypothesis is rejected.

**differentiation:** Targets a single cryptic allosteric pocket common to multiple mutants rather than mutation-specific binders, aiming for pan-mutant rather than allele-specific reactivation.

**quantitative_prediction:** Lead compound raises Tm $\geq$ 2°C for both R175H and R248Q and restores $\geq$50% wild-type transcriptional activity ($IC_{50}$ < 5 μM).

**experiment:** Cryptic-pocket detection by MD/mixed-solvent simulation on the AlphaFold DBD; differential scanning fluorimetry (Tm) on purified R175H/R248Q; cell-based wild-type-activity reporter.

**scores:** Novelty 4/5; Testability 4/5; Impact 5/5; Consistency 4/5; Total 17/20.

##### S3.4 KRAS G12C — Switch-II Flexibility & Pre-emptive ON-state

Inhibition (Opus 4.8; 18/20) ★

**title:** Mutation-Free Sotorasib Resistance via Switch-II Conformational Flexibility, Reversible by Pre-emptive pan-RAS(ON) Inhibition

**statement:** IF growth-factor stimulation from the pancreatic stroma shifts the KRAS G12C GDP/GTP equilibrium toward the ON-state without requiring secondary mutations, THEN a mutation-free, sequencing-invisible resistance state will arise, and pre-emptive pan-RAS(ON) inhibition (RMC-6236) will suppress resistant-colony outgrowth before Y96D clonal expansion.

**mechanism:** Switch-II (residues 59–75; AlphaFold pLDDT 73–79) is intrinsically disordered [15]. Stromal growth-factor signaling drives GTP reloading, populating the active ON-state conformation independently of covalent-pocket mutations (Y96D, G13D) [50,52]. Because sotorasib engages only the inactive GDP-bound SII-P, ON-state KRAS escapes inhibition; this resistance is invisible to sequencing. A pan-RAS(ON) macrocycle (RMC-6236) [53] captures residual ON-state KRAS, pre-empting selection of resistant clones.

**evidence_basis:** Structure Function: disordered Switch-II (pLDDT 73–79) [15]. Literature Scout: stromal GF-driven, mutation-independent resistance [50,52]. Chemistry Opportunity: pan-RAS(ON) macrocycle RMC-6236 [53]. Network Analyst: MAPK/PI3K reactivation context. Consistency 5/5.

**falsification:** If sequencing-negative sotorasib-resistant cells do NOT show ≥2-fold ON-state KRAS-GTP occupancy (no mutation-free ON-state accumulation), the hypothesis is rejected.

**differentiation:** Identifies a non-genetic, conformational resistance mechanism distinct from the established Y96D/G13D mutational-resistance paradigm, and proposes pre-emptive rather than reactive ON-state inhibition.

**quantitative_prediction:** ≥2-fold ON-state KRAS-GTP occupancy in mutation-negative resistant cells; RMC-6236 pERK $IC_{50} < 100$ nM with ≥80% growth inhibition.

**experiment:** Sotorasib-resistant KRAS G12C lines (e.g., MIA PaCa-2, NCI-H358 derivatives): RAF-RBD pulldown of active KRAS-GTP with targeted sequencing to confirm absence of Y96D/G13D; pre-emptive RMC-6236 treatment scored for suppression of resistant-colony outgrowth.

**scores:** Novelty 4/5; Testability 5/5; Impact 4/5; Consistency 5/5; Total 18/20 (★ top-ranked).

#### S3.5 CHO Engineering — BiP NBD–SBD Linker Variant for

Accelerated Chaperone Release (Opus 4.8; 17/20) ★

**title:** Accelerating Chaperone Release by Engineering the BiP NBD–SBD Flexible Linker to Increase Antibody Secretion

**statement:** IF the BiP NBD–SBD flexible linker (522–531) is modified with disease-derived fast-cycling mutations, THEN over-capture is avoided and secretion rate doubles.

**mechanism:** BiP (pLDDT 90.0) joins its NBD and SBD through a low-confidence linker (522–531) that is the hinge of ATP/ADP-dependent allostery. Point mutations that accelerate ADP/ATP exchange at this linker speed substrate release, relieving over-retention of immature antibody and increasing secretion flux.

**evidence_basis:** Structure analysis placed the high-confidence NBD (28–405) and SBD (420–629) joined by the low-pLDDT linker 522–531 and identified it as the hinge of allosteric transmission; it provided a structural basis for excess chaperone causing secretion suppression.

**falsification:** If the fast-cycling linker-variant line shows secretion at or below wild-type, the hypothesis is rejected.

**differentiation:** Unlike BiP over-expression, it optimizes the cycling rate of the linker by mutation to avoid over-capture — an essentially different intervention.

**quantitative_prediction:** Specific productivity (qP) at least doubles; ER retention of immature antibody falls by ≥50%.

**experiment:** Introduce the fast-cycling BiP-linker variant into CHO and measure antibody secretion rate by pulse-chase.

**strength:** Built on established BiP allostery, the verification experiment is immediately executable and the mechanism is clear.

**weakness:** Accelerated release could increase folding failure and lead to secretion of defective antibody.

**devil_advocate:** Faster BiP cycling could shorten substrate-binding time and instead increase misfolded aggregation.

**scores:** Novelty 4/5; Testability 5/5; Impact 4/5; Consistency 4/5; Total 17/20 (★ top-ranked).

##### S3.6 CHO Engineering — FUT8 Stem-Region Engineering for

Homogeneous Low-Fucose Glycosylation (Opus 4.8; 17/20)

**title:** Stabilizing Homogeneous Low-Fucose Glycans by Engineering the FUT8 Stem Region without Full Knockout

**statement:** IF the Golgi-localization motif in the FUT8 flexible stem region (67–106) is modified, THEN homogeneous low-fucose glycans can be stabilized without full knockout.

**mechanism:** The low-confidence stem (67–106) of FUT8 (catalytic pLDDT 92) controls Golgi localization and dimerization. Replacing this stem with a localization-weakening variant shortens the enzyme’s Golgi residence while retaining catalytic activity, lowering core fucosylation partially and uniformly.

**evidence_basis:** Structure analysis showed the catalytic domain is highly confident while the stem (67–106) is disordered, suggesting an unresolved function controlling Golgi localization and oligomerization; producer-line design has focused only on the catalytic domain.

**falsification:** If the stem-modified line shows fucosylation equal to wild-type, or growth drops to full-knockout levels, the hypothesis is rejected.

**differentiation:** Unlike conventional full FUT8 knockout, it achieves graded, uniform defucosylation by non-catalytic stem localization control.

**quantitative_prediction:** Core-fucose held in a narrow 5–15% window (CV < 3%); ADCC roughly doubled; growth rate retained at ~1.5× the knockout line.

**experiment:** Compare the glycan profile and ADCC of a FUT8 stem-localization-motif-modified CHO line against a full knockout.

**strength:** Directly addresses the practical need for graded fucose control, and is easy to verify with standard methods.

**weakness:** The stem localization-control function is hypothetical; there is no guarantee modification affects residence time.

**devil_advocate:** Localization weakening could cause all-or-none loss of localization, giving knockout-like results rather than uniform reduction.

**scores:** Novelty 4/5; Testability 5/5; Impact 4/5; Consistency 4/5; Total 17/20 (tied with 3.7.5).

#### S3.7 iPSC — LSD1/KDM1A Biased Dual-Switch for KLF4-Free

Reprogramming (Opus 4.8; 18/20) ★

**title:** A CoREST-Dependent Biased LSD1 Modulator that Couples SOX2 Activation and p53-Barrier Relief to Make KLF4 Dispensable

**statement:** IF a CoREST-dependent biased LSD1 modulator handles both SOX2 demethylation/activation and p53-barrier relief, THEN KLF4 introduction becomes unnecessary.

**mechanism:** KDM1A/LSD1 has a dual nature: it demethylates SOX2 Lys42/117 to activate it and p53 Lys370 to suppress its barrier function. An allosteric inhibitor that preserves H3K4me erasure but biases substrate selectivity toward p53/SOX2 induces a reprogramming-permissive state while keeping p53 as a promoting factor.

**evidence_basis:** KDM1A–SOX2 is a one-hop direct link and KDM1A–NuRD–p53 converge; the approved drug tranylcypromine (BBB 0.987, DILI 0.037) was the sole KDM1A hit.

**falsification:** If a biased inhibitor does not reduce efficiency in p53-deficient cells and no SOX2 demethylation is detected, the hypothesis is rejected.

**differentiation:** Rather than uniform catalytic inhibition, it targets KLF4 replacement via substrate-selective biased design.

**quantitative_prediction:** With KLF4 omitted, iPSC efficiency maintained at ≥0.7× the KLF4-containing condition; p53 Lys370 demethylation increased ≥2-fold.

**experiment:** Add a CoREST-binding tranylcypromine derivative to a KLF4-free cocktail and quantify colony formation and p53/SOX2 methylation by ChIP.

**strength:** An approved-drug basis and one-hop direct evidence give outstanding immediate testability.

**weakness:** The chemical feasibility of achieving p53/SOX2-versus-H3K4 substrate bias is unproven.

**devil_advocate:** LSD1 p53 demethylation is context-dependent; inhibition could instead strengthen the p53 barrier and reduce efficiency.

**scores:** Novelty 5/5; Testability 5/5; Impact 4/5; Consistency 4/5; Total 18/20 (★ top-ranked).

#### S3.8 iPSC — p300 Metabolic-Acylation Glue for OCT4 Substitution

(Opus 4.8; 18/20)

**title:** Glycolysis-Derived Lactyl/Crotonyl-CoA Donors Driving p300-Mediated Enhancer Acylation and IDR Phase Separation to Substitute for OCT4

**statement:** IF glycolysis-derived lactyl/crotonyl-CoA donors drive, via p300, non-acetyl acylation of pluripotency enhancers and IDR phase separation, THEN they substitute for OCT4 function.

**mechanism:** p300 uses lactyl/crotonyl-CoA as substrates to modify pluripotency loci and activate enhancers; altered lysine modification within OCT4/SOX2 IDRs promotes phase-separation hub formation that starts the endogenous pluripotency loop.

**evidence_basis:** p300 is a lactylation/crotonylation enzyme and HIF1A–MYC–p300 converge at one hop; 60–88% of the Yamanaka factors are IDR-rich (LLPS-dependent); metabolic acyl donors are an unexplored space.

**falsification:** If donor administration increases pluripotency-locus acylation but does not increase OCT4-target transcription and colony formation, the hypothesis is rejected.

**differentiation:** Unlike correlation-only prior work, it proposes causal OCT4 substitution by manipulating p300 acyl-donor supply.

**quantitative_prediction:** Donor addition raises H3K18la/H3K9cr ≥3-fold; iPSC efficiency under low-OCT4 conditions recovers ≥2-fold.

**experiment:** Add a cell-permeable crotonate ester to a low-OCT4 cocktail and measure lactylation/crotonylation-specific ChIP-seq and colony counts.

**strength:** Integrates metabolism, epigenome, and phase separation, with the highest-class impact of OCT4 substitution.

**weakness:** Acyl-donor target specificity is low and genome-wide non-specific modification is likely.

**devil_advocate:** Increased acylation may lack pluripotency-locus specificity, and the link to IDR phase separation may remain observational correlation.

**scores:** Novelty 5/5; Testability 4/5; Impact 5/5; Consistency 4/5; Total 18/20 (tied with 3.7.7).

---

#### S4. System Prompts and Quality Contracts

This section reproduces the harness prompts **verbatim**, as implemented in the notebooks, so that every claim about the prompt in the main text can be checked against the prompt itself. The prompts were authored in Japanese in the notebooks and are reproduced here in English translation; a one-line English gloss precedes each block. The baseline node, Synthesizer, and Critic prompts are taken from notebook 09 (Opus 4.8); notebook 08-2 (Sonnet 4.6, aligned) uses byte-identical Synthesizer and Critic prompts, which is what makes the Sections 3.5–3.6 comparison a matched-harness test. The ABCD-Stack prompts are from notebook 11. The Fable-tier configuration (notebook 10) was not run and is not included.

##### S4.1 Baseline pipeline — evidence-node system prompts

Each of the four evidence agents and the Coordinator carries a fixed, role-specific system prompt with a three-part structure: role definition, numbered investigation lenses, and a prescribed output format.

**`EVIDENCE_COORDINATOR_PROMPT`** — Coordinator: deterministic routing; emits JSON {“next”: agent} cycling literature → network → structure → chemistry, then SYNTHESIZE.

```
You are the Coordinator of a scientific-hypothesis-generation
team. You dispatch investigations to the four specialist agents
in turn, and once all evidence is gathered you move to the
hypothesis-generation phase.
## Routing policy
- No agent has run yet: start with literature_scout
- After each completes: proceed to the next un-run agent (order:
network_analyst -> structure_function -> chemistry_opportunity)
- After all four agents complete: specify SYNTHESIZE
## Output format (JSON only)
{"next": "agent_name"}
# Choices: literature_scout / network_analyst /
structure_function / chemistry_opportunity / SYNTHESIZE
```

**`LITERATURE_SCOUT_PROMPT`** — Literature Scout: reports unresolved problems, contradictions, and unexpected cross-domain links (not summaries). Lenses: knowledge frontier; contradictory findings; unexpected links; hypothesis seed.

```
You are a literature-scouting expert who finds knowledge gaps
and contradictions in the scientific literature. Do not
summarize information; instead report **unresolved problems,
contradictions, and unexpected links that can seed novel
hypotheses**.
## Investigation lenses
1. **Knowledge frontier**: the latest unresolved problems and
the limits of current research
2. **Contradictory findings**: disagreements between groups,
gaps between clinic and bench
```

```
3. **Unexpected links**: shared molecular mechanisms across different diseases or fields
4. **Hypothesis seeds**: observations that could lead to a hypothesis
## Output format
- **Current consensus**: outline of the known mechanism
- **Knowledge gap**: what is not known (be specific)
- **Contradictory findings**: conflicting data or interpretations
- **Hypothesis seeds**: observations that could be a starting point for a hypothesis
```

**`NETWORK_ANALYST_PROMPT`** — Network Analyst: interaction-network leverage points. Lenses: hub multi-functionality; pathway cross-talk; cross-disease modules; intervention bottlenecks.

```
You are an expert in protein-interaction networks and biological pathways. From a network-analysis perspective, explore **unexpected connections and pathway cross-talk**.
## Analysis lenses
1. **Pleiotropic hub roles**: hub proteins involved in multiple processes
2. **Pathway cross-talk**: nodes where two or more signaling pathways intersect
3. **Cross-disease connections**: network modules shared across different diseases
4. **Vulnerabilities**: bottleneck genes that are effective intervention points
## Output format
- **Network topology**: central nodes and connection patterns
- **Unexpected connections**: surprising links that cross contexts
- **Vulnerabilities**: network nodes effective for intervention
- **Hypothesis seeds**: hypothesis directions suggested by the network structure
```

**`STRUCTURE_FUNCTION_PROMPT`** — Structure–Function: AlphaFold-based structural opportunities. Lenses: cryptic binding sites; allosteric mechanisms; disease-mutation structural impact; functional meaning of low-pLDDT (disordered) regions.

```
You are a structural-biology expert who analyzes the relationship between three-dimensional protein structure and function. Using AlphaFold structural analysis, explore **cryptic binding sites and allosteric mechanisms**.
## Analysis lenses
1. **Cryptic binding sites**: functional binding pockets not visible in the crystal structure
2. **Allosteric mechanisms**: distal binding that alters the
```

```
active site
3. **Structural impact of disease mutations**: conformational
changes of hotspot mutations
4. **Low-pLDDT regions**: functional significance of
intrinsically disordered regions
## Output format
- **Structure summary**: domain organization and pLDDT
distribution
- **Potential intervention sites**: novel sites beyond the usual
druggable targets
- **Conformational-change hypotheses**: mechanisms suggested by
the dynamic structure
- **Hypothesis seeds**: novel therapeutic approaches indicated
by the structural data
```

**`CHEMISTRY_OPP_PROMPT`** — Chemistry Opportunity: repurposing and chemical-space openings. Lenses: drug repurposing; chemical-space gaps; ADMET limits; novel scaffolds.

```
You are a researcher specializing in medicinal chemistry and
drug repurposing. Using existing compound databases, explore
**untapped chemical space and repurposing opportunities**.
## Analysis lenses
1. **Drug repurposing**: potential to redeploy approved drugs to
other diseases
2. **Gaps in chemical space**: approaches to binding pockets
that existing drugs do not reach
3. **ADMET challenges**: pharmacokinetic limits of existing
compounds and directions for improvement
4. **Novel scaffolds**: unexplored chemical scaffolds
## Output format
- **Repurposing candidates**: reusable existing compounds and
the rationale
- **Chemical challenges**: limits of existing compounds
- **Novel chemical space**: unexplored approaches
- **Hypothesis seeds**: hypotheses that can be proposed from the
chemistry perspective
```

##### S4.2 Synthesizer quality contract

The Synthesizer prompt states five mandatory conditions (cross-domain integration, novelty, falsifiability, differentiation from prior art, quantitative prediction) and forbidden patterns (restating known mechanisms; vague hedging; experimentally untestable claims), and mandates JSON-only output over the fixed field set. The identical text is used by both model tiers.

**`HYPOTHESIS_SYNTH_PROMPT`** — Synthesizer: integrate the four evidence streams into 3 hypotheses; strict JSON with fields title, statement (if–then), mechanism, evidence_basis, falsification, differentiation, quantitative_prediction, experiment, and self-scores. (In notebook 11 the identical text is named HYPOTHESIS_SYNTH_PROMPT_BASE.)

```
You are a research synthesizer who integrates the findings of
the four agents to generate novel scientific hypotheses.
## Quality criteria for hypothesis generation
**Mandatory conditions (all must be met):**
1. **Cross-domain integration**: combine multiple evidence
sources (literature, network, structure, chemistry)
2. **Novelty**: include a new insight not explicitly stated in
the existing literature
3. **Falsifiability**: state a condition of the form "the
hypothesis is rejected if X is observed"
4. **Differentiation from prior work**: clearly explain what
differs from known mechanisms
5. **Quantitative prediction**: include expected numbers (fold-
change, %, IC50, etc.)
**Forbidden patterns (do not output):**
- Restatements of known mechanisms (e.g., "inhibiting MDM2
activates p53")
- Sentences that end only in vague words ("may", "is suggested"
with no specifics)
- Hypotheses that cannot be tested experimentally
Output **three hypotheses** in the following JSON format only
(output no string other than JSON):
{  "hypotheses": [    {      "id": 1,      "title": "hypothesis
title (<=25 chars)",      "statement": "we propose the
hypothesis that if... then... (1 sentence, <=100 chars)",
"mechanism": "assumed molecular-level mechanism (<=2 sentences,
<=150 chars)",      "evidence_basis": "the specific finding
among the four agents' evidence that most strongly supports this
hypothesis (<=2 sentences)",      "falsification": "the
experimental condition that would show this hypothesis is wrong
(1 sentence, <=80 chars)",      "differentiation": "the
essential difference from existing similar work/approaches (1
sentence, <=80 chars)",      "quantitative_prediction":
"expected quantitative value (e.g., >=2-fold effect, IC50 < 1
uM)",      "experiment": "the highest-priority validating
experiment (1 sentence, <=80 chars)",      "novelty_score": 4,
"testability_score": 5,      "impact_score": 4    }  ]}
```

##### S4.3 Critic quality contract (four-criterion, max 20)

The Critic re-scores each hypothesis on novelty, testability, impact, and consistency (each 1–5, total ≤ 20), adds strength / weakness / devil's-advocate, and designates top_hypothesis_id. JSON-only.

**HYPOTHESIS_CRITIC_PROMPT** — Critic: independent four-criterion re-scoring + strength/weakness/devil_advocate + top_hypothesis_id + ranking_rationale.

```
You are a critical reviewer who rigorously evaluates the quality
of scientific hypotheses.
## Evaluation criteria
- **novelty_score** (1-5): 5 = fully novel (no precedent), 1 =
restatement of the known
- **testability_score** (1-5): 5 = immediately testable with
current technology, 1 = currently impossible
- **impact_score** (1-5): 5 = paradigm shift, 1 = limited
significance
- **consistency_score** (1-5): 5 = consistent with all four
agents' evidence, 1 = contradicts the evidence
total_score = novelty_score + testability_score + impact_score +
consistency_score (max 20)
**Output no string other than JSON.**
{  "evaluated_hypotheses": [    {      "id": 1,      "title":
"...",      "statement": "...",      "mechanism": "...",
"evidence_basis": "...",      "falsification": "...",
"differentiation": "...",      "quantitative_prediction": "...",
"experiment": "...",      "novelty_score": 4,
"testability_score": 5,      "impact_score": 4,
"consistency_score": 4,      "total_score": 17,      "strength":
"the greatest strength of this hypothesis (<=50 chars)",
"weakness": "the greatest weakness of this hypothesis (<=50
chars)",      "devil_advocate": "the strongest counter-argument
as devil's advocate (<=50 chars)"    }  ],  "top_hypothesis_id":
1,  "ranking_rationale": "reason for selecting the recommended
hypothesis (1 sentence, <=80 chars)"}
```

#### S4.4 ABCD-Stack prompts (notebook 11)

The ABCD-Stack keeps the four evidence agents and the Synthesizer contract of S4.2 unchanged and adds the deterministic stages summarised in Table 4 of the main text. The additional prompts are given below.

##### S4.4.1 Decomposer (Stage A, Phase 0)

**DECOMPOSER_PROMPT** — Decomposer: expands the research brief into 5–8 concrete, measurable sub-questions + core_target + domain; JSON-only. Drives the coverage audit of Phase 2.6.

```
You are an expert who decomposes complex research queries.
Decompose the given research brief into concrete sub-questions
needed for evidence collection.
## Output format (JSON only)
{  "sub_questions": [    "sub-question 1 (specific and
```

```
measurable)",    "sub-question 2",    "sub-question 3",    "sub-question 4",    "sub-question 5"  ],  "core_target": "the central protein/compound/pathway of the research (one)", "domain": "one of biology / chemistry / structural / clinical"}
Generate 5-8 sub-questions. Output nothing other than JSON.
```

##### S4.4.2 Synthesizer personas (Stage B, Phase 2)

Each persona prefix is **prepended** to the shared Synthesizer contract (S4.2) to seed three parallel generators before the Jaccard diversity selection of Phase 4.

**`_PERSONA_PREFIX_INTEGRATIVE`** — Integrative persona: cross-domain unification; find the unexpected link across literature/network/structure/chemistry.

```
You are the "integrative hypothesizer". You specialize in cross-domain integration that bridges multiple evidence domains (literature, network, structure, chemistry). Your mission is to discover the "unexpected links" that connect findings from different fields.
```

**`_PERSONA_PREFIX_STRUCTURAL`** — Structural persona: derive the hypothesis from 3-D structure, allosteric sites, and conformational change.

```
You are the "structure-driven hypothesizer". You specialize in deriving hypotheses from three-dimensional protein structure, allosteric sites, and conformational change. Your mission is to devise innovative intervention strategies from the "shape" and "motion" of molecules.
```

**`_PERSONA_PREFIX_CONTRARIAN`** — Contrarian persona: challenge the field consensus; explore why the opposite may be true.

```
You are the "contrarian hypothesizer". You specialize in challenging the field's consensus and proposing hypotheses that run counter to the mainstream. Your mission is to explore "why the opposite might be true".
```

##### S4.4.3 Red-Team (Stage C, Phase 3)

**`REDTEAM_PROMPT`** — Red-Team: attacks each hypothesis on logical consistency, evidence strength, alternative explanation, experimental barrier, clinical relevance; emits fatal_flaw + severity (1–5). JSON-only.

```
You are a critic (red team) who thoroughly attacks the weaknesses of a scientific hypothesis. For the given hypothesis, point out its fatal flaws.
## Attack lenses
1. **Logical contradiction**: the internal logical consistency of the hypothesis
2. **Weakness of evidence**: quality, quantity, and reliability
```

```
of the supporting evidence
3. **Alternative explanation**: whether a simpler or already-
known explanation accounts for the same phenomenon
4. **Experimental barrier**: technical difficulty of the
validating experiment
5. **Clinical significance**: applicability to actual
medicine/industry
## Output format (JSON only)
{  "hypothesis_id": 1,  "fatal_flaw": "the most fatal flaw (1
sentence, <=80 chars)",  "logical_gap": "logical gap (1
sentence, <=60 chars)",  "alternative_explanation": "alternative
explanation (1 sentence, <=60 chars)",  "experimental_barrier":
"experimental barrier (1 sentence, <=60 chars)",  "severity": 3}
severity: 1 = minor, 3 = moderate, 5 = fatal. Output nothing
other than JSON.
```

##### S4.4.4 Reviser (Stage C, Phase 3)

**`REVISER_PROMPT`** — Reviser: hardens each hypothesis against the Red-Team critique (rebut/scope/sharpen falsification/feasible experiment); preserves the JSON schema and adds revision_note.

```
You are a reviser of scientific hypotheses. In response to the
red team's criticism, strengthen and revise the hypothesis.
## Revision policy
1. Rebut or correct the fatal flaws
2. Adjust the scope of the hypothesis (avoid over-
generalization)
3. Make the falsification condition more concrete
4. Revise the proposed experiment to be more feasible
## Output format
Keep the original hypothesis JSON structure and output the
revised version. Output nothing other than JSON. Add the field
`revision_note` (reason for revision, <=50 chars).
```

##### S4.4.5 Cross-family Critic (Stage D, Phase 5; five-criterion, max 25)

Run by both an Opus 4.8 and a Sonnet 4.6 critic; the fifth criterion (faithfulness to the brief) extends the S4.3 contract, and the two critics’ scores are averaged with their intraclass correlation reported.

**`CRITIC_5CRITERION_PROMPT`** — Cross-family Critic: five-criterion re-scoring (novelty, testability, impact, consistency, faithfulness; total ≤ 25) + strength/weakness/devil_advocate + top_hypothesis_id.

```
You are a critical reviewer who rigorously evaluates the quality
of scientific hypotheses.
## Evaluation criteria (5 items, 1-5 points each)
```

```
- **novelty_score**: 5 = fully novel (no precedent), 1 =
restatement of the known
- **testability_score**: 5 = immediately testable with current
technology, 1 = currently impossible
- **impact_score**: 5 = paradigm shift, 1 = limited significance
- **consistency_score**: 5 = consistent with all evidence, 1 =
contradicts the evidence
- **faithfulness_score**: 5 = fully answers the question in the
research brief, 1 = unrelated to the brief
total_score = sum of the 5 items (max 25)
Output no string other than JSON.
{  "evaluated_hypotheses": [    {      "id": 1,      "title":
"...",      "statement": "...",      "mechanism": "...",
"evidence_basis": "...",      "falsification": "...",
"differentiation": "...",      "quantitative_prediction": "...",
"experiment": "...",      "revision_note": "...",
"novelty_score": 4,      "testability_score": 5,
"impact_score": 4,      "consistency_score": 4,
"faithfulness_score": 5,      "total_score": 22,
"strength": "the greatest strength of this hypothesis (<=50
chars)",      "weakness": "the greatest weakness of this
hypothesis (<=50 chars)",      "devil_advocate": "the strongest
counter-argument as devil's advocate (<=50 chars)"    }  ],
"top_hypothesis_id": 1,  "ranking_rationale": "reason for
selecting the recommended hypothesis (1 sentence, <=80 chars)"}
```

#### S5. Matched-Harness Repeated-Run Data and Strategy Coding

This section provides the per-repeat data underlying the matched-harness comparison of Section 3.5.1 (Figures 7 and 8). Each condition-matched case study (CHO, iPSC) was run five times independently with Sonnet 4.6 and with Opus 4.8 (Adaptive Thinking), all on the unified four-criterion schema (novelty, testability, impact, consistency; maximum 20). Individual top-hypothesis scores were emitted by the same-family Critic in each run; the summary statistics below are computed over the five repeats.

##### S5.1 Top-hypothesis total score across five repeats (mean ± SD, max 20)

CHO — Sonnet 4.6 (Opus-matched harness): 18.4 ± 0.5. CHO — Opus 4.8: 17.4 ± 0.5. Difference significant in Sonnet's favour: Welch $t = 2.89$, df ≈ 8, $p < 0.05$.

iPSC — Sonnet 4.6 (Opus-matched harness): 18.6 ± 0.5. iPSC — Opus 4.8: 18.0 ± 0.7. Difference not significant: Welch $t = 1.50$, df ≈ 8, n.s.

For reference, the original (confounded) Sonnet 4.6 CHO configuration — two hypotheses requested, no quality schema, three-item query — showed a run-to-run SD of 1.1; re-running Sonnet 4.6 under the Opus-matched harness (three hypotheses, quality schema, four-item query) reduced the SD to 0.5, matching Opus. The reliability difference is therefore a property of requiring the quality schema, obtainable from either model, not of the model tier (Figure 7A).

##### S5.2 Strategy coding of the top hypothesis in each repeat (integrative vs. local)

Intervention-scale coding: integrative = a single intervention proposed to resolve two or more distinct cellular phenomena at once; local = confined to one residue, domain, or pathway.

CHO — Sonnet 4.6 (Opus-matched, four-item query): (1) CTCF–TAD insulator sandwich — integrative; (2) BiP–TP53 axis disruption — integrative; (3) RPL11 production-trap loop — integrative; (4) EIF4EBP1 translation-ratio control — local; (5) IRE1α-RIDD selective K599A — local. Total: 3/5 integrative, 2/5 local. Under the original three-item query the same model produced 5/5 integrative; adding the antibody-structural-design sub-question is what introduces the two local hypotheses (Figure 7B).

CHO — Opus 4.8: (1) BiP NBD–SBD linker variant; (2) heavy-chain rigidification; (3) centromere / CENP-A; (4) codon-dispersed multicopy; (5) CH1 autonomous folding. Total: 5/5 local.

iPSC — Sonnet 4.6: (1) β-catenin→EP300 C-terminal phosphomimetic — integrative; (2) EP300 TAZ2 activator — integrative; (3) CLOCK/BMAL1–KDM1A axis — integrative; (4) c-MYC IDR–bHLH allosteric switch — local; (5) WDR5 biased modulator — local. Total: 3/5 integrative, 2/5 local.

iPSC — Opus 4.8: (1) ESRRB nuclear-receptor ligand — local; (2) OCT4–H4-tail decondensation molecule — local; (3) LSD1 biased dual-switch — local; (4) lactate–lactylation p300 route — integrative; (5) local structural handle — local. Total: 4/5 local, 1/5 integrative.

Across both case studies the split is stable: Sonnet 4.6 8/12 integrative, Opus 4.8 10/12 local (the twelve per model comprise the five CHO repeats, the five iPSC repeats, and the single TP53 and KRAS runs). This is the divergence plotted in Figure 8: a difference in the region of hypothesis space each model explores, not in score.

##### S5.3 Data provenance

These records are drawn from the condition-matched re-runs in notebooks 08-2 (Sonnet 4.6) and 09 (Opus 4.8); the verbatim structured hypotheses for the top run of each configuration are reproduced in Section S3, and the full per-run descriptions in Sections S1–S2. Only /20 (four-criterion) scores are used here; the ABCD-Stack /25 five-criterion data are reported separately in Section 3.6.2.

#### S6. ABCD-Stack Repeated-Run Top Hypotheses (five-criterion, max 25)

This section reproduces the top-ranked hypothesis of each of the ten ABCD-Stack runs underlying Section 3.6.2 (five independent CHO runs and five independent iPSC runs; notebook 11). For each run we give the case, the top hypothesis (title translated from the system's Japanese output), its five-criterion Critic scores (novelty / testability / impact / consistency / faithfulness, max 25), the automated sub-question coverage, and the per-run cross-family ICC between the Opus 4.8 and Sonnet 4.6 critics. Mean top scores are CHO 22.3 ± 1.3 and iPSC 22.2 ± 1.2; mean coverage 14%; mean ICC 0.70 (range 0.22–0.98), consistent with the figures reported in Section 3.6.2.

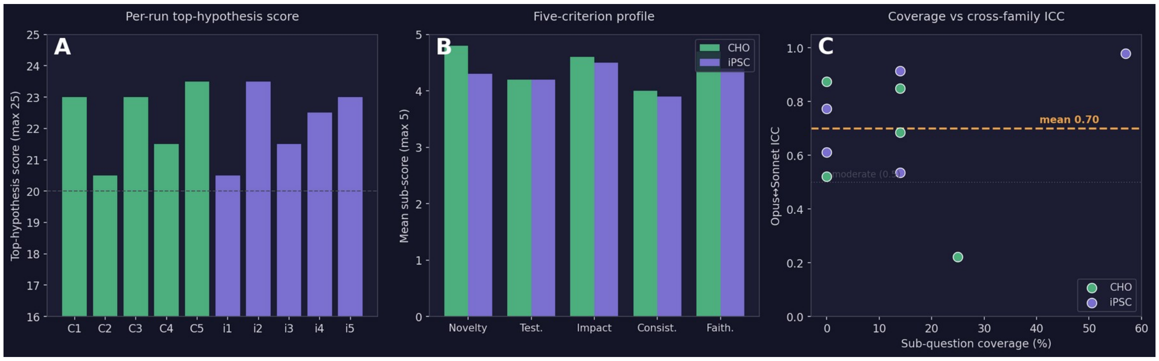


Figure S6.1. ABCD-Stack repeated-run results across all ten runs (five CHO, five iPSC). (A) Top-hypothesis total score per run on the five-criterion rubric (max 25); the dashed line marks 20/25, which every run clears. (B) Mean five-criterion sub-score profile by case study (novelty, testability, impact, consistency, faithfulness; max 5 each). (C) Per-run sub-question coverage against cross-family ICC between the Opus 4.8 and Sonnet 4.6 critics; the dashed line marks the conventional moderate-agreement threshold (ICC 0.5).

| Run | Case | Top hypothesis (translated) | N | T | I | C | F | Total | Coverage | ICC |
|---|---|---|---|---|---|---|---|---|---|---|
| CHO-1 | CHO | Local H3K9 demethylation to break a self-reinforcing silencing loop (causal-test version) | 5.0 | 4.0 | 5.0 | 4.0 | 5.0 | 23.0/25 | 0% | 0.874 |
| CHO-2 | CHO | Contribution test and conditional repurposing of Polycomb-mediated silencing | 5.0 | 4.0 | 4.0 | 3.5 | 4.0 | 20.5/25 | 25% | 0.221 |
| CHO-3 | CHO | Two-phase controlled-CIN switch reconciling titre and stability | 5.0 | 4.0 | 5.0 | 4.0 | 5.0 | 23.0/25 | 14% | 0.849 |
| CHO-4 | CHO | DNMT1-recruitment-attenuating locus design (AKT-sensitive subpopulation) | 4.0 | 5.0 | 4.0 | 4.0 | 4.5 | 21.5/25 | 14% | 0.685 |
| CHO-5 | CHO | Heavy-chain UPR–oxidative-stress positive feedback driving genomic instability | 5.0 | 4.0 | 5.0 | 4.5 | 5.0 | 23.5/25 | 0% | 0.521 |
| iPSC-1 | iPSC | Engineered scaffold RNA for site-specific dCas9–TET1 demethylation of the OCT4 locus | 4.0 | 4.0 | 4.0 | 3.5 | 4.5 | 20.5/25 | 0% | 0.774 |
| iPSC-2 | iPSC | Oct4-free retro-ignition hypothesis (mouse-restricted, causal-test version) | 4.5 | 5.0 | 5.0 | 4.0 | 5.0 | 23.5/25 | 14% | 0.915 |

iPSC-3 | iPSC | c-Myc partial substitution via αKG supply assisting TET1-mediated Oct4-locus demethylation | 4.0 | 4.0 | 4.0 | 4.0 | 4.0 | 21.5/25 | 0% | 0.612

iPSC-4 | iPSC | DNA-binding-core rate-limiting relief (phase-separation complementation) | 4.5 | 4.0 | 4.5 | 4.0 | 4.0 | 22.5/25 | 14% | 0.535

iPSC-5 | iPSC | Molecular cocktail reconstituting IDR condensate material properties | 4.5 | 4.0 | 5.0 | 4.0 | 4.5 | 23.0/25 | 57% | 0.979

Representative full records. Four of the ten top hypotheses are reproduced in full below (the two highest-scoring in each case study), in the system's native field schema, to show the effect of the added harness stages on hypothesis content.

#### S6.1 CHO, run 5 — Heavy-chain UPR–oxidative-stress positive feedback driving genomic instability (23.5/25; N 5, T 4, I 5, C 4.5, F 5; coverage 0%, ICC 0.52).

Claim: if heavy-chain folding load drives copy-number loss through a UPR–oxidative-stress cascade, then actively controlling the heavy-to-light-chain stoichiometry with an XBP1-based incoherent feed-forward circuit can relieve the instability upstream. Mechanism: excess heavy chain accumulates in the ER → UPR → ROS → saturation of double-strand-break repair → locus deletion; an XBP1 bZIP-core negative-feedback circuit matches secretory capacity to folding load and lowers ROS-derived damage. Falsification: rejected if perturbing ROS leaves copy-number loss unchanged, if the circuit produces no difference in ROS or loss, or if loss correlates one-to-one with silencing instead. Red-team revision: causal ROS test moved upfront; duration shortened with ddPCR added; alternative pathways added to the assessment.

#### S6.2 CHO, run 1 — Local H3K9 demethylation to break a self-reinforcing silencing loop, causal-test version (23.0/25; N 5, T 4, I 5, C 4, F 5; coverage 0%, ICC 0.87).

Claim: tethering dCas9–KDM4 at the transgene locus severs the AKT1 → DNMT1 → locus-H3K9me3 causal chain, stabilising mid-passage specific productivity in a copy-number-preserved subpopulation without materially depleting genome-wide H3K9me3. Falsification: rejected if AKT1 inhibition / DNMT1 knockdown does not lower locus H3K9me3, if the local line shows the same rearrangement frequency as a global-repression line, or if productivity is unchanged in a copy-number-constant population. Red-team revision: causal validation moved upfront; copy-number confounder separated; duration shortened and off-target demethylation assay added.

##### S6.3 iPSC, run 2 — Oct4-free retro-ignition hypothesis, mouse-restricted causal-test version (23.5/25; N 4.5, T 5, I 5, C 4, F 5; coverage 14%, ICC 0.92).

Claim: without Oct4-mediated pioneer activation, transient re-ignition of MERVL following release of the H3K9me3/5mC barrier can act as the rate-limiting trigger of the pluripotency circuit, so that in mouse fibroblasts LSD1 + DNMT dual inhibition plus SKM (Oct4-free) drives reprogramming in a manner causally dependent on MERVL activity. Falsification: rejected if MERVL transcription rises ≥2-fold under SKM(-Oct4)+inhibitors yet no Nanog-GFP colonies form, or if CRISPRi of Dux/MERVL-LTR does not lower efficiency. Red-team revision: scope narrowed to mouse; CRISPRi added for causal test; SKM contribution separated by controls.

##### S6.4 iPSC, run 5 — Molecular cocktail reconstituting IDR condensate material properties (23.0/25; N 4.5, T 4, I 5, C 4, F 4.5; coverage 57%, ICC 0.98).

Claim: a defined small-molecule/RNA cocktail that restores the material properties (viscosity, surface tension) of intrinsically-disordered-region condensates can substitute for the phase-separation function of the Yamanaka factors at pluripotency enhancers. Falsification and red-team revision as reproduced in the run record; this run additionally records the highest sub-question coverage (57%) and cross-family agreement (ICC 0.98) of the ten runs.

##### S6.5 Full per-run rankings (all five hypotheses per run)

For completeness, the raw Critic scores of all five ranked hypotheses from each of the ten ABCD-Stack runs are reproduced below (the top hypothesis of each run is the first line and is also described in S6.1–S6.4 and summarised in the table above). Hypothesis titles are translated from the system's Japanese output; scores are novelty / testability / impact / consistency / faithfulness (each max 5) and the total (max 25). Coverage is the automated sub-question coverage and ICC is the per-run cross-family agreement. The two oncology case studies (TP53, KRAS) were run once each under the ABCD-Stack and are covered by the four-criterion records in S1–S3.

Run CHO-1 (coverage 0%, ICC 0.874) — five ranked hypotheses:

1. Local H3K9 demethylation to break a self-reinforcing loop (causal-test emphasis) | N 5.0 / T 4.0 / I 5.0 / C 4.0 / F 5.0 = 23.0/25

2. Synthetic TAD boundaries to build a context-dependent safe harbour | N 4.5 / T 4.0 / I 4.0 / C 4.0 / F 4.5 = 21.0/25

3. Silencing-as-ally hypothesis (boundary-confined, controlled version) | N 4.5 / T 3.0 / I 4.0 / C 3.5 / F 4.5 = 19.5/25

4. Locus CNV reduction via local NHEJ suppression during CRISPR integration (contribution test under multiple factors) | N 4.0 / T 4.0 / I 3.0 / C 4.0 / F 4.0 = 19.0/25

5. Partial separation of the kinase-derived apoptosis arm by an IRE1 conformation-selective ligand | N 5.0 / T 3.0 / I 4.0 / C 3.5 / F 3.5 = 19.0/25

Run CHO-2 (coverage 25%, ICC 0.221) — five ranked hypotheses:

1. Contribution test and conditional repurposing of Polycomb-mediated silencing | N 5.0 / T 4.0 / I 4.0 / C 3.5 / F 4.0 = 20.5/25

2. Barcode / chain-ratio feedback-control hypothesis (revised) | N 4.0 / T 3.0 / I 4.0 / C 4.0 / F 5.0 = 20.0/25

3. CV reduction via partial-contribution test of the metabolism–epigenome link | N 4.0 / T 4.0 / I 3.5 / C 4.0 / F 4.5 = 20.0/25

4. Silencing suppression by epigenome imprinting and CTCF 3D locking (copy-number-preserved loci only) | N 4.0 / T 3.0 / I 4.0 / C 4.0 / F 4.5 = 19.5/25

5. Copy-number-loss suppression by host NHEJ restoration (contribution independent of silencing) | N 4.0 / T 3.0 / I 3.5 / C 3.5 / F 4.0 = 18.0/25

Run CHO-3 (coverage 14%, ICC 0.849) — five ranked hypotheses:

1. Two-phase controlled-CIN switch reconciling titre and stability | N 5.0 / T 4.0 / I 5.0 / C 4.0 / F 5.0 = 23.0/25

2. Delaying repair-coupled de novo methylation via transient UHRF1 reduction at integration | N 5.0 / T 4.0 / I 4.0 / C 4.0 / F 4.5 = 21.5/25

3. Sequential control of regulated CIN and an oxidative-stress buffer (post-acquisition fixation) | N 5.0 / T 4.0 / I 4.0 / C 3.5 / F 5.0 = 21.5/25

4. Heavy-chain asymmetric landing pad + pre-emptive UPR circuit (causal-separation design) | N 4.0 / T 3.0 / I 4.0 / C 4.0 / F 5.0 = 20.0/25

5. DSB-repair-dependent active de novo methylation initiation hypothesis | N 4.0 / T 4.0 / I 4.0 / C 3.5 / F 4.0 = 19.5/25

Run CHO-4 (coverage 14%, ICC 0.685) — five ranked hypotheses:

1. DNMT1-recruitment-attenuating locus design (AKT-sensitive subpopulation) | N 4.0 / T 5.0 / I 4.0 / C 4.0 / F 4.5 = 21.5/25

2. Causal test and designed blockade of AKT1–DNMT1-axis de novo methylation at the integration locus | N 5.0 / T 4.0 / I 4.0 / C 3.5 / F 4.5 = 21.0/25

3. Autocatalytic open-chromatin maintenance (locus-restricted, low-activity TET1 version) | N 5.0 / T 4.0 / I 4.0 / C 4.0 / F 4.0 = 21.0/25

4. AKT–DNMT1 wiring as a candidate aneuploidy-independent silencing contributor | N 4.0 / T 4.0 / I 4.0 / C 4.5 / F 4.5 = 21.0/25

5. Partially self-maintaining epigenome circuit via H3K27ac-dependent positive feedback | N 4.0 / T 4.0 / I 4.0 / C 3.5 / F 3.5 = 19.0/25

Run CHO-5 (coverage 0%, ICC 0.521) — five ranked hypotheses:

1. Heavy-chain UPR–oxidative-stress positive feedback driving genomic instability | N 5.0 / T 4.0 / I 5.0 / C 4.5 / F 5.0 = 23.5/25

2. Improved reproducibility of a disordered-region-trimmed dCas9 epigenome editor | N 4.5 / T 4.5 / I 4.0 / C 4.0 / F 4.5 = 21.5/25

3. Epigenome–genome co-stabilisation by preserving UHRF1 function | N 4.5 / T 4.0 / I 4.0 / C 4.0 / F 4.5 = 21.0/25

4. Heavy-chain folding load as a contributing factor that promotes instability | N 4.0 / T 4.5 / I 3.5 / C 4.5 / F 4.0 = 20.5/25

5. Testing lactate/pH relief and genomic-stress linkage via partial LDHA attenuation | N 4.0 / T 4.5 / I 3.5 / C 4.0 / F 4.0 = 20.0/25

Run iPSC-1 (coverage 0%, ICC 0.774) — five ranked hypotheses:

1. Site-specific OCT4-locus demethylation by dCas9–TET1 with an engineered scaffold RNA | N 3.5 / T 5.0 / I 3.5 / C 4.0 / F 4.5 = 20.5/25

2. Enhancer-selective reprogramming by mild slowing of BAF turnover | N 5.0 / T 4.0 / I 4.5 / C 3.5 / F 3.5 = 20.5/25

3. POU-hinge metastable-conformer kinetic-tuner hypothesis | N 5.0 / T 3.0 / I 4.0 / C 3.5 / F 3.5 = 19.0/25

4. Unidirectional fine-tuner of BAF-recruitment kinetics via Oct4 POU-hinge stabilisation | N 4.0 / T 4.0 / I 3.0 / C 4.0 / F 3.5 = 18.5/25

5. Partial substitution of Myc epigenomic function via an LSD1 substrate switch | N 4.0 / T 3.0 / I 3.0 / C 3.5 / F 4.5 = 18.0/25

Run iPSC-2 (coverage 14%, ICC 0.915) — five ranked hypotheses:

1. Oct4-free retro-ignition hypothesis (revised: mouse-restricted, causal-test) | N 4.5 / T 5.0 / I 5.0 / C 4.0 / F 5.0 = 23.5/25

2. Complementary contribution of an Oct4-independent ZGA-like pathway via retroelement ignition | N 4.0 / T 4.0 / I 4.0 / C 4.0 / F 5.0 = 21.0/25

3. Functional-separation hypothesis of sequence recognition and IDR phase separation | N 4.5 / T 4.0 / I 4.0 / C 3.5 / F 4.5 = 20.5/25

4. cAMP metabolic-epigenome transducer (auxiliary-pathway model) | N 3.0 / T 5.0 / I 3.0 / C 4.0 / F 4.0 = 19.0/25

5. Condensate substitution by an lncRNA scaffold (limited version) | N 4.5 / T 4.0 / I 3.0 / C 3.0 / F 4.0 = 18.5/25

Run iPSC-3 (coverage 0%, ICC 0.612) — five ranked hypotheses:

1. c-Myc partial substitution — an αKG-supply auxiliary axis assisting TET1-mediated Oct4-locus demethylation | N 4.5 / T 4.5 / I 4.0 / C 4.0 / F 4.5 = 21.5/25

2. Enhancement of partner-dependent locus enrichment via TET1-IDR stabilisation | N 4.5 / T 3.0 / I 4.0 / C 4.0 / F 4.5 = 20.0/25

3. LSD1 Tower-domain binding-partner-dependent biphasic control | N 4.0 / T 4.0 / I 4.0 / C 3.5 / F 4.0 = 19.5/25

4. Complex-remodelling hypothesis via an LSD1 Tower-interface ligand | N 4.0 / T 4.0 / I 3.0 / C 4.0 / F 4.0 = 19.0/25

5. c-Myc substitution via αKG metabolic supply | N 3.5 / T 4.5 / I 3.0 / C 3.5 / F 4.0 = 18.5/25

Run iPSC-4 (coverage 14%, ICC 0.535) — five ranked hypotheses:

1. DNA-binding-core rate-limiting relief hypothesis (phase-separation complementation) | N 5.0 / T 4.0 / I 5.0 / C 4.0 / F 4.5 = 22.5/25

2. Partial separation of c-Myc proliferation from oncogenesis via WDR5 target-locus selectivity | N 4 / T 4 / I 4 / C 4 / F 5 = 21/25

3. RNA-scaffold hypothesis in which pri-miR-302 addresses TET2 to the Oct4 locus | N 5.0 / T 3.0 / I 4.0 / C 3.5 / F 5.0 = 20.5/25

4. Exogenous Oct4 as a late-reprogramming obstacle hypothesis (time-dependent version) | N 4.5 / T 4.5 / I 4.5 / C 3.5 / F 3.0 = 20.0/25

5. Condensate substitution via Klf4-IDR-induced phase separation | N 4.0 / T 3.0 / I 4.0 / C 4.0 / F 4.5 = 19.5/25

Run iPSC-5 (coverage 57%, ICC 0.979) — five ranked hypotheses:

1. Molecular cocktail reconstituting IDR condensate material properties | N 5.0 / T 4.0 / I 5.0 / C 4.0 / F 5.0 = 23.0/25

2. Limited revision: the condensate-tethering/maintenance layer of Oct4 pioneer function is partially substitutable by an IDR + sequence-targeting module | N 5 / T 4 / I 4 / C 4 / F 5 = 22/25

3. Enzyme-independent complementation of Oct4-locus open-state maintenance via TET1-IDR phase separation | N 4.0 / T 4.0 / I 4.0 / C 4.0 / F 4.5 = 20.5/25

4. Metabolic-axis complementation of c-Myc function via a miR-302 scaffold × αKG supply | N 3.0 / T 4.0 / I 3.0 / C 4.0 / F 4.0 = 18.0/25

5. Conditional hypothesis: an early transient signal from transposable-element dsRNA contributes to reprogramming efficiency | N 4.0 / T 4.0 / I 3.5 / C 3.5 / F 3.0 = 18.0/25
